# Super-resolution fight club: A broad assessment of 2D & 3D single-molecule localization microscopy software

**DOI:** 10.1101/362517

**Authors:** Daniel Sage, Thanh-An Pham, Hazen Babcock, Tomas Lukes, Thomas Pengo, Jerry Chao, Ramraj Velmurugan, Alex Herbert, Anurag Agrawal, Silvia Colabrese, Ann Wheeler, Anna Archetti, Bernd Rieger, Raimund Ober, Guy M. Hagen, Jean-Baptiste Sibarita, Jonas Ries, Ricardo Henriques, Michael Unser, Seamus Holden

## Abstract

With the widespread uptake of 2D and 3D single molecule localization microscopy, a large set of different data analysis packages have been developed to generate super-resolution images. To guide researchers on the optimal analytical software for their experiments, we have designed, in a large community effort, a competition to extensively characterise and rank these options. We generated realistic simulated datasets for popular imaging modalities – 2D, astigmatic 3D, biplane 3D, and double helix 3D – and evaluated 36 participant packages against these data. This provides the first broad assessment of 3D single molecule localization microscopy software, provides a holistic view of how the latest 2D and 3D single molecule localization software perform in realistic conditions, and ultimately provides insight into the current limits of the field.

## INTRODUCTION

Image processing software is central to single molecule localization microscopy (SMLM), which delivers an order of magnitude resolution improvement on diffraction limited conventional fluorescence microscopy, from 250 nm to approximately 20 nm resolution, by temporal separation of fluorophores within a sample^1–3^. Efficient and automated image processing is essential to extract the super-resolved positions of individual molecules from thousands of raw microscope images, containing millions of blinking fluorescent spots.

Improvements in SMLM image processing algorithms have been crucial in maximizing spatial resolution and in reducing the imaging time of SMLM for compatibly with live cell imaging^4–6^. If SMLM is to achieve a resolving power approaching that of electron microscopy, the analysis software employed needs to be robust, accurate, and performing at current algorithmic limits. This can only be achieved through rigorous quantification of SMLM software performance.

The first localization microscopy software challenge was carried out in 2013, to enable robust benchmarking of 2D localization microscopy software packages^7^. But biology is not just a 2D problem, and a key focus of localization microscopy is the imaging of 3D imaging of nanoscale cellular processes^8,9^. 3D localization microscopy is a more difficult image processing problem than 2D SMLM. In addition to finding the center of diffraction limited spots to super-resolve lateral position, 3D SMLM algorithms must also extract axial information from the image, usually by measuring small changes in the shape of a point-spread function^10^ (PSF).

There are roughly three common approaches for 3D SMLM. First, point spread function engineering, where the axial asymmetry of the microscope point spread function (PSF) is increased by introducing intentional aberrations in the system, ranging from simple astigmatism^10^ to more complex PSF manipulation such as the double helix PSF method^11^. Second, biplane or multiplane imaging, where axial position is measured based on simultaneous measurement of PSF shape at two or more focal planes^12^. Third, dual objective based interferometry, where Z-position is calculated from single photon interference between opposing objectives^13^. Multiplane and PSF engineering methods typically obtain axial resolutions on the order of 50 nm^10,11^. Interferometry achieves the best axial resolution, 10-20 nm^13^, but is not yet widely adopted.

Despite the widespread use of 3D localization microscopy, and challenging nature of 3D SMLM image processing, the performance of software for 3D single molecule localization microscopy has previously only been assessed for 2 or 3 software packages at a time, and without standard test data or metrics^14–17^. In the absence of common reference datasets and reliable assessment procedure of 3D software performance, it is not possible to objectively assess how different software affects final image quality, or which algorithmic approaches are most successful. Crucially, end-users cannot determine which 3D SMLM software package and imaging modality is optimal for their application.

We therefore ran the first 3D localization microscopy software challenge, to assess the performance of 3D SMLM software. We assessed software performance on synthetic datasets for three popular 3D SMLM modalities: astigmatic imaging, biplane imaging and double helix point spread function microscopy. We also assessed astigmatism software performance on two real STORM datasets. We ran a second 2D localization microscopy software challenge, to reassess the 2D SMLM software state-of-the art on new, tougher, more realistic datasets.

Our simulations incorporate experimentally acquired point spread functions for maximal authenticity, used signal and noise levels based closely on common experimental conditions, and incorporated a realistic 4-state model of fluorophore photophysics^18^. Our synthetic data was designed to mimic two common classes of cellular structure: narrow line-like microtubules (MT) and larger tubes similar to the endoplasmic reticulum (ER) or mitochondria. Our simulations also included conditions with low density (LD) of active fluorophores, used experimentally to obtain maximal resolution, and with high density (HD) of active fluorophores, used experimentally for fast or live cell imaging.

## RESULTS

### Competition design

We established a large committee from within the SMLM research community, including experimentalists and software developers, to define the scope of the challenge, ensure realism of the datasets and define analysis metrics. We further opened this discussion to the whole community, through an open forum, discussing best practices for the implementation of this contest^19^.

Thirty-six software packages have been entered in the competition thus far, including four packages used in commercial software (Table S1, Supplementary Note 5). Excitingly, participation in the competition actually led at least 8 teams to their software to support additional 3D SMLM modalities, showing how competition fosters microscopy software development.

In 2016, we ran a first round of the 3D SMLM competition with explicit submission deadlines, with 30 competitor teams, culmination in a special session at the 6th annual Single Molecule Localization Microscopy Symposium (SMLMS 2016). Since then, the challenge has been opened to continuously accept new entries. We have had 12 new registrations of which 5 have submitted localizations, including a multiple best-in-class performer (SMAP-2018^20^, an updated version of previously entered software) demonstrating the utility of the competition as an evolving measure of the state of the field.

### Realistic 3D simulations

Testing super-resolution software on experimental data lacks the ground truth information required for rigorous quantification of software performance. Therefore, realistic simulated 3D SMLM datasets are required. After comparison of simulated microscope PSFs with multiple experimental PSFs from SMLM microscopes around the world, we observed that a critical challenge to realistic 3D SMLM simulations was to accurately model the experimental microscope PSF for each 3D modality. Even experimental 2D PSFs showed significant aberrations away from the focal plane (Fig S10).

3D SMLM inherently involves addition of aberrations to the microscope PSF to encode the Z-position of the molecule. For the PSF models included in the competition: 2D, astigmatic (AS), double helix (DH), and biplane (BP), we observed that the PSFs showed complex aberrations not well described by simple analytical models (Fig S10). We thus combined experimental 3D PSFs with simulated ground truth by performing simulations using PSFs directly derived from experimental calibration data (Fig 1, *Methods*). The experimental PSFs used to generate the simulated data are available online (*Methods*). As the goal of this study was to compare software obtained on typical SMLM microscopes, we deliberately chose PSFs representative of common implementations of each 3D modality. However, additional PSF engineering should improve results of any specific modality, for example adaptive-optics corrected astigmatism^21^, or reduced Z-range, higher SNR DH-PSF designs^22^.

**Figure 1:**
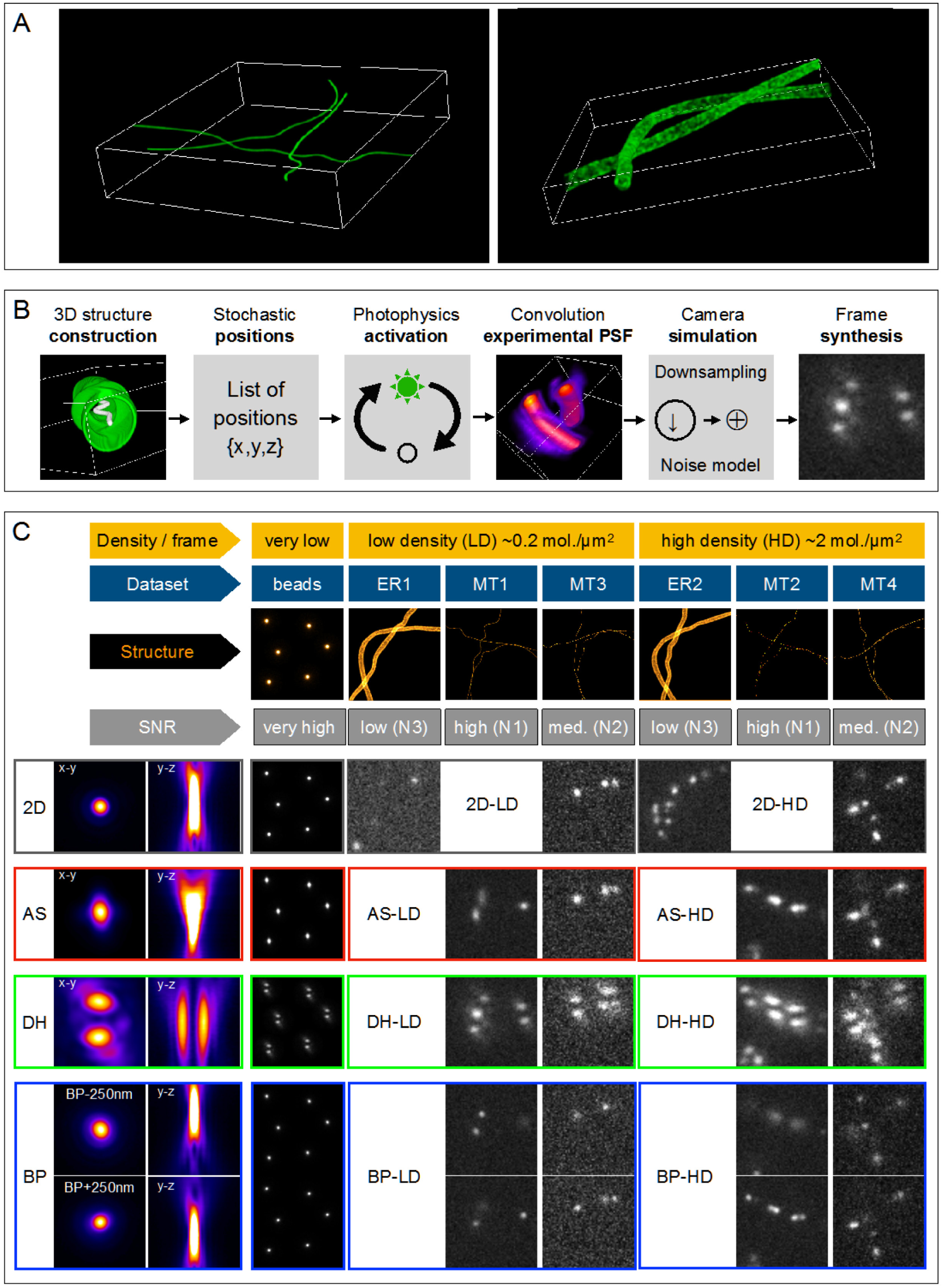
Summary of SMLM challenge simulations. **A**. 3D rendering of microtubules and endoplasmic reticulum samples in a 6.4 *μ*m × 6.4 *μ*m x 1.5 *μ*m volume. **B**. *Key simulation steps*. The structure is constructed from 3D tubes continuously defined by three B-spline functions in the volume of interest. Membranes of the tubes are densely populated with possible positions. Fluorophores follow a 4-state photophysics model. Activations of a given frame are convolved with the experimental PSF and shot & camera noise is added. **C**. Summary of all 16 challenge datasets, calibration data and experimental PSFs. Each dataset is characterized by its structure (endoplasmic reticulum (ER) or microtubules (MT)), by it modality (2D, AS, DH, BP), its density (LD or HD) and by its SNR determined by the level of noise N1, N2, and N3. Left column: orthogonal projections of the experimentally-derived PSF. Eight categories were proposed for the challenge containing two datasets each, 2D-LD and 2D-HD, grey; AS-LD and AS-HD, red, DH-LD and DH-HD, green; BP-LD and BP-HD, blue.

For the 3D competition, we simulated synthetic 25 nm diameter microtubules (Fig 1). For the 2D competition, in addition to synthetic microtubules (MT), we simulated larger diameter 150 nm cylinders, designed to approximate larger cellular structures such as mitochondria and the endoplasmic reticulum (ER) (Fig 1). We incorporated a 4-state model of fluorophore photophysics, including a transient dark state (dye “blinking”) and a bleaching pathway (Fig S1C).

As performance at different density of active emitters is a key challenge for SMLM software, we generated 3D competition datasets at both sparse emitter density (0.2 mol. [molecule] μm^−2^) and high emitter density (2 mol. μm^−2^). We additionally generated a very high density dataset (5 mol. μm^−2^) for the 2D competition.

We generated data at three different signal-to-noise ratio (SNR) levels, based on real signal to noise levels encountered under common SMLM experimental scenarios: fixed cells antibody labelled with organic dye^10^, fluorescent protein labelling^1^, and live cell affinity dye labelling^23,24^.

Together, these simulations closely resemble experimental 3D and 2D data under a range of challenging conditions of SNR, spot density, axial thickness and structure summarized in Table S2. In addition, we provide simulated z-stacks of bright beads for software calibration. The competition datasets are available online (*Methods*).

### Quantitative performance assessment of 3D software

We assessed software performance by 26 quality metrics (*Supplementary Note 1*). The complete set of summary statistics, axially resolved performance and super-resolved images is available for each competition software on the competition website. We built an interactive ranking and graphing interface that allows easy ranking and graphing of software performance by any metric, including new user defined metrics (Fig S11). Detailed individual software reports can also be accessed, along with a tool for side-by-side comparison of software (Fig S11, S22).

We focused our analysis primarily on metrics directly derived from single molecule localizations. Choice of ranking metric is discussed in detail in Supplementary Note 1.6, where several alternative ranking metrics are also presented.

#### 1. Single molecule localization error

The foremost consideration for localization software is how accurately it finds the position of labelled molecules. This was quantified as the root mean squared localization error (RMSE) between the measured molecule position and the ground truth, in both the lateral (XY) and axial (Z) dimensions.

#### 2. Ability to successfully detect fluorescent molecules

In addition to localization precision, SMLM image resolution depends critically on number of localized molecules^25^, so it is crucial for SMLM software to accurately detect a large fraction of molecules in a dataset, and minimize false localizations. For every frame, we identified the localizations that are close enough to a ground-truth position as true-positives (TP), the spurious localizations as false-positives (FP) and the undetected molecules as false-negatives (FN). We then computed the *Jaccard index* (JAC, %), which measures the fraction of correctly detected molecules in a dataset:

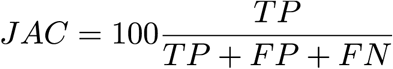

For ranking purposes, we developed a single summary statistic for overall evaluation of software performance, which we term the *efficiency (E)*, encapsulating both the software’s ability to find molecules, measured by the Jaccard index, and the software’s ability to precisely localize molecules.

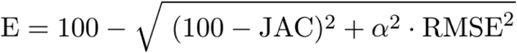

The trade-off between these two metrics is controlled by a parameter α. In a retrospective analysis, we chose α = 1 nm^−1^ for the lateral efficiency E_lat_, α = 0.5 nm^−1^ for the axial efficiency E_ax_, based on the linear regression slope between the localization errors and Jaccard index (Fig 17J-K). Using this definition, an average software performance has an efficiency in the range 25-75, a perfect software would have the maximum efficiency of 100. Overall 3D efficiency was calculated as the average of lateral and axial efficiencies. Overall software rankings (Fig 2) were calculated as the sum of rankings for high and low SNR datasets.

**Figure 2:**
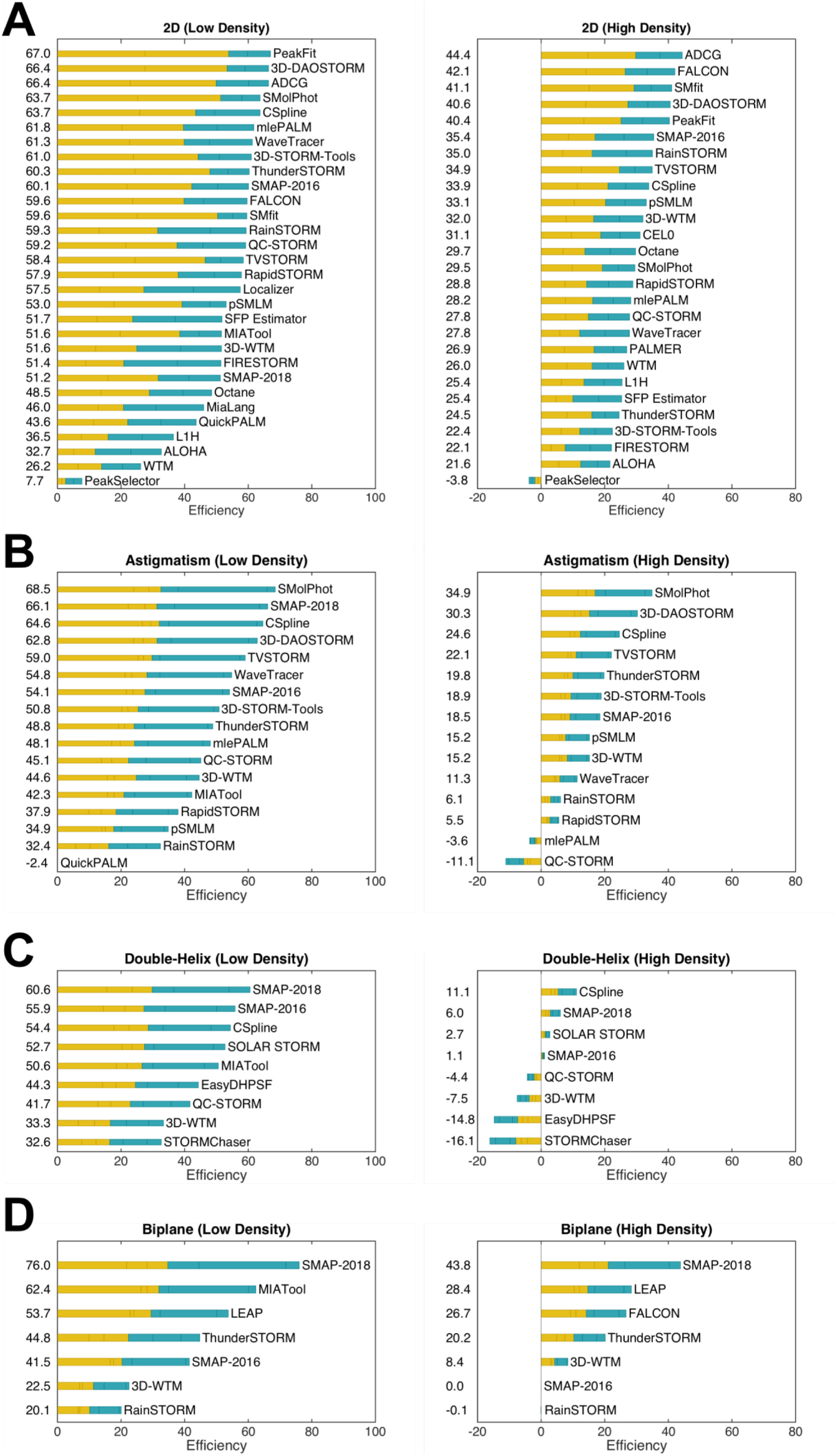
Leaderboards for each competition modality, at low and high spot density. Ranking is based on the efficiency of software based on fraction of successfully detected molecules (Jaccard index) and precision of localization (RMSE, root mean square error, lateral & axial). The contribution of the high SNR dataset is plotted in orange and the contribution of the low SNR dataset to the efficiency is plotted in blue.

### Performance of 3D software

Complete rankings for each imaging modality and spot density are presented (Fig 2), together with summary information on all competition software (Table S1, *Supplementary Note 1*). As these data are continuously updated on the competition website, this resource provides microscopists with a quick reference for the current state of the art, including current best-in-class performers for each category.

After assembling an overall summary of best performers for each competition category, we investigated the performance of software within each imaging modality.

#### Astigmatic localization microscopy

Astigmatic localization microscopy is probably the most popular imaging 3D SMLM modality, reflected by the highest number of software submissions in the 3D competition (Fig 2). For astigmatism, we observed a large spread of software performance, even for the most straightforward high SNR, low spot density (LD) conditions (Fig 3, Table S5). The best-in-class software (SMAP-2018) has significantly better localization error and Jaccard index performance than average (lateral RMSE 26 nm best vs 38 nm average, axial RMSE 29 nm best vs 66 nm average, Jaccard index 85 % best vs 74 % average). Clearly, the quality of the image reconstruction depends strongly on choice of 3D software.

**Figure 3:**
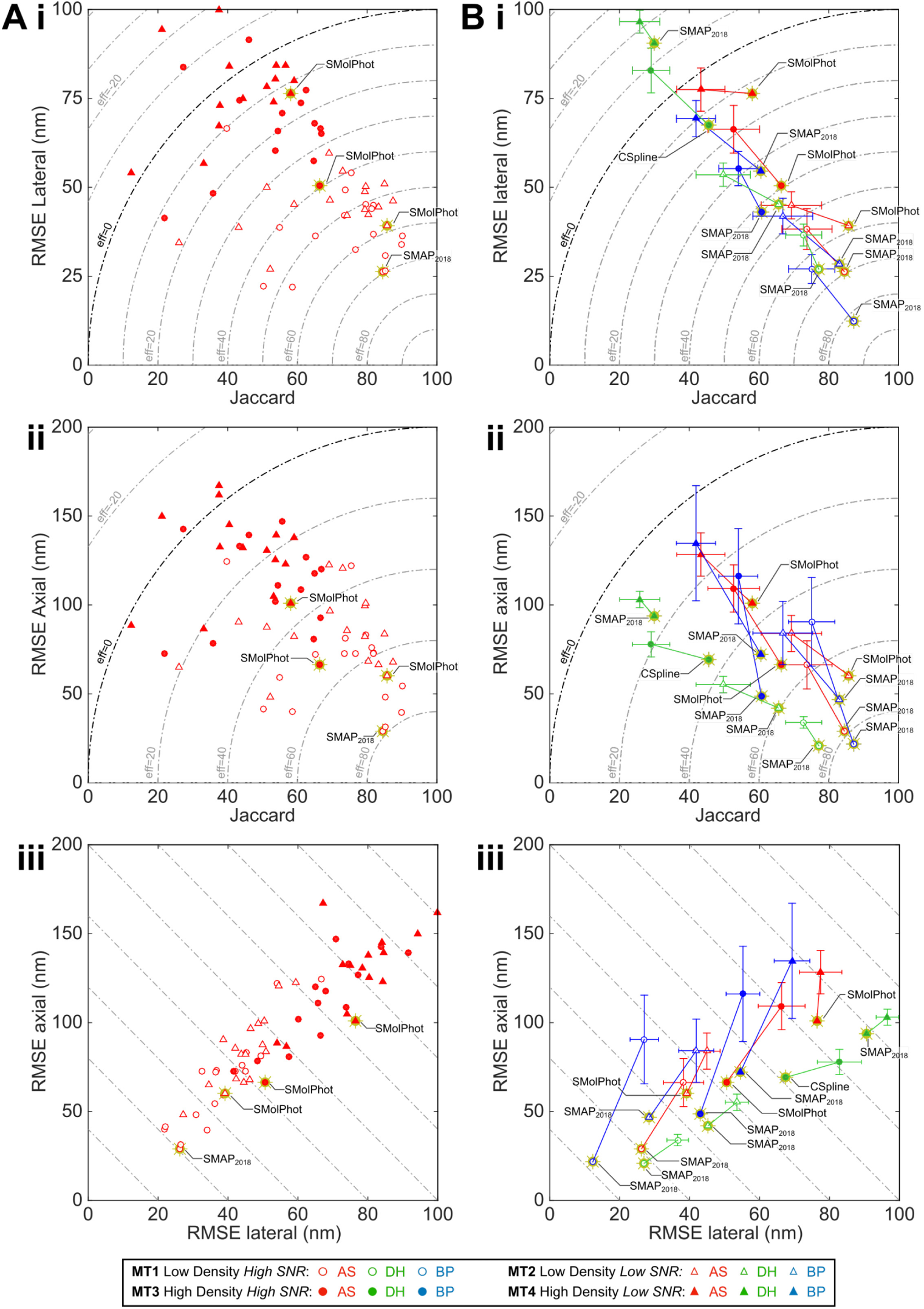
Comparison of 3D software performance. Gold stars indicate top performers for each dataset. Dashed lines in top, middle panels indicate overall efficiency (higher is better). ***A***. Localization error and spot detection performance of all astigmatic SMLM software. ***B***. Average (colored marker with *s.d*. error bars) and best-in-class (colored marker with gold star) software performance for all competition modalities. *AS, astigmatism; DH, double helix; BP, biplane*.

To investigate the reasons for software variation, we inspected plots of software performance as a function of axial position in the low density, high SNR dataset for best-in-class and representative middle-range software (Fig S7A). We observed that the key cause of the spread in software performance is variation in software performance away from the focal plane. Near the focal plane, most software packages perform well. However, the axial and lateral RMSE away from the plane of focus is significantly higher for the best in class software, and the Jaccard index is also slightly improved (Fig S7A). This is also visibly apparent in the super-resolved images (Fig 4A). We observed that best-in-class software had a Z-range (the FWHM range of axially resolved software recall, *Methods*) of 1170 nm, greater than two-thirds of the simulated range. Outside this range, the recall and Jaccard index dropped sharply, probably due the large increase in PSF size and decrease in effective SNR at significant defocus (Fig S10).

**Figure 4:**
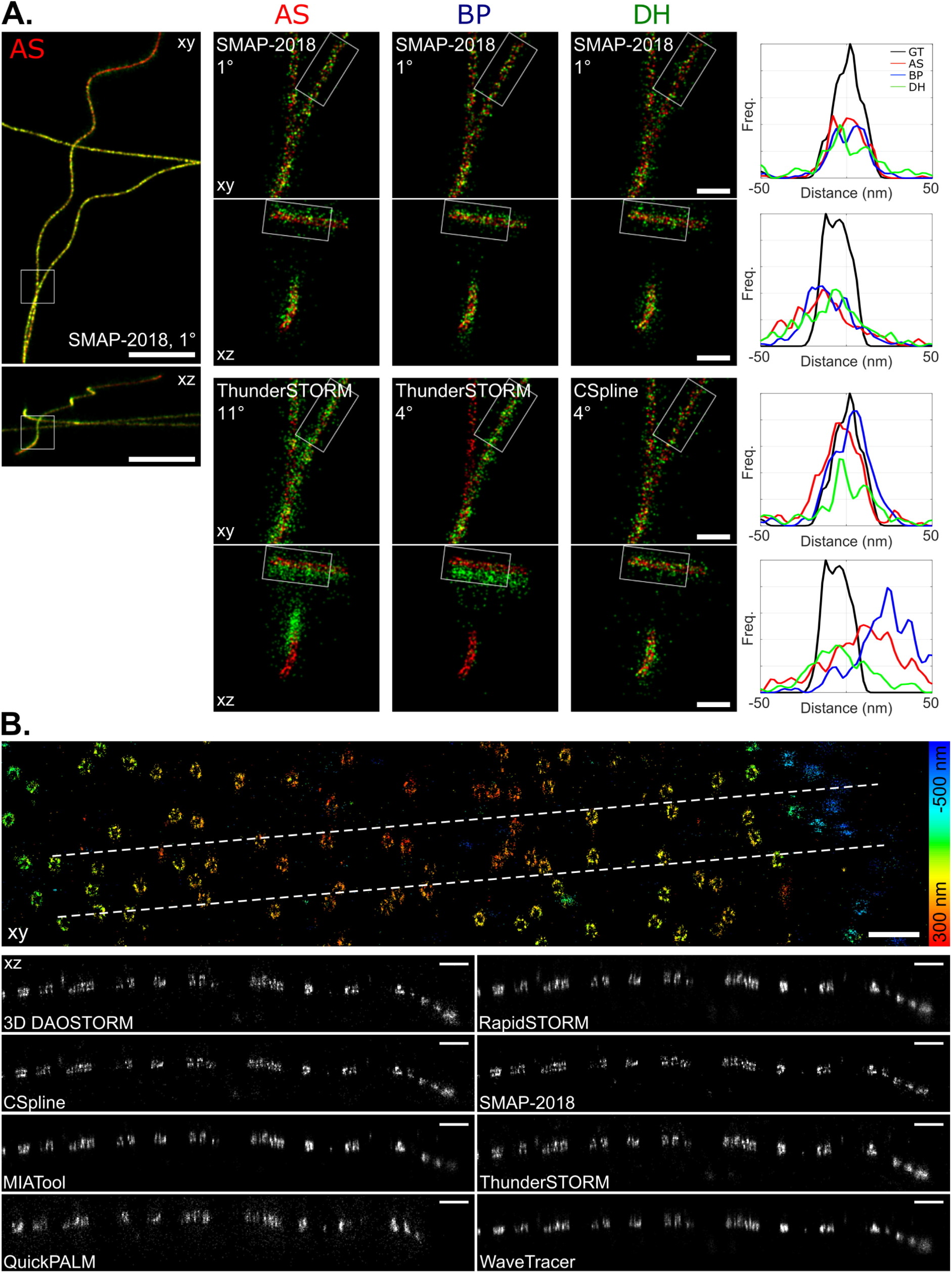
**A**. *Super-resolved xy and xz projection images of 3D competition datasets for best-in-class (top) and representative average (bottom) software in each modality, for high SNR low density dataset*. *Left*: *xy* and *xz* overview images for winning AS software. *Middle*: *xy* and *xz* zoom images of boxed regions in left panel, for winning and mid-range software, each modality. *Right*: *Xy* and *xz* line profiles of winning and mid-range software for each modality, for boxed regions in middle panel. *Image colors*: red, ground truth; green, software results. *Line profiles*: GT, ground truth, black; AS, astigmatism, red; BP, biplane, blue; DH, double helix, green. *Panel key:* Software-name Dataset-Ranking°. *Scale bar*: full image, 1 μm, magnified regions, 100 nm. ***B***. *Performance of astigmatism software on real 3D STORM images of the nuclear pore complex. Top:* Super-resolved overview image in *Xy* for 3D-DAOSTORM software, color coded for depth. *Bottom: Xz* orthoslices along 600 nm wide dashed region indicated in top panel for 8 astigmatism software packages. The best performing software clearly resolves the top and bottom of the NPCs. *Scale bar*s, 500 nm.

When we examined results for the low SNR, low density dataset (Fig 2B, 3B), we found an expected two-fold degradation in best-in-class RMSE (lateral RMSE 39 nm, axial RMSE 60 nm), due to the decrease in image SNR. However, the best-in-class software (SMolPhot^26^) Jaccard index was effectively constant between the low and high SNR datasets (86 % vs 85 %), although the Z-range did drop at lower SNR (930 nm vs 1120 nm). The best astigmatism software packages were thus remarkably good at finding spots at low SNR, even away from the plane of focus.

We analyzed how close software performance was to theoretical limits by calculating the Cramér-Rao lower bound (CRLB) as a function of axial position for each dataset and comparing it to the best-in-class software results (Fig S8, S9, Supplementary Note 4). Close to the focus, best-in-class software was close to CRLB performance (within 25 %), but significant deviations for the CRLB limit occurred > 200 nm. This could be due to the difficulty in actually detecting the spots away from focus.

When we examined astigmatic software performance for the challenging high spot density datasets (Fig 2B, 3), performance was reduced. For the high SNR high spot density dataset (best software, SMolPhot), localization error increased and Jaccard index decreased significantly compared to the low density condition (lateral RMSE best HD 51 nm vs best LD 27 nm, axial RMSE best HD 66 nm vs best LD 29 nm, Jaccard index best HD 66 % vs best LD 85 %). Inspection of the super-resolved images (Fig S3) nevertheless shows acceptable results for the HD dataset, particularly in the lateral dimension. In many circumstances, the performance reduction at 10x higher spot density should be acceptable for 10x faster, potentially live-cell-compatible, imaging speed. We also observed a large spread of software performance for the high density datasets, probably because a significant fraction of the software packages were primarily designed for low density conditions.

We observed poor performance for the most challenging low SNR high spot density astigmatism dataset (Fig 2, 3, S4, best software SMolPhot). Best-in-class localization precision and Jaccard index decreased significantly (lateral RMSE 76 nm, axial RMSE 101 nm, Jaccard index 58 %). These data suggest that low SNR high density 3D astigmatic localization microscopy entails a significant reduction in image resolution.

#### Double helix point spread function localization microscopy

We next analyzed the performance of the double helix software (Fig 3B, S14A). For the software in the high SNR low spot density condition, double helix software showed more uniform performance than astigmatism. Best-in-class software (SMAP-2018) showed only a limited improvement compared with average software (Fig 3B, lateral RMSE, 27 nm best vs 37 nm average; axial RMSE 21 nm best vs 34 nm average; Jaccard index 77 % best vs 73 % average). In general software localization performance was close to the CRLB (Fig S8, S9). We observed that performance of the software away from the focal plane is relatively uniform (Fig 4A, S7A), and best-in-class Z-range at high SNR was large at 1180 nm (Fig S7, Table S5). Double helix imaging may show less software-to-software variation and large Z-range at low spot density than astigmatic imaging because the PSF shape and intensity are fairly constant as a function of Z – compared to astigmatic imaging, where spot size, shape and intensity vary greatly as a function of Z (Fig S10).

Double helix software performance decreased significantly for the low spot density low SNR condition (best software SMAP-2018), particularly in terms of best-in-class Jaccard index (66 % low SNR vs 77 % high SNR, Fig 3B, S4, S14A). DH Jaccard index was also significantly worse than astigmatism results at either high or low SNR (85 % high SNR, 86 % low SNR). This indicates that it was quite hard to successfully find localizations in the low SNR DH dataset, likely because the large size of the DH PSF spreads emitted photons over a large area, lowering effective image SNR. DH PSF designs with reduced Z-range but more compact PSF would likely be less sensitive to this issue^22^.

Double helix software performed poorly on the high spot density datasets at high SNR (best software CSpline^27^), especially in terms of the Jaccard index (Fig 3B, S14A, best lateral RMSE 67 nm, best axial RMSE 69 nm, best Jaccard index 46 %). The poor performance at high spot density is again probably because the large DH PSF size increases spot density and decreases SNR (Fig S10). DHPSF performance at high spot density and low SNR was also not reliable (Fig. 3B, S14A, best software SMAP-2018).

#### Biplane localization microscopy

Best-in-class biplane software (SMAP-2018), at low spot density and for both high and low SNR, delivered the best performance in any modality (high SNR: lateral RMSE 12.3 nm, axial RMSE 21.7 nm, Jaccard 87 %), despite a slightly decreased image SNR for the biplane simulations (*Methods*). We observed a significant spread in software performance in terms of lateral RMSE and Jaccard index, with the best-in-class software significantly outperforming the other competitors (Fig S14B, 2D). At low spot density, best-in-class biplane software (SMAP-2018) showed good performance as a function of Z, with high Jaccard index over almost the entire Z-range of the simulations, and with a Z-range of 1200 nm at high SNR (Fig S7, Table S5). The axial RMSE was relatively uniform as a function of Z and close to the CRLB limit (Fig S7). As axial and lateral RMSE are both averaged over the entire Z-range, the strong biplane results arise from good performance across a large Z-range (Fig S7).

At high spot density and high SNR, best-in-class biplane software (SMAP-2018) showed acceptable super-resolved performance (Fig 3B, S3, S14B, best lateral RMSE 43 nm, best axial RMSE 49 nm, best Jaccard index 61 %). Uniquely among the 3D modalities, best-in-class biplane software also gave acceptable performance at high spot density and low SNR (Fig 3B, S3, S14B, best lateral RMSE 55 nm, best axial RMSE 72 nm, best Jaccard index 61 %, best software SMAP-2018).

### Performance of 2D software

Alongside the 3D challenge, we ran a second edition of the 2D localization microscopy software challenge^7^ to assess how the latest 2D software performed on more challenging, more realistic datasets, and to provide an assessment of how the field had progressed since the last challenge. We used the new simulation software, including an experimentally derived PSF and a realistic blinking model, and also simulated a very high spot density condition (5 molecules/μm^2^). We created a more spatially extended test structure, “pseudo-endoplasmic reticulum” (pseudo-ER), composed of 150 nm diameter hollow tubes, to avoid artefacts due to 1D simulated structures^28^. We generated two different imaging conditions with overall similar SNR but different brightness properties; one with low fluorophore brightness and low autofluorescence (the low SNR condition for the 3D challenge, designed to simulate fluorescent protein based SMLM, Fig S5) and one with high fluorophore brightness and high autofluorescence (to simulate affinity-dye-based live cell SMLM, Fig S6). We used lateral RMSE, Jaccard index and overall lateral efficiency to rank the 2D software (Fig 2, S2, Table S1).

For the pseudo-ER dataset, at low density, best-in-class software (ADCG) performed well (Fig. S2, S5), with a Jaccard index of 90 % and lateral RMSE of 31 nm, substantially better than the class average (Jaccard index 72 %, lateral RMSE 36 nm). Low density results for the dimmer fluorophore microtubules dataset were similar to the brighter pseudo-ER dataset (Fig S2, best software SMolPhot). For the very high density 2D dataset, which had 25x higher spot density than the LD dataset, best-in-class software (ADCG) showed excellent performance, with Jaccard index of 75% and lateral RMSE of 45.5 nm (Fig S2). Best-in-class performance (ADCG) on the dimmer fluorophore data at high spot density was also strong (Fig S2, best Jaccard index 70 %, best lateral RMSE 51 nm).

### Algorithms

We identified several classes of algorithm participant software (Table S1):

1. *Non-iterative* software tends to regroup the pixels in the local neighborhood of the candidates, like interpolation, center of mass (QuickPALM^29^) or template matching (WTM^30^). These (often older) algorithms are fast but tend to achieve poor performance (Table S1).
2. S*ingle emitter fitting* software is usually built on a multi-step strategy of detection, spot localization, and optional spot rejection. The detection step finds bright spots in noisy images on the pixel grid. The selection of candidates is usually performed by local maximum search after a denoising filter. Others rely on more complex algorithms like the wavelet transform (*e.g.*, WaveTracer^31^). We did not observe software ranking to depend significantly on the choice of optimization scheme, least-square, weighted least-square or maximum-likelihood estimator (Table S1).
3. *Multi-emitter fitting* software groups clusters of overlapping spots, and simultaneously fits multiple model PSFs to the data. Typically, fitted spots are added to the cluster until a stopping condition is met^4,5^. This leads to improved localization performance at high spot density, at the cost of reduced speed. This class of software (*e.g.*, 3D-DAOSTORM^14^, CSpline, PeakFit^32^, ThunderSTORM^33^) was amongst the top performers in each 2D and 3D competition category (Table S1). As expected, single- and multiple-emitter fitting methods both performed well on low density data (Table S1); apparently at the densities studied, exclusion of occasionally overlapping spots by single-emitter software is sufficient for strong performance; explicit multi-emitter fitting is not required. For the 2D challenge, multi-emitter fitting showed a clear advantage over single emitter fitting at high density (Table S1). Surprisingly however, well-tuned single-emitter fitting algorithms (SMolPhot, SMAP-2018) outperformed multi-emitter algorithms for the 3D high density conditions.
4. *Compressed sensing algorithms*. One subset of these algorithms utilize deconvolution with sparsity constraints to reconstruct super-resolved images^34–36^. Although deconvolution approaches can give good results, they are limited by the necessary use of a sub-pixel grid; increased localization precision requires smaller grid resolution, which must be balanced against increased computational time. Recent approaches address this issue by localizing the point sources in a grid-less manner using an alternating descent conditional gradient scheme under some sparsity constraint (ADCG^37^, SMfit, SOLAR_STORM, TVSTORM^38^). This software class consistently gave the overall best performance for 2D high-density (ADCG^37^ 1^st^, FALCON^36^ 2^nd^, SMfit 3^rd^).
5. *Other approaches*. Of the alternative algorithmic approaches used (Table S1), the annihilating filter-based method LEAP^39^ gave good performance for biplane imaging (the only modality for which it was entered). Recently, we received the first challenge submission from a deep learning SMLM software (DECODE); these promising preliminary results are available on the competition website.

#### Post-hoc temporal grouping

Because molecule on-time is stochastically distributed across multiple frames, a common post-processing approach to improve localization precision is to group molecules detected multiple times in adjacent frames, and average their position^40^ (Supplementary Note 3). Temporal grouping was used by the top performers (including SMolPhot, MIATool^41^ and SMAP-2018), and is visibly apparent as a more punctate super-resolved image (Fig 4A).

#### Choice of PSF model

Most software used a variant of Gaussian PSF model. A few participants designed more accurate PSF models (Table S1). Either diffraction theory was used (MIATool^41^, LEAP^39^) or spline fitting of an analytical function to the experimental PSF was adopted (CSpline, SMAP-2018). Although simple Gaussian model PSFs were sufficient to obtain best-in-class performance for the 2D and astigmatic modalities (ADCG^37^, PeakFit, SMolPhot), top results for the more optically complex biplane and double helix modalities were exclusively PSF-modelling algorithms (SMAP-2018, CSpline, MIATool, LEAP).

#### Multi-algorithm packages

Several software packages take a Swiss army knife approach of integrating multiple optional localization algorithms into one program, to be flexible enough to suit various experimental conditions^20,33^. SMAP-2018 and ThunderSTORM achieved strong across-the-board performance supporting this rationale.

#### Software run time

Software run time is an important parameter for ease of use, and to facilitate real time analysis. We did not see any correlation between software localization performance (Efficiency) and software run time (Fig S24A). We thus created an alternative ranking metric, *Efficiency-Runtime*, which gave 25 % weighting to run time (Supplementary Note 1.7, Fig S24B). Many good performers in the efficiency-only ranking were relatively fast and thus retained good ranking (SMAP-2018, SMolPhot, 3D-DAOSTORM). Interestingly, two software packages highly optimized for speed gained top ranking in this analysis: pSMLM-3D^42^ and QC-STORM.

#### Diagnostic tools for software and algorithm performance

During our analysis, we frequently noticed common types of deviation between software results and ground truth which were easily diagnosed by visual inspection of the super-resolved comparison overlay of ground truth and observed localizations (Fig S19-20). This included not only obvious issues of poor localization precision or spot averaging at high density, but also other problems such as a common error of structural warping which significantly reduced software performance. On the competition website, we provide detailed diagnostic software reports including multiple examples of software performance on individual frames which should help developers to identify algorithm and software limitations and maximize software performance (Fig S21-22).

### Assessment of software performance on real 3D STORM data

We investigated the performance of a representative subset of astigmatism software on real STORM datasets of well-characterized test structures, microtubules and nuclear pore complex, NPC (Fig 4B, S15). This qualitative assessment was consistent with findings for simulated data. No performance difference between single and multi-emitter fitters was observed, which is not surprising since spot density in these datasets was low. Relatively poor software performance was immediately obvious from visual inspection (QuickPALM). Temporal grouping noticeably improved resolution (3D DAOSTORM, CSpline, MIAtool, SMAP-2018). Gaussian fitting software (3D DAOSTORM, MIATool, ThunderSTORM) gave robust nanoscale resolution images. Interestingly, experimental PSF fitting software (CSpline, SMAP-2018) gave noticeably improved resolution of fine structural features such as the top and bottom of the NPC (Fig 4B) or the hollow core of antibody-labelled microtubules (Fig S15).

## DISCUSSION

We performed the first broad evaluation of software for 3D single molecule localization microscopy, to assess the state of the field and to allow non-specialists to determine the optimal software for their experiments.

In order to provide a realistic assessment of 3D software performance we tested software on simulations incorporating experimentally acquired microscope point spread functions. Our experimental-PSF-derived simulation approach is readily adaptable to novel engineered 3D SMLM PSFs^43^ or to the PSF of individual microscopes. For instance, it would be possible to combine our derived-PSF approach with the SMLM sample simulation tool SuReSim^44^ in order to generate ultra-realistic synthetic data, which could then be personalized to each experimentalists sample and microscope, to easily determine the blocker factors to maximal resolution, for a given experiment.

The strongest conclusion we draw from the 3D localization microscopy challenge is that choice of localization software greatly affects the quality of final super-resolution data, even at “easy” high SNR, low spot density conditions. Biplane performance was particularly dependent on software choice, with only one software (SMAP-2018) achieving near-Cramér-Rao lower bound performance. Double helix SMLM showed less sensitivity to choice of software than biplane, with astigmatic SMLM intermediate between the two. The best software in each modality performed close to the Cramér-Rao lower bounds over a wide focal range and successfully detected most molecules, even at low signal to noise. Average software in all three modalities was significantly worse, with the obtained axial resolution being particularly sensitive to software choice.

The second major conclusion of the 3D challenge is that localization software that explicitly includes the experimental PSF in the fitting model gives a significant performance increase for 3D SMLM. For the more optically complex biplane and double helix modalities in particular, the best results were exclusively from software using PSF modelling approaches (SMAP-2018, CSpline, MIATool). This result also highlights the need for experimental PSF modelling not only in SMLM software, but also emphasizes the high degree of experimental realism required of SMLM simulations. The clear performance advantage of experimental PSF modelling software in the 3D software challenge would have been unobservable had it been run with a simple Gaussian PSF.

Of the different algorithm classes, well-tuned single-emitter and multi-emitter fitting algorithms (each capable of dealing well with occasional molecule overlap) gave good results for low density 3D SMLM. We also found that several software packages for astigmatic or biplane imaging gave adequate performance for the challenging case of high molecule densities, as long as the image SNR was high. Current software packages gave poor performance when molecule density was high and image SNR was low. These results suggest that, at least with current algorithms, high density 3D SMLM performance is mediocre at high SNR, and poor at low SNR. Surprisingly, multi-emitter fitting did not show significant improvement over well-tuned single emitter fitting for the 3D high-density datasets; this may indicate that significant potential for improvement remains in this category.

Many software packages did not apply temporal grouping^40^, resulting in reduced software performance. Since temporal grouping is a simple step for maximum precision, we urge all software developers to integrate this approach into their software as an optional final step in the localization process.

The second 2D localization microscopy challenge provided the opportunity to reassess the state of the field. The performance of best-in-class 2D software over a range of conditions, at both high and low spot density, is excellent. The performance of the best-in-class software at high spot density (ADCG^37^) was only moderately decreased compared with the low spot density results, with nearly identical molecule detection performance, and a 30 % increase in localization error. Interestingly, the top three performers in the 2D high density condition were all compressed sensing algorithms (ADCG^37^, FALCON^36^, SMfit). In low density 2D conditions, the best single-emitter, multi-emitter and compressed sensing algorithms all gave comparable, excellent, performance. We speculate that performance in this category may now be near optimal levels.

In future we plan to extend the SMLM challenge website and software into an open platform where the assessment process is fully automated, and where new competition simulations and assessment metrics can easily be created and contributed by the community. Scientific CMOS cameras are rapidly becoming a major platform for single molecule localization microscopy^6^ and it will be important to include sCMOS simulations in future SMLM software assessments. Furthermore, there remain two important classes of super-resolution microscopy for which software performance is crucial, but no broad software assessment has yet been performed: fluorescence-fluctuation-based super-resolution microscopies (*e.g*., 3B^45^, SOFI^46^, SRRF^47^) and structured illumination microscopy^48^.

The results of this competition clearly demonstrate the formidable algorithmic performance of the best 2D and 3D localization microscopy software. However, a key outstanding challenge that often hinders adoption of new algorithms is that only a small subset of algorithms are packaged in, or compatible with fast, well-maintained, user-friendly software packages, which include all stages of the SMLM data analysis pipeline – analysis, visualization and quantification. One solution would be for the SMLM software community to collectively adopt both a standard data format and a single software platform for future software development, such as FIJI/ ImageJ^49^. Any new algorithm released in this environment could be immediately and widely adopted by users, and easily integrated into existing packages for SMLM analysis, visualization and quantification.

Both the 3D and 2D localization challenges remain open and continuously updated on the competition website. This continuously evolving analysis of state of the art super-resolution software performance provides a valuable resource to super-resolution microscopists, helping to ensure they use software that gets the best out of hard-won data. It also provides SMLM software developers with a robust means of benchmarking new algorithms against current state of the art.

## Supporting information

## ACKNOWLEDGEMENTS

Authors acknowledge the following funding sources: a Newcastle University Research Fellowship and a Wellcome Trust & Royal Society Sir Henry Dale Fellowship grant number 206670/Z/17/Z to SH; an European Research Council (ERC) under the European Union’s Horizon 2020 research and innovation programme, Grant Agreement no. 692726 to DS, TAP, MU; UK BBSRC grants BB/M022374/1, BB/P027431/1, BB/R000697/1 grant and MRC grants MC-UU-12018/2, MR/K015826/1 to RH; European Research Council (ERC) grant CoG-724489, CellStructure to JR; National Institutes of Health grant 1R15GM128166-01 to GMH; and NSF SBIR grants 1353638, 1534745 to Double Helix LLC. We thank R. Piestun at University of Colorado for providing DH-PSF phase mask designs to Double Helix LLC. We thank all the localization microscopy challenge participants for their contribution: Hazen Babcock (3D-DAOSTORM, Cspline, L1H), Fabian Hauser (3D-STORM Tools), Shigeo Watanabe (3D-WTM,WTM), Nicholas Boyd (ADCG), Junhong Min, Kyong Jin and Jong Chul Ye (ALOHA, FALCON), Hervé Rouault (B-recs), Emmanuel Soubies (CEL0-STORM), Artur Speiser, Srinivas Turagas and Jakob Macke (DECODE), Alex von Diezmann, Camille Bayas and W. E. Moerner (Easy-DHPSF), Thomas Vomhof and Jochen Reichel (FIRESTORM), Hanjie Pan (LEAP), Ann Wheeler (Localizer), Zhen-li Huang and Yujie Wang (MaLiang), J. Chao, R. Velmurugan, A. V. Abraham and R. J. Ober (MIATool), Hendrik Deschout (mlePALM), Thomas Pengo (Octane, PeakSelector), Yi-na Wang (PALMER), Alex Herbert (PeakFit), Koen Martens and Johannes Hohlbein (pSMLM-3D), Luchang Li (QC-STORM), Ricardo Henriques (QuickPALM), G. Tamas and J. Sinko (RainSTORM), Steve Wolter and Markus Sauer (RapidSTORM), Manfred Kirchgessner and Frederik Gruell (SFP Estimator), Yiming Li and Jonas Ries (SMAP), Hayato Ikoma (SMfit), A. Loot, A. Valdmann, M. Eltermann, M. Kree and M. Pärs (SMolPhot), Yoon J. Jung, Anthony Barsic Rafael Pietsun, and Nikta Fakhri (SOLAR_STORM), Anna Archetti (STORMChaser), Martin Ovesny, Guy Hagen and Pavel Krizek (ThunderSTORM), Jiaqing Huang (TVSTORM), Adel Kechkar and Jean-Baptiste Sibarita (WaveTracer) and Benoît Lelandais (ZOLA-3D). We thank the SMLMS 2016 organizers (S. Manley and A. Radenovic, EPFL) for hosting a localization microscopy challenge special session. We also thank Double Helix LLC and Molecular Devices LLC for sponsoring the SMLMS 2016 special session. The sponsors had no input or influence on the research.

## AUTHOR CONTRIBUTIONS

DS and SH conceived and coordinated the study. DS, SH, TAP, AAr, HB, SC, AW, GMH, RH, TL, TP, JBS designed the study. SH, AAg, RH, JBS collected experimental PSFs. DS, TAP, SH, TL wrote simulation code. BR shared unpublished software. DS generated simulated datasets. JR shared experimental STORM data. AH, JR, JC, RV provided feedback and quality control on simulations and analysis methods. TAP carried out the assessment of software performance. TAP, DS, SH analysed and interpreted the results. DS, HB, RO, BR, GMH, JBS, JR, RH, MU, SH directed research. SH, DS, TAP wrote the manuscript with feedback from all authors.

## METHODS

### 1. CHALLENGE ORGANIZATION

We first ran the 3D SMLM software challenge as a time limited competition, with a results session hosted as a special session of the 6^th^ Annual Single Molecule Localization Microscopy Symposium in August 2016. The competition has now been converted to a permanent software challenge accepting new submissions. Special mention to the software SMAP and 3D-WTM that participate to our eight categories (*density* × *modality*). The current list of participants is at: http://bigwww.epfl.ch/smlm/challenge2016/index.html?p=participants

All datasets, methods, participations, and results of the challenge 2016 made available at http://bigwww.epfl.ch/smlm/challenge2016/. Software for simulation and analysis is hosted on the competition GitHub repository: https://github.com/SMLM-Challenge/Challenge2016/

## 2. LOCALIZATION MICROSCOPY SIMULATIONS

### 2.1. Structure

The synthetic datasets were designed to be similar to images derived from cellular structures in real experimental conditions. We defined mathematical models for cellular structures that imitate cytoskeletal filaments such as microtubules and larger tubular structures such as the endoplasmic reticulum or mitochondria (Fig S1A). These structures have a tubular shape in the 3D space. Psuedo-microtubules are defined with their central axis elongating in a 3D space having an average outer diameter of 25 nm with an inner, hollow tube of 15 nm diameter. Pseudo-endoplasmic reticulum is defined as having a diameter of approximately 150nm.

The underlying sample structure is formalized in a continuous space which allows rendering of digital images at any scale, from very high resolution (up to 1 nm/pixel) to low resolution (camera resolution: 100 nm/ pixel). The continuous-domain 3D curve is represented by means of a polynomial spline. The sample is imaged in a 6.4 × 6.4 μm^2^ field of view, and the center lines of the microtubules have limited variation along the z (vertical) axis, *i.e*., less than 1.5 μm. The fluorescent markers are uniform randomly distributed over the structure according to the required density. The photon emission rate of each fluorophore is controlled by a photo-activation model (see below).

The exact locations of all fluorophores are stored at high precision floating-point numbers expressed in nanometers. This ground-truth file is useful for conducting objective evaluations without human bias.

## 2.2. Photophysics activation model

Given a list of source locations from the structure simulator, fluorophore blinking was simulated by a 4-states Markov chain model. The states are ON, OFF, BLEACH, DARK and the transition are Poisson distributed (Fig S1C), except for the OFF to ON transitions which follow a uniform random distribution, to reflect that in typical experimental conditions, constant imaging density is maintained by tuning the photoactivation rate during the experiment. All switching is calculated at sub-frame resolution and then total fluorophore on-time was integrated over each frame.

Due to two decay paths, the actual mean lifetime of the state ON is

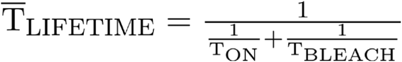

Switching rates were chosen to approximate photoactivatable fluorescent proteins T_ON_ = 3 frame, T_DARK_ = 2.5 frames, and T_BLEACH_=1.5 frames.

Fractional fluorophore ON-times per frame (between 0 and 1) were then multiplied by the mean flux of photon emission. The flux of photons expressed in photons/seconds was given by the relation

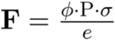

Φ is the quantum yield of the dye, P is power of the laser in W/cm^2^, e = h c / λ is the energy of one photon, σ = 1000 ln(10) ε / N_A_ is the absorption cross section in cm2 and ε is the molar extinction coefficient (EC) or absorptivity in cm^2^/mol which is a characteristic of a given fluorophore. The laser power was Gaussian distributed over the field of view. At the end of this process a list of XY positions, on-frames and (noise-free) intensities for all activated fluorophores was obtained.

Analysis of the resulting simulated photon counting distribution is presented in Supplementary Note 1.

## 2.3. Experimental Point-Spread Function

Model PSFs, stored as high resolution look up tables, were derived from experimentally measured PSFs. Although the algorithmic approach is distinct, this concept of accurately modelling the experimental PSF based on calibration data bears relation to the PSF phase retrieval approach previously employed by Hanser and coworkers^50^.

Images of fluorescent beads were recorded for each modality (Table S4). Signal to noise ratio of recorded PSFs was maximized in all cases by maximizing exposure time and averaging over several frames to increase dynamic range.

To acquire experimental PSFs, we took 100 nm Tetraspek beads (Invitrogen) adsorbed to #1.5 (170 μm thick) coverglass, imaged in water. The excitation wavelength was between 640 nm and 647 nm, and a Cy5 emission filter was used. Data acquisition parameters for each modality are listed in Table S4.

## 2.4. Simulation PSF construction

For each modality, 3-6 beads were selected within a small (< 32 μm) region, to minimize PSF variation due to spherical aberration. Images for each selected bead were interpolated in XY to a pixel size of 10 nm. Beads were then coaligned by cross-correlation on the in-focus frame. Coaligned beads were averaged in XY to minimize pixel quantization artefacts and to increase SNR. Where necessary, Z-stacks were interpolated to a Z-step size of 10 nm. A central Z-range of 1.5 μm was selected that represents 151 optical planes with a Z-step of 10 nm. The Z-range covers −750 nm to +750 nm. The plane of best focus was chosen as the simulation 0 nm plane. Each model PSF was normalized such that the total intensity of the PSF in the in-focus frame within a diameter of 3 FWHM from the PSF center was equal to 1.

For the DH PSF, the transmission of the combined phase mask system was measured as 96 %, which was approximated as 100 % brightness relative to the 2D and astigmatic PSFs.

In biplane super-resolution microscopy, emitted fluorescence is split into two simultaneously imaged channels, with a small (500-1000 nm) defocus introduced between the two channels^12^. As the small defocus should introduce minimal additional aberration into an optical system, we semi-synthetically constructed a realistic biplane PSF from the experimental 2D PSF. The two defocused PSFs were constructed by duplicating the 2D PSF and offsetting it by −250 nm and 250 nm for each Z-plane.

This yielded five high SNR model PSFs with an isotropic voxel size of 10×10×10 nm^3^.

The ground truth XY=0 was defined as the image centre of mass of the in-focus frame of the model PSF, and Z=0 was defined as the in-focus frame. Accounts for shifts in the fitted XY centre of the model PSF by localization software due to systematic offsets and Z-dependent variation of the model PSF centre of mass are dealt with below (wobble correction).

## 2.5. Noise model

A constant mean autofluorescent background was added to the noise-free simulated images, and these images were then fed through the noise model representing Poisson distributed fluorescence emission recorded on a high quantum efficiency back-illuminated EMCCD^51,52^.

The proposed noise model assumed as main contributions to the stochastic noise:

- σ_*S*_, the shot noise produced by the fluorescence background and signal and the spurious charge. Shot noise can be derived from the second moment of the Poisson distribution
- σ_*R*_, the read noise of EMCCD camera, which is described by second moment of the Gaussian distribution
- σ_*EM*_, the electron multiplication noise introduced by the gain process, which is described by the second moment of the Gamma distribution^52^.

We assumed as camera parameters the ones specified for the Photometrics Evolve Delta 512 EMCCD camera:

- QE = 0.9, Evolve quantum efficiency at 700 nm absorption wavelength.
- σ_*R*_ = 74.4 electrons, manufacturer measured root mean square noise for Evolve 512 camera
- c = 0.002 electrons, manufacturer quoted spurious charge (clock induced charge only, dark counts negligible)
- EM_gain_ = 300
- e_adu_ = 45 electron per analog to digital unit (ADU), analog to digital conversion factor
- G = 0.9*300/45 = 6, total system gain
- BL = 100 ADU

The final simulated photon electrons will thus be given by:

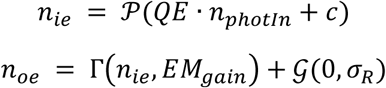

which leads to the final pixel counts:

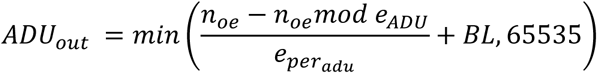

## 2.6. Depth-dependent lateral distortion: Wobble

As the PSF models are experimentally derived, the 3D estimated localizations exhibit a depth-dependent lateral distortion, here called *wobble*. This optical distortion is due to a combination of a systematic offset (arbitrary definition of PSF center) and optical aberrations^53^. In order to compare estimated and true localizations, we correct this effect during the assessment (Section 3.1).

## 2.7. Comparison of software results between different modalities

The intensities of the PSF in each imaging modality were normalized to facilitate comparison of results between different modalities. Software results between 2D, 3D AS and 3D DH modalities are expected to be directly comparable.

For the biplane model PSF, as the emitted fluorescence is split into two channels, the intensity in each of the two simulated biplane channels was additionally reduced by 50 %. We note that the fluorescence background was not reduced by 50 % as intended, leading to artificially high background for the biplane simulation. *I.e*., the background in each of the two biplane channels is the same as in the single channel of the other modalities. However, due to the low background level in the 3D simulations, the effect on image SNR and thus localization error is small (see Fig S7), less than 5 nm near the plane of focus. Therefore, as long as the small drop in image SNR is taken into account, approximate comparisons of the biplane data to the other modalities can still be made.

## 3. SOFTWARE ASSESSMENT

### 3.1 Protocol

Each localization file submitted by the participants was manually checked for erroneous systematic errors in the definition of the dataset coordinate system, such as offsets, XY axis flips or clear scaling errors. Datasets were then programmatically standardized into a consistent output format. All modifications are publicly available. If required, the modifications consisted of columns reordering, reversing axes, XY axis swap, and shifting the lateral positions by a half camera pixel.

The assessment pipeline includes three main parts: localization processing, the pairing between true and estimated localization and the metrics calculations. The first one depends on the assessment settings. There are two switchable properties: photon thresholding and wobble correction. Their combinations yield four different assessment settings. Up to 64 assessment runs per software were possible (*i.e.*, 4 modalities, 4 datasets per modality). For any setting, we excluded the fluorophores within a lateral distance of 450 nm from the border. This value corresponds to the radius of the largest PSF, *i.e.*, Double Helix. The activations too close from the border are more difficult to localize and could bias the results.

The pairing between true and estimated localizations was performed frame by frame. The procedure matches two sets of localizations. We deployed the presorted nearest-neighbor search for its efficiency, with a linking threshold of 250 nm. The results are effectively similar to the computationally intensive Hungarian algorithm^7^.

#### Photon thresholding

A photon threshold was required primarily due to the use of a realistic fluorophore blinking model. Since a fluorophore could activate/ bleach at any point in a simulated frame, this led to many frames containing very dim, undetectable localizations, eg. where a molecule had been active for one or more frames previously, and then bleached during the first 5 % of a frame. These fractional localizations should also be present but practically undetectable in an experimental dataset.

We decided to focus the software analysis on the localizations where the molecule was active for the majority of a frame, to be consistent with experimental expectations. Therefore, we implemented a photon threshold means where we kept the 75% brightest ground truth fluorophore activations. Because this was performed *after* the pairing step, observed localizations that were paired to discarded ground truth activations were also removed from the metric calculations.

#### Wobble correction

The centroid of experimental point spread functions shifts laterally by as much as 50 nm, as a function of axial position^10,53^. This is most often ignored by localization software, and instead corrected post-hoc by reference to a calibration curve^37^. Since our simulated PSF is experimentally derived, it was necessary to correct for these artefactual shifts between the observed localizations and ground truth, as part of the assessment process. This correction was performed using calibration data uploaded by competitors, similar to the correction typically performed on experimental data^53^.

Three scenarios were proposed to the participants: no correction was applied during the assessment; the correction was based on a file provided by the participant itself or the correction was calculated by ourselves. The latter nevertheless requires the participant to localize a stack of beads we provided. Since the true positions of the beads are known, the difference between the estimated and true positions could be calculated and averaged. It thus yields the values for wobble correction.

In certain specific cases (identified on the competition website), at the request of authors, we did not apply this correction, for example because the software explicitly considered the whole 3D PSF during fitting and was thus immune to this lateral shift artefact. For accurate results, application of lateral shift correction is critical for analysis of localization microscopy simulations using experimentally derived PSFs, as can be seen by comparison of typical software results with and without wobble correction (Fig S11).

## 3.2 Metrics

We calculated a large number of analysis metrics to quantify the performance of software relative to ground truth. These are discussed in detail in Supplementary Note 1. The metrics are split into two categories: localization based and image based metrics.

The former directly relies on the localizations positions and notably includes the Recall, the Precision, the Jaccard Index, the RMSE (axial and lateral) and the consolidated Z-range. For the calculation of average software performance (Fig 3B) outlier software with an efficiency less than *eff=-30* were excluded from the measurement.

The image based metrics are computed from a rendered image and includes the Signal-to-Noise Ratio (SNR) and the Fourier Ring / Shell Correlation (FRC/FSC). To render the image, we added the contribution of each localized molecule at the corresponding pixels. A contribution takes the form of a 3D additive Gaussian with a Full-Width Half Maximum (FWHM) of 20 nm. A complete list of all computed metrics is shown in the Supplementary Note 2.

We also calculated localization based metric results as a function of axial position. We proceeded by considering a subset of activations lying within an interval of axial positions (*i.e.*, from the true localizations). Then, most of the metrics (*e.g.*, Recall) are locally computed. This yields a curve providing information on the depth performance of each software / modality.

In order to summarize software axial performance, we analyzed how the recall varied as a function of Z. A typical recall versus axial position curve (Fig S9) will drop at positions far from the focal plane, *i.e.*, where software can no longer detect spots to defocus. We first smoothed the curve using a sliding window. Then we computed the software Z-range, defined as the full width half maximal Recall of the smoothed curve (Fig S12). This quantity is visually intuitive and useful for discussion of the recall performance if considered alongside a plot of recall vs axial position. However, because FHWM recall depends on the maximal recall, ranking based on this procedure would promote a software which poorly performed everywhere (*i.e.*, flat curve), whereas a software which performed well in the focal plane but less well outside would obtain a worse FWHM recall. This observation leads us to produce a so-called consolidated Z-range, by multiplying the Z-range value by the maximal Recall, which should provide a robust metric that avoids the previous case scenario.

### Principal component analysis

In order to analyse the relationship between analysis metrics we computed the covariance matrix between each metric and the principal component analysis (PCA) on the metrics (Fig S14B). Each metric was standardized before applying the covariance and the PCA. For convenience, we took the additive inverse of the metrics for which lower values are best (*i.e.*, FP, FN, RMSE, FRC, FSC).

Summary statistics and detailed results for each software are available on the competition website (http://bigwww.epfl.ch/smlm/challenge2016/index.html?p=results), which also includes a tool for side-by-side comparison of the results of multiple software packages

## 3.3 Baseline Localization Software

We developed a minimalist Java tool software that performs localizations of bright emitters on the 4 modalities of the challenge 2016: 2D, Astigmatism, Double-Helix, and Biplane. This SMLM_BaselineLocalization software is only designed to establish the performance baseline for the SMLM challenge. It has intentionally limited lines of code and relies only on few threshold parameters to localize particles. It has basic calibration tool that has to run on a z-stack of beads to find the linear f(x) relation between the axial position Z and the shape of the bead.

- Astigmatism: Z = f(W_X_ − W_Y_), where W_X_ and W_Y_ are respectively an estimation of the size in X and Y.
- Double-Helix: Z = f(θ), where θ is the angle formed the pairing of two close points.
- Biplane: Z = f (W_left_ − W_right_), where W_left_ and W_right_ are respectively an estimation of the size of the spots in left and the right plane.

The Java code is available: https://github.com/SMLM-Challenge/Challenge2016

## 4. REAL DATA ASSESSMENT

Astigmatism software was tested on previously published real 3D STORM datasets of microtubules and nuclear pore complex^20^. The tubulin dataset corresponds to the raw data for Fig S6 in Ref 20, and the nuclear pore complex dataset corresponds to raw data for Fig S9 in Ref 20. Key acquisition parameters for data analysis are summarized on the competition website.

Data were analyzed by software authors or expert users, and submitted via the competition website. All data were drift corrected via cross-correlation. STORM images were rendered with a constant Gaussian blur with 3 nm standard deviation and saturated by 0.1 – 0.5 %. The complete scripts used for assessment and image rendering are available on the competition GitHub page.

## 5. DATA AVAILABILITY

### 5.1 Data availability statement

Simulated competition datasets are available at http://bigwww.epfl.ch/smlm/challenge2016/, together with the parameters used to generate the data. The ground truth list of simulated molecule positions for each competition dataset remains secret in order to allow the software challenge to remain continuously open to new submissions. However, ground truth data is available for the simulated training datasets.

### 5.2 Code availability statement

All software is available at https://github.com/SMLM-Challenge/Challenge2016

