## Supplementary material for "Super-resolution fight club: A broad assessment of 2D & 3D single-molecule localization microscopy software"

1: Biomedical Imaging Group, School of Engineering, Ecole Polytechnique Fédérale de Lausanne (EPFL), Switzerland
2: Harvard Center for Advanced Imaging, Harvard University, Cambridge, Massachusetts, USA
3: Laboratory of Nanoscale Biology & Laboratoire d'Optique Biomédicale, STI - IBI, EPFL, Lausanne, Switzerland
4: University of Minnesota Informatics Institute, University of Minnesota Twin Cities, USA
5: Electrical Engineering, University of Texas Dallas, Richardson, Texas, USA
6: MRC Genome Damage and Stability Centre, School of Life Sciences, University of Sussex, Brighton, UK
7 : Double Helix LLC, Boulder, Colorado, USA
8 : Istituto Italiano di Tecnologia, Genova, Italy
9: Edinburgh Super-Resolution Imaging Consortium, University of Edinburgh, UK
10 : Laboratory of Experimental Biophysics, École Polytechnique Fédérale de Lausanne (EPFL), Lausanne, Switzerland
11: Department of Imaging Physics, Faculty of Applied Sciences, Delft University of Technology, The Netherlands
12: UCCS center for the Biofrontiers Institute, University of Colorado at Colorado Springs, Colorado, USA
13: Institut Interdisciplinaire de Neurosciences, University of Bordeaux, France
14: European Molecular Biology Laboratory (EMBL), Cell Biology and Biophysics Unit, Heidelberg, Germany
15: Quantitative Imaging and Nanobiophysics Group, MRC Laboratory for Molecular Cell Biology, University College London, UK
16: Centre for Bacterial Cell Biology, Institute for Cell and Molecular Biosciences, Newcastle University, UK

#### SUPPLEMENTARY FIGURES AND TABLES

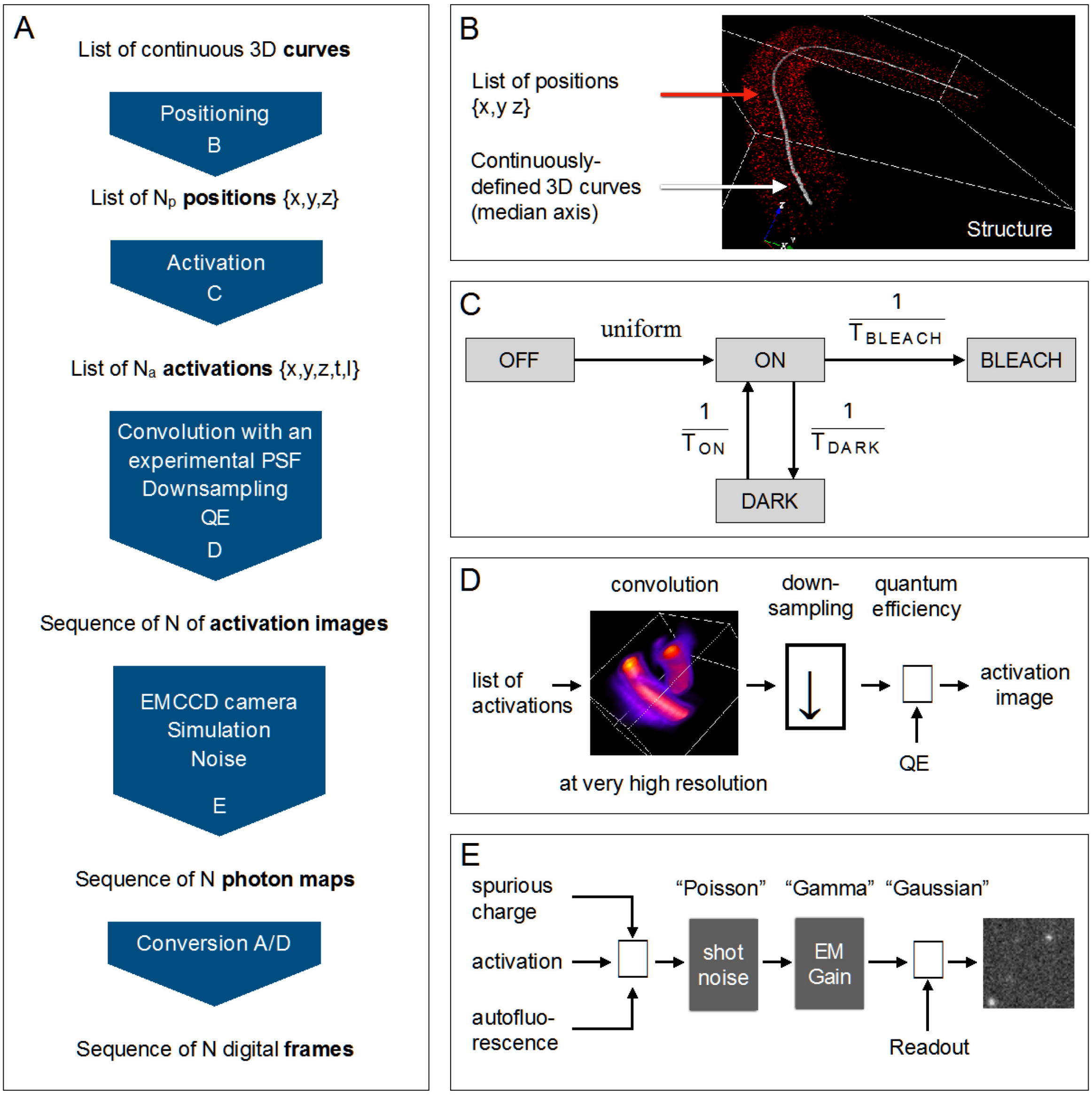


***Figure S1**: Simulations*. ***A***. The overall steps of the simulation are shown, from the 3D geometry of the structure to the sequence of digital frames. ***B***. The geometry of the structure is constructed from a series of 3D tubes where every median axis of tubes (white line) is a 3D curve continuously defined by a three B-spline functions in the volume of interest. The membranes of the tubes are densely populated with possible positions of fluorophores according a given radius and a given thickness of membrane. ***C***. The activation process takes the list of positions and activates the fluorophores following the 4-states model of activation. The parameters $T_{BLEACH}$, $T_{ON}$ and $T_{DARK}$ allow to control the blinking rate and the lifetime of the emitters. The output of the activation consist to a list of activations defined by a position (x,y,z), a time (t) and a number of emitted photons (I) in the current frame. ***D***. The creation of the sequence run frame-to-frame. All activations of a given frame are convolved with the experimental PSF at high resolution (2 nm), and reduced to the camera resolution. The number of photons per pixel is converted to electron by pixel using the quantum efficiency (QE) of the camera. ***E.*** The noise model is applied to the three sources of electrons: the activation of the emitters, the spurious charge of the camera and the autofluorescence background (photons converted to electrons using QE). The shot noise follows a Poisson distribution; the EMCCD gain is simulated by a Gamma function; and a Gaussian noise is added to simulate the read-out noise. Finally, the acquired photons map at a given time are converted to digital frame in 16-bits, a baseline is also added.

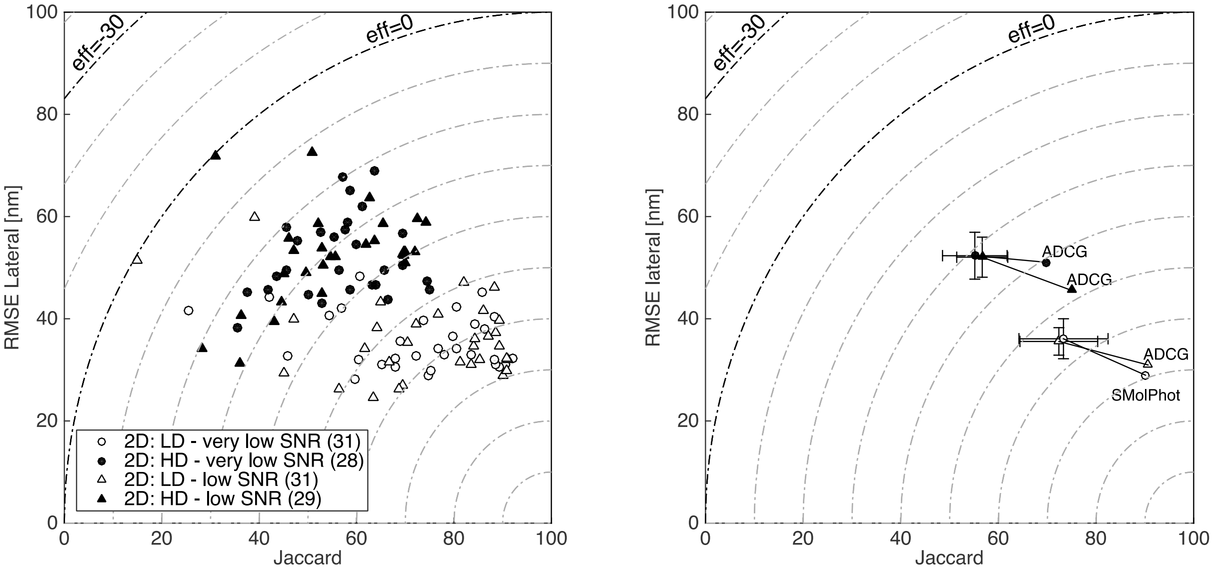


***Figure S2***: *Comparison of 2D software performance*. Dashed lines indicate efficiency (higher is better). ***A.*** Performance of all 2D SMLM software. ***B.*** Average (marker with *s.d.* error bars) and best-in-class (marker with name) software performance for 2D modalities. LD, low density; HD, high density.

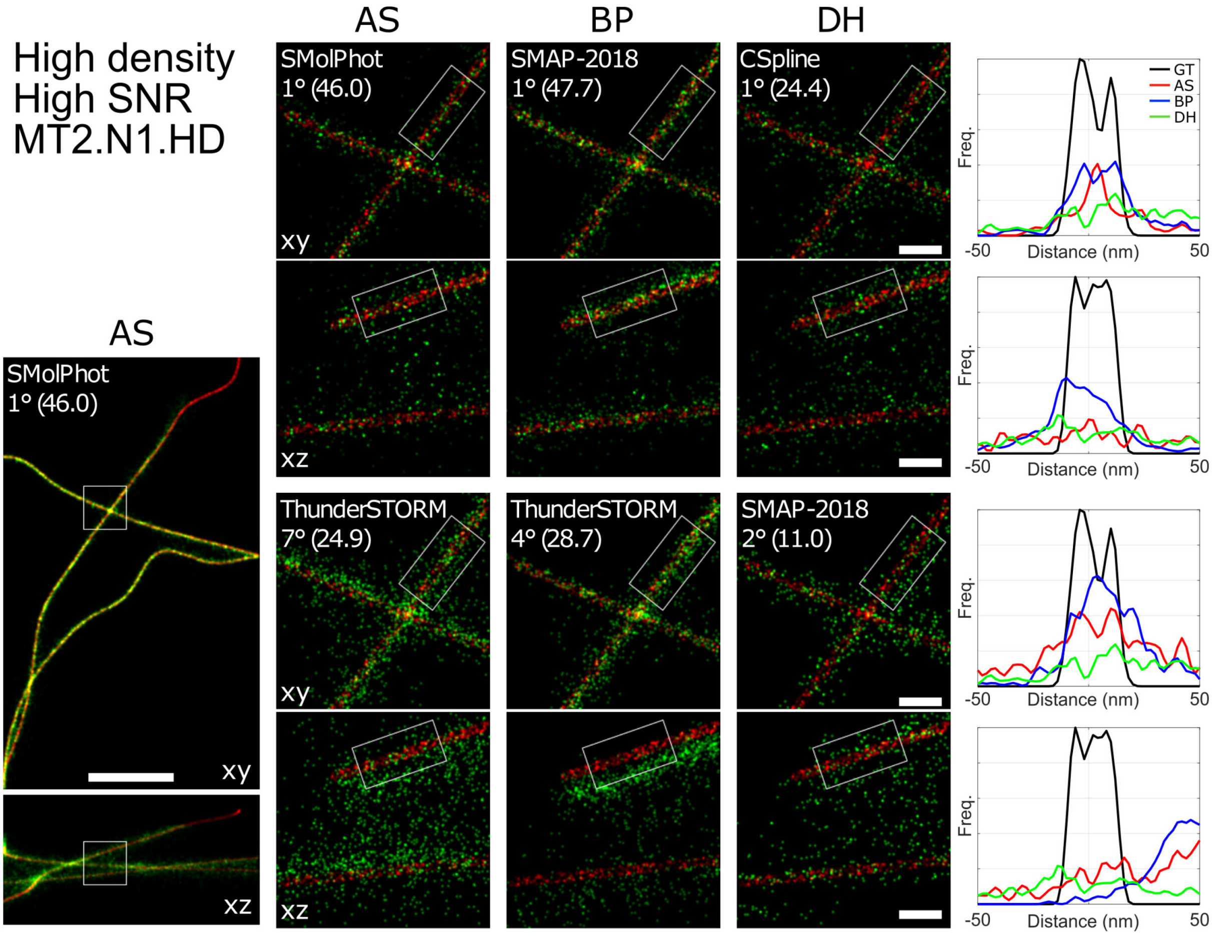


***Figure S3.*** *Super-resolved xy and xz projection images of 3D competition datasets for best-in-class (top) and representative average (bottom) software in each modality, for high SNR high density dataset. Left*: *xy* and *xz* overview images for winning AS software. *Middle*: *xy* and *xz* zoom images of boxed regions in left panel, for winning and mid-range software, each modality. *Right*: *Xy* and *xz* line profiles of winning and mid-range software for each modality, for boxed regions in middle panel. *Image colors*: red, ground truth; green, software results. *Line profiles*: GT, ground truth, black; AS, astigmatism, red; BP, biplane, blue; DH, double helix, green. *Panel key: Software-name Dataset-Ranking° (Efficiency-score). Scale bar*: full image, 1 µm, zoomed regions, 100 nm.

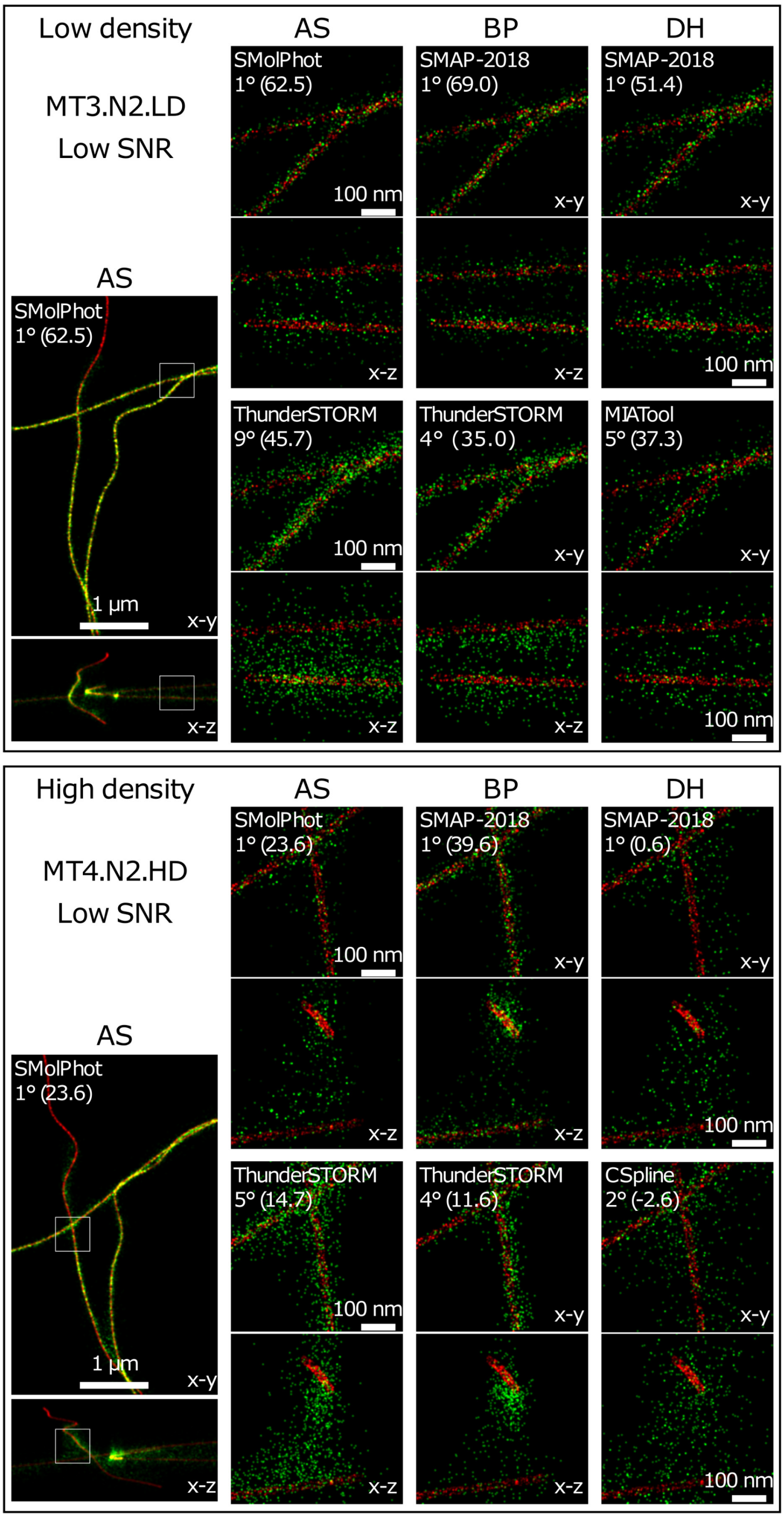


**Figure S4:** *Super-resolved xy and xz projection images of 3D competition datasets for best-in-class (top) and representative average (bottom) software in each modality, for low SNR datasets. Left*: *xy* and *xz* overview images for winning AS software. *Right*: *xy* and *xz* zoom images of boxed regions in left panel, for winning and mid-range software, each modality. *Image colors*: red, ground truth; green, software results. *Panel key: Software-name Dataset-Ranking° (Efficiency-score). Scale bar*: full image, 1 µm, zoomed regions, 100 nm.

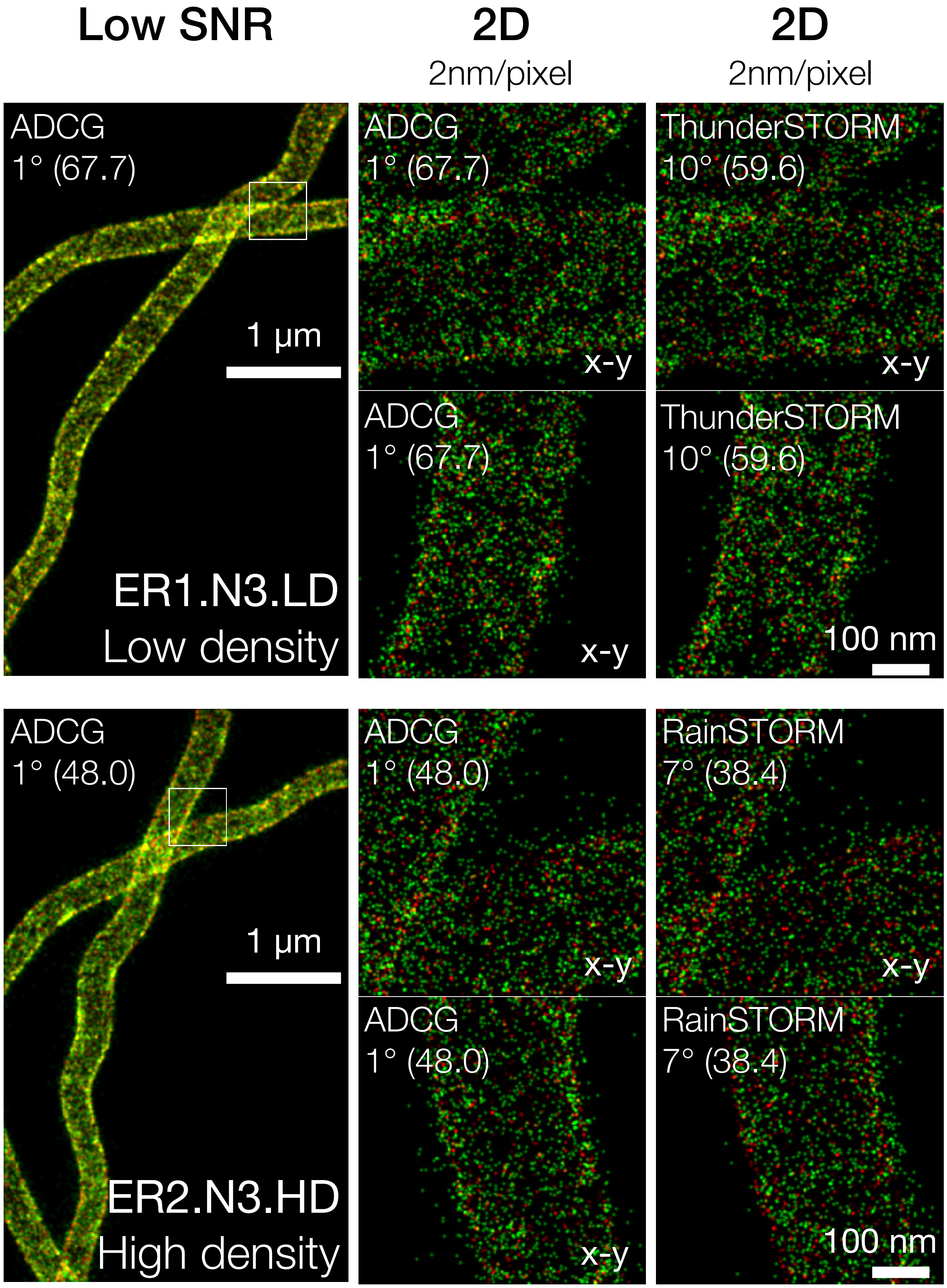


**Figure S5:** Super-resolved images of 2D competition datasets for best-in-class (top) and representative average (bottom) software in each modality, for low SNR psuedo-ER datasets. Box indicates zoomed region. Red, ground truth; green, software results. Panel label key*: Software_name Ranking° (Efficiency).*

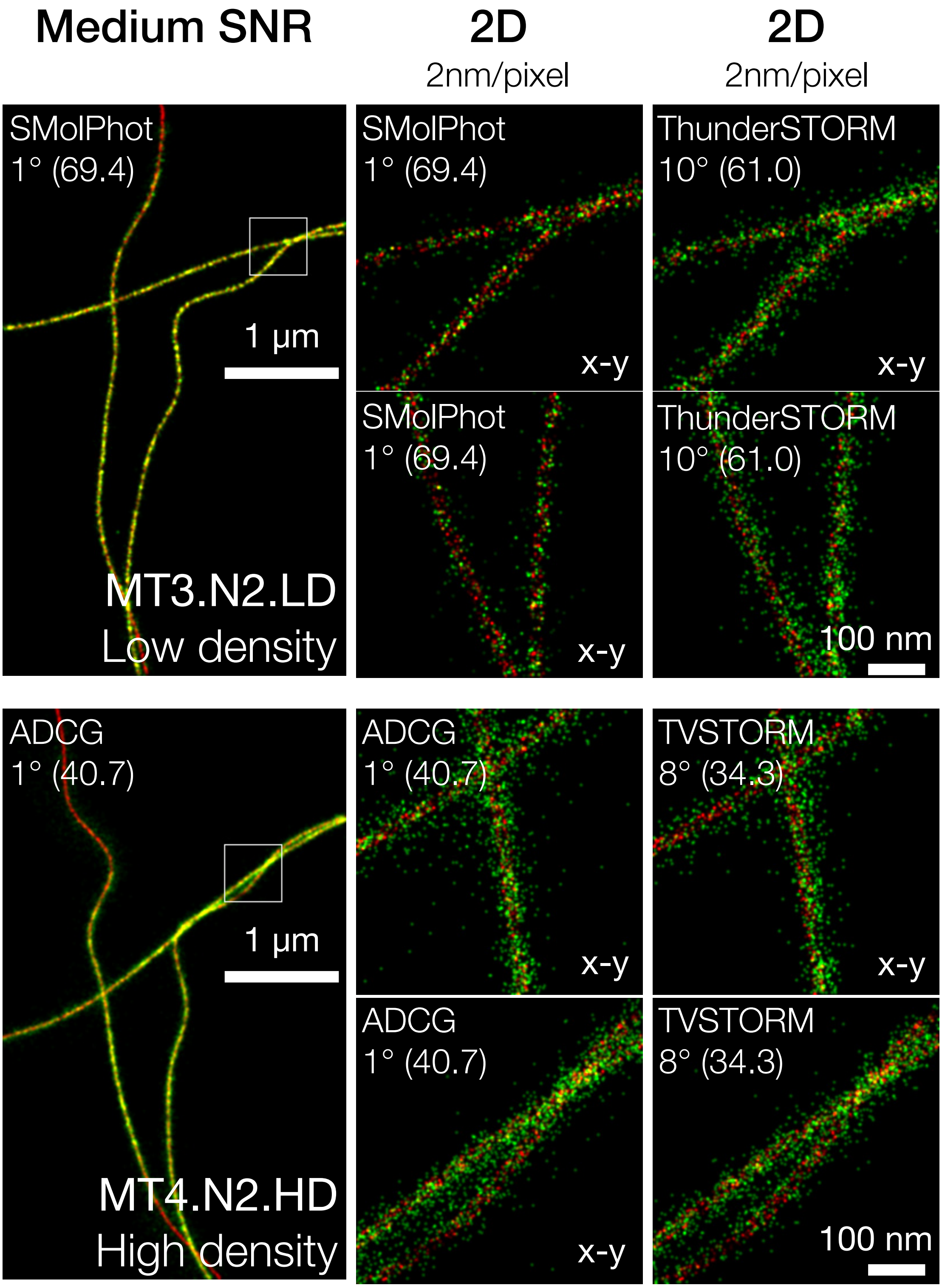


**Figure S6:** Super-resolved images of 2D competition datasets for best-in-class (top) and representative average (bottom) software in each modality, for medium SNR psuedo-microtubule datasets. Box indicates zoomed region. Red, ground truth; green, software results. Panel label key*: Software_name Ranking° (Efficiency).*

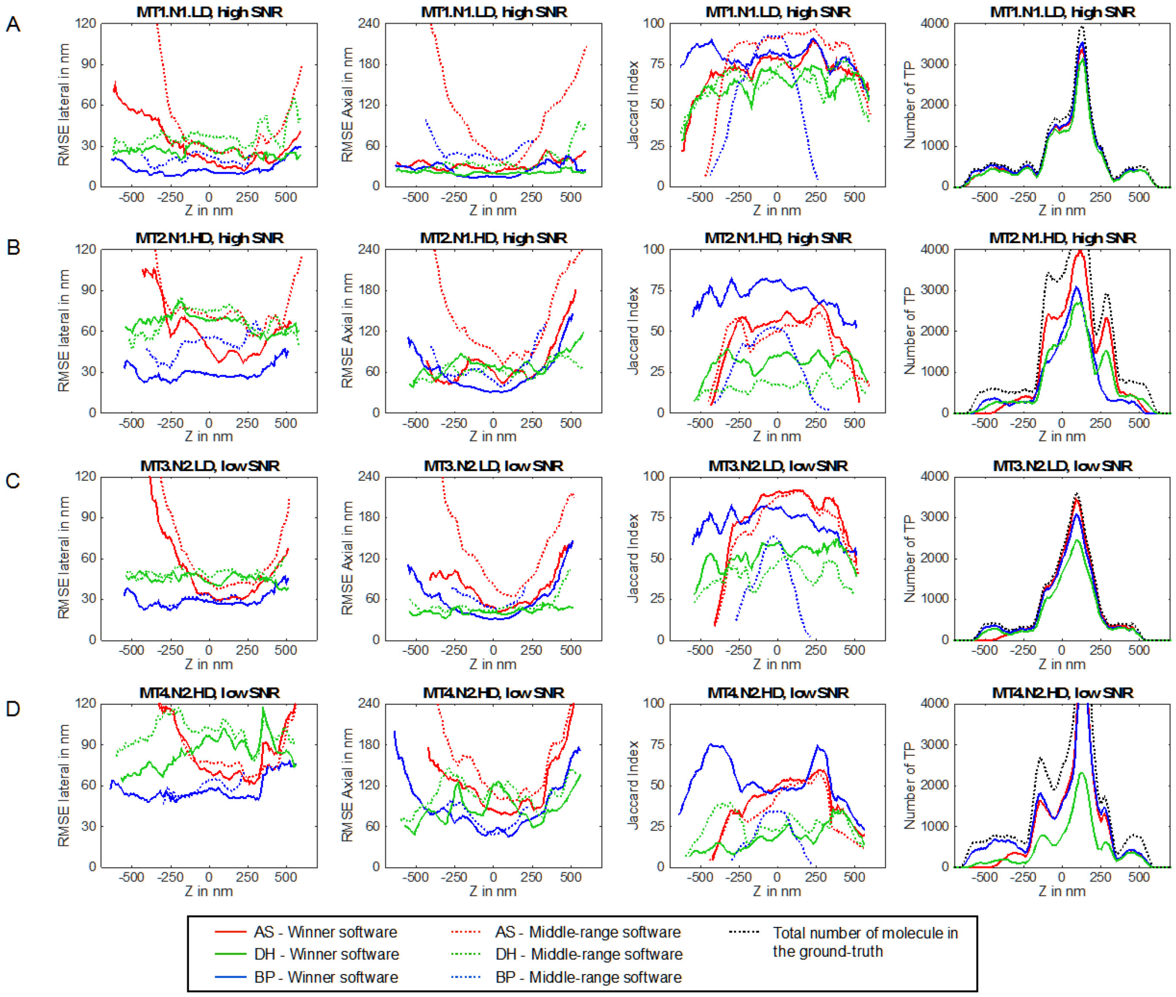


**Figure S7:** *3D software performance as a function of depth Z (axial position of the molecule) for each competition dataset (A-D)*. The metrics are locally computed within a depth interval. Based on the true axial position of the molecules, we exclude the ones out of the depth interval of interest $Z_i$. From this subset and their (previously) paired software localisations, the lateral (1st column) and axial (2nd column) RMSE are computed. The Jaccard Index (3rd column) requires a false positive count for each bin, which is approximated using the non-paired software localisations that fall in $Z_i$. The winners (for AS, DH and BP in red, green and blue respectively (full line)) for each dataset MT1.N1.LD, MT2.N1.HD, MT3.N2.LD, MT4.N2.HD are plotted at the row 1 to 4 respectively. In addition, a software with average performance is also displayed for the metrics lateral and axial RMSE and the Jaccard Index (dotted line). The number of true positive (TP, 4th column) is the number of paired molecule included in $Z_i$. In addition to the winners, the total number of activations per depth interval is also displayed (Ground-truth, dotted black line).

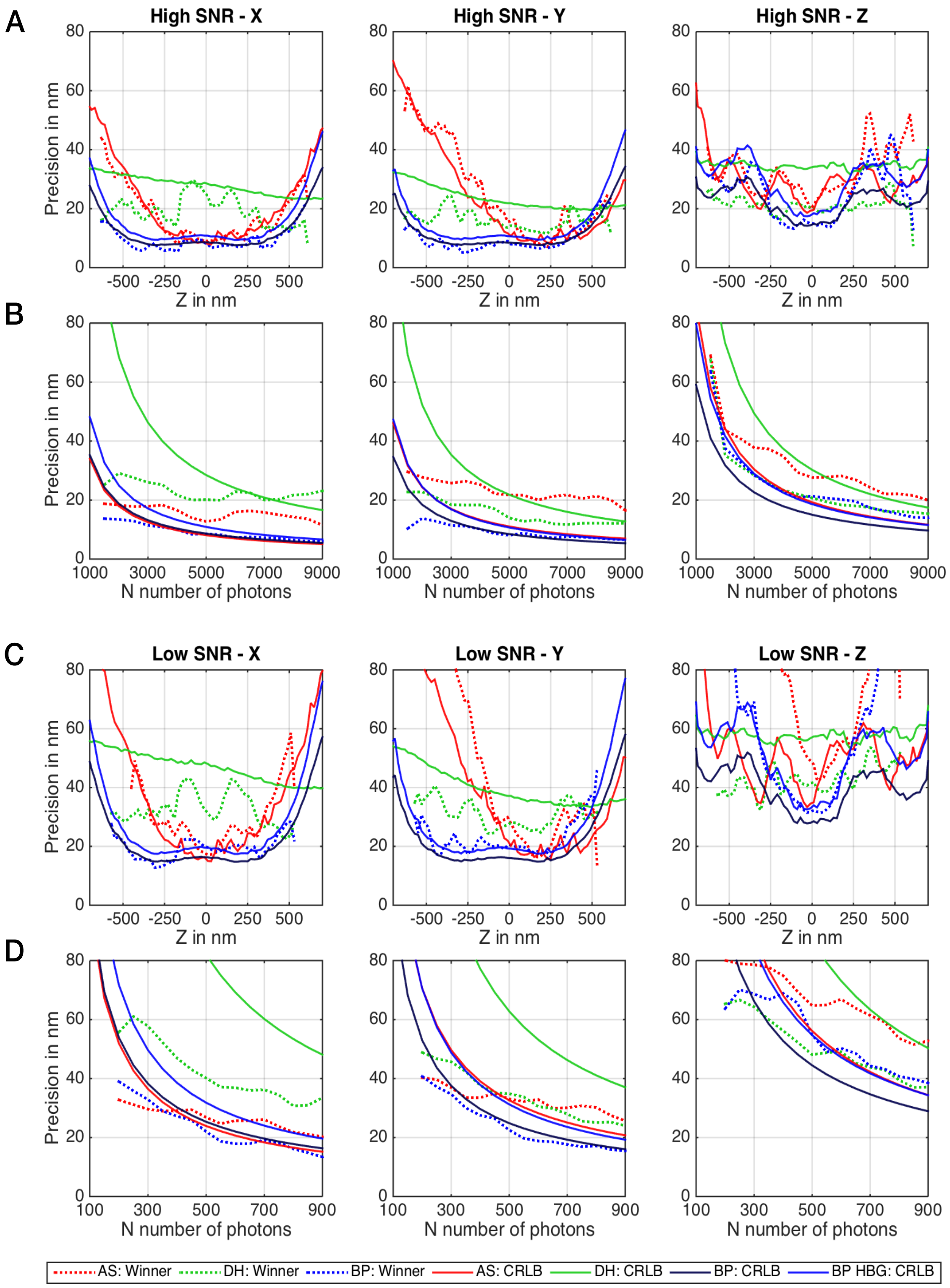
A
High SNR - X
High SNR - Y
High SNR - Z
Precision in nm
Z in nm
B
N number of photons
C
Low SNR - X
Low SNR - Y
Low SNR - Z
D
AS: Winner
DH: Winner
BP: Winner
AS: CRLB
DH: CRLB
BP: CRLB
BP HBG: CRLB

***Figure S8:*** *Comparison inter-modality of the Cramér-Rao lower bounds (CRLB) in high SNR and low SNR conditions on a single frame (without temporal grouping). For each condition, the 1D RMSE performance of the winning software is plotted for comparison with the CRLB limit*[1,2]*.* **A.** CRLB as a function of the depth Z (axial position of the molecule) for noise condition N1 are computed with for each PSF: astigmatism AS; double helix DH; biplane, BP, biplane high background BP HBG. Calculations for AS, DH, BP were for a single molecule emitting N=5’000 photons on a frame with an auto-fluorescence background B=100 photons for AS, DH and BP. BP (black line) is the CRLB if BP simulations were identical to the AS and DH conditions. Actually, the background used for the biplane competition simulations had a higher background, B=200 (BP HBG, blue, BP high background). BP software results should be compared to the BP HBG CRLB (blue line). **B**. CRLBs as a function of the number of emitted photons N for noise condition N1 are computed for each PSF (AS, DH, BP, BP HBG) for a single molecule at the focal plane (Z=0). **C.** CRLBs as a function of depth Z for noise condition N2. **D.** CRLBs as a function of the number of emitted photons N for noise condition N2. The computation of the CRLBs is described in Supplementary Note 4.

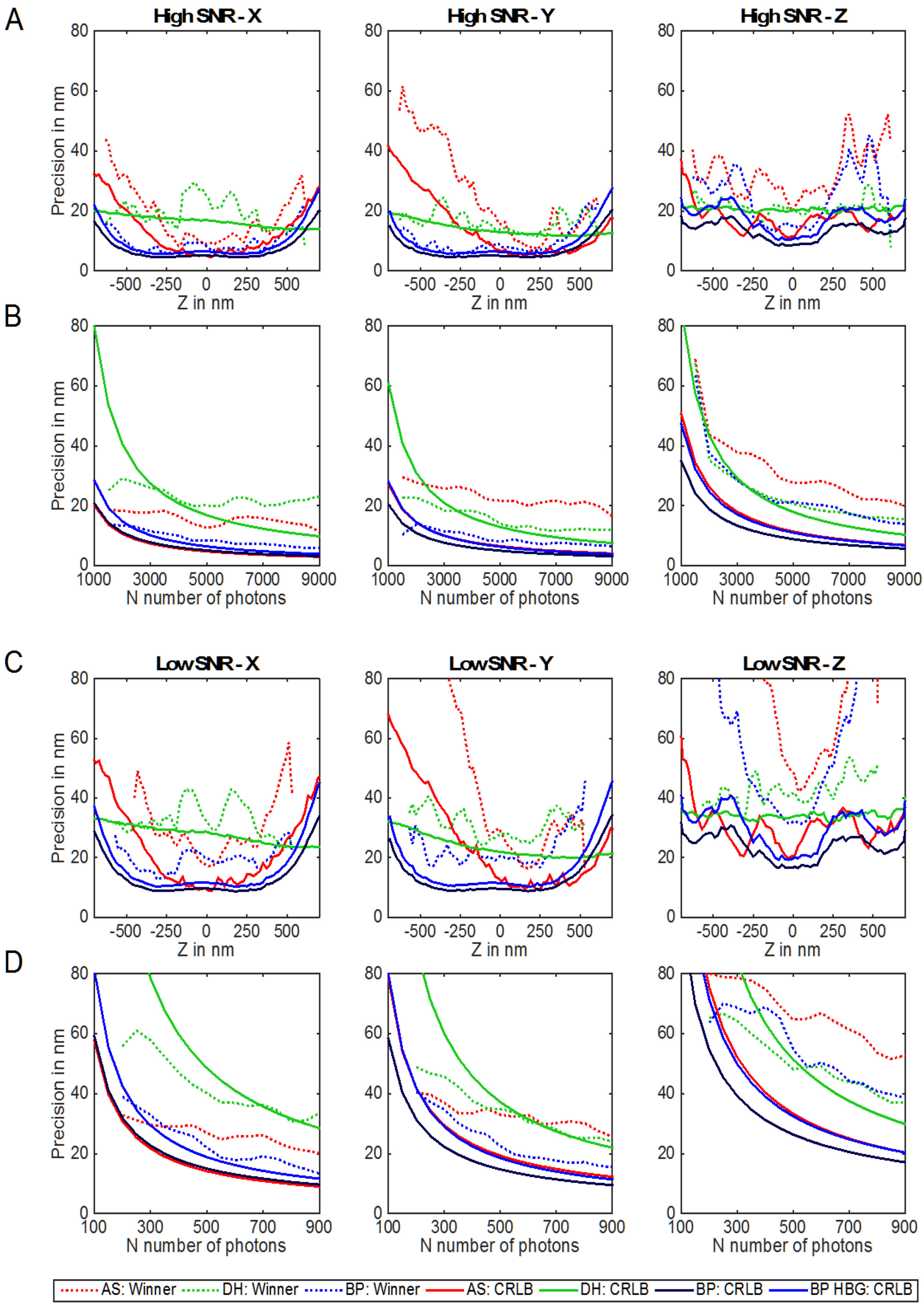
A
High SNR - X
High SNR - Y
High SNR - Z
Precision in nm
Z in nm
B
N number of photons
C
Low SNR - X
Low SNR - Y
Low SNR - Z
D
AS: Winner
DH: Winner
BP: Winner
AS: CRLB
DH: CRLB
BP: CRLB
BP HBG: CRLB

**Figure S*9***: *Comparison of the winner with the Cramér-Rao lower bounds (CRLB), combining localizations from multiple frames (temporal grouping), in high SNR and low SNR conditions (solid lines). For each condition, the 1D RMSE performance of the winning software is plotted for comparison with the CRLB limit*[1,2] *(dashed lines).* **A.** CRLB as a function of the depth Z (axial position of the molecule) are computed for noise condition N1 with each PSF (AS, DH, BP in red, green, blue respectively) for a single molecule emitting N=5'000 photons per frame on F = 2.85 frames (average number of frames for which a molecule is activated) with an auto-fluorescence background B=100 photons for AS, DH and BP. The BP was also displayed for B=200 because it was the setting of the datasets of the competition (BP HBG). BP software results should be compared to the BP HBG CRLB (blue line). **B.** CRLB as a function of the number of emitted photons N per frame for noise condition N1 are computed for each PSF (AS, DH, BP, BP HBG) for a single molecule at the focal plane (Z = 0) for F = 2.85 frames (average number of frames for which a molecule is activated). **C.** CRLBs as a function of depth Z for noise condition N2 for F = 2.85 frames. **D.** CRLBs as a function of the number of emitted photons N for noise condition N2 for F = 2.85 frames. The computation of the CRLBs is described in Supplementary Note 4.

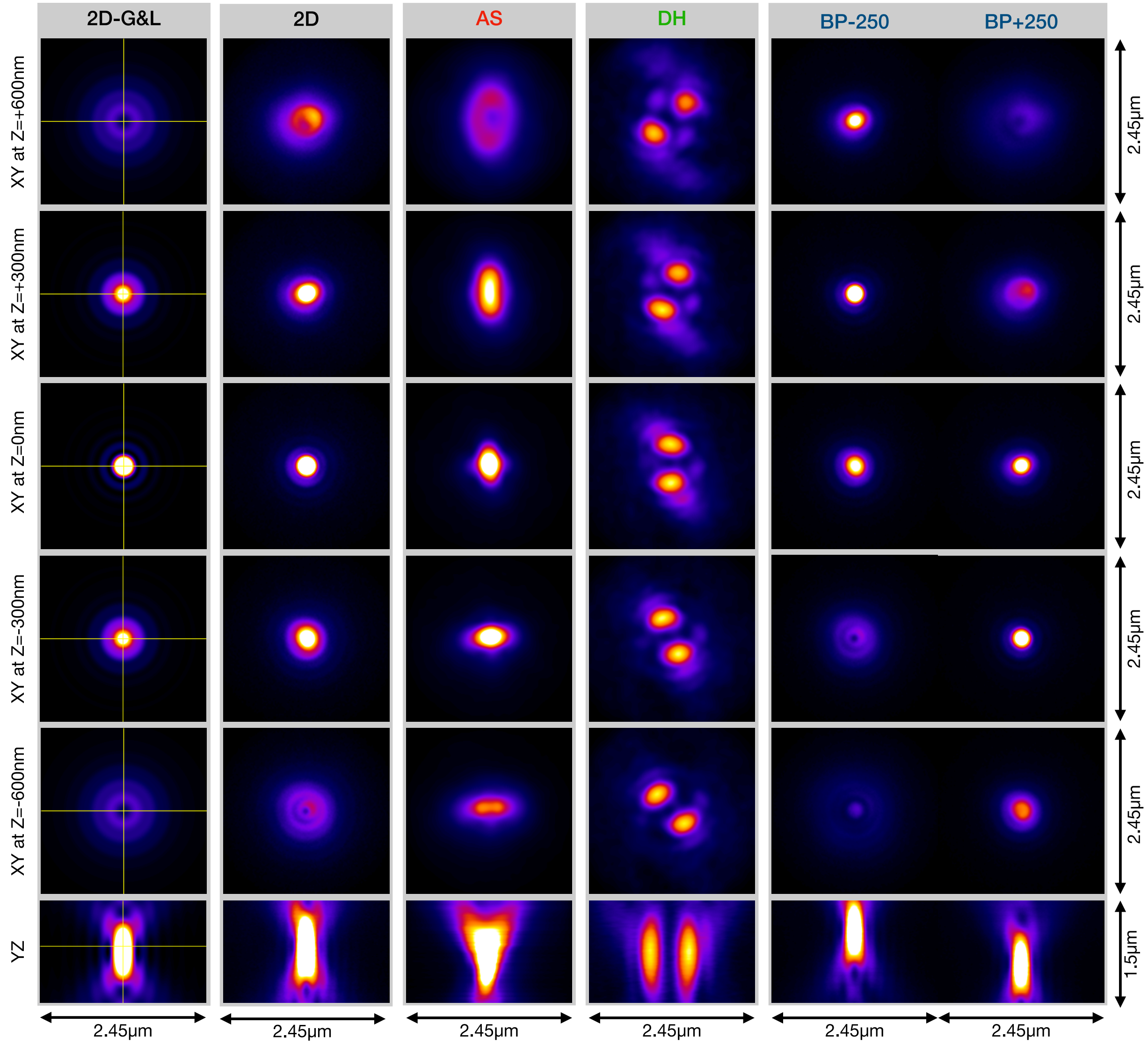


**Figure S*10***: *XY and YZ profiles of experimentally derived competition PSFs for different imaging modalities compared with theoretical Gibson & Lanni PSF*[3]. Experimentally derived PSFs were constructed as described in Online Methods, and under conditions summarized in Table S4. Gibson-Lanni model PSF calculated under corresponding conditions (NA 1.49, 700 nm emission wavelength).

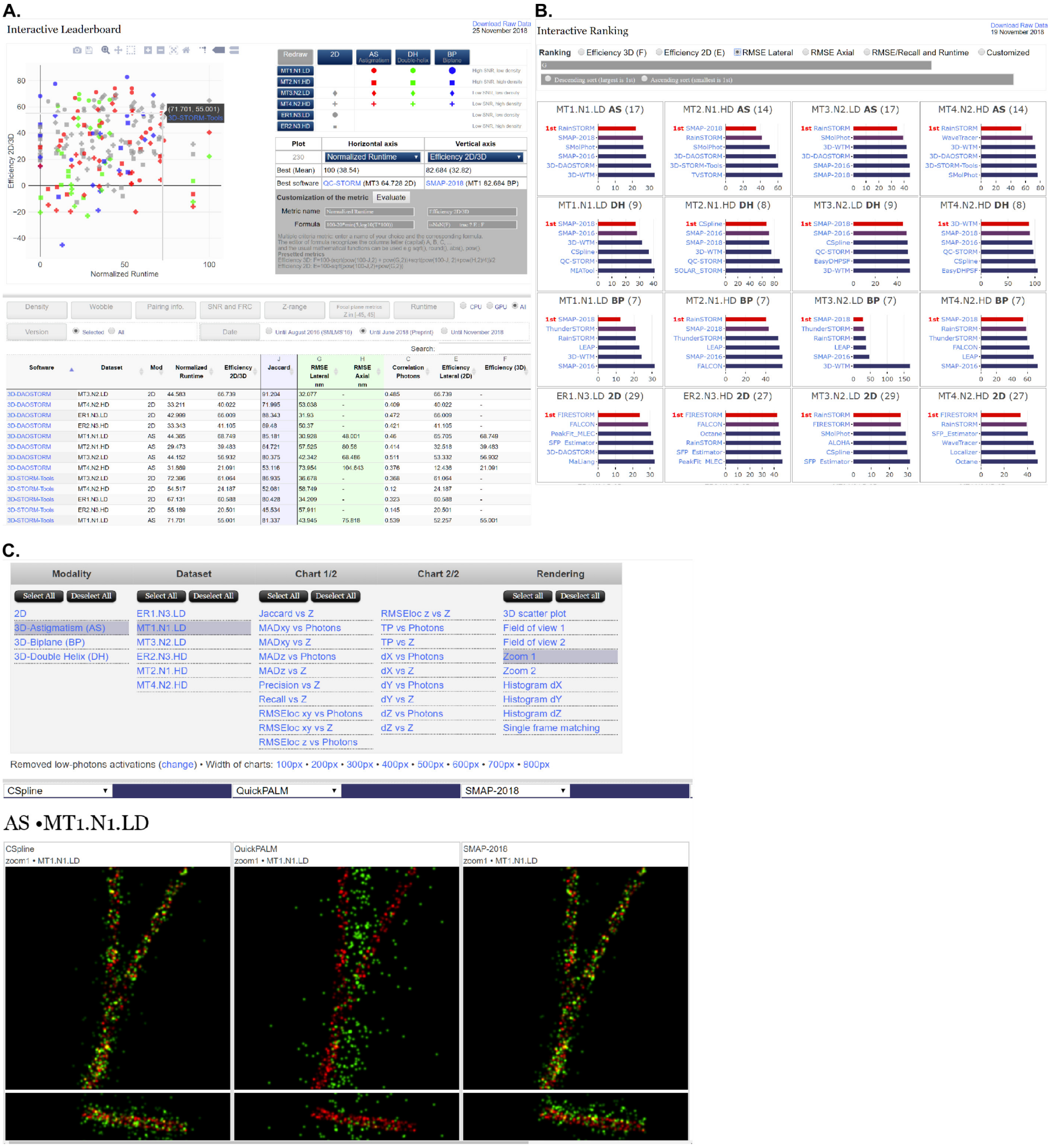


**Figure S11**: *Screenshot of the interactive data exploration tools on competition website*. **A.** [2D plotting tool & leader board](). Software results according to 21 performance metrics can be plotted and explored. Hovering over any data point reveals the corresponding software and individual score. User-defined analysis metrics can be calculated from existing metrics. **B.** *[Software ranking]()*. Software performance can be ranked by pre-defined metrics, or any user-defined combination of existing metrics. **C.** *[Side-by-side software comparison.]()* Results, including super-resolved images, per-frame performance examples and localization precision distributions for up to 5 software packages can be compared side by side.

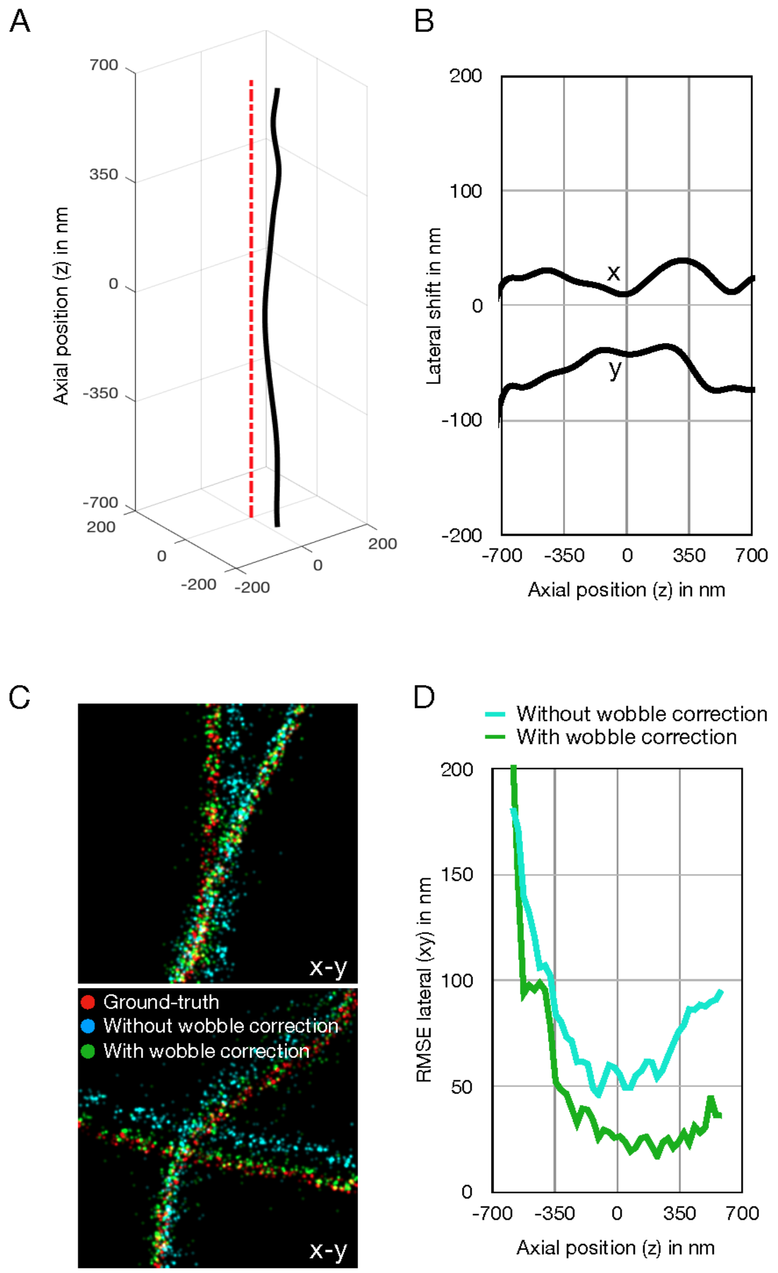


**Figure *S12***: Wobble correction is required for accurate software-simulation comparison with experimentally derived PSFs (unless the software incorporates wobble directly in the analysis model). ***A-B.*** Profile of representative software localization offset as a function of axial position. ***C-D.*** Comparison of representative software versus ground truth results with and without wobble correction.

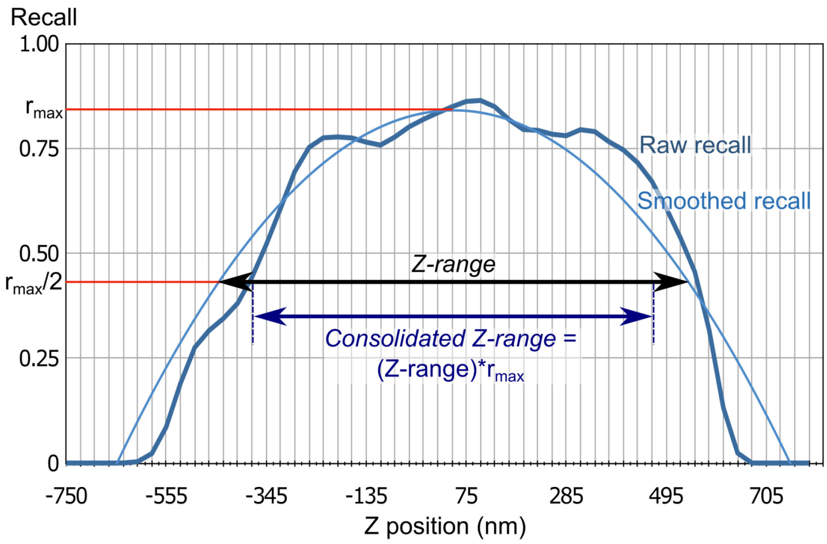


**Figure S*13***: Metrics for software Z-range. *Z-range* measures the range over which software detects >0.5*(max recall) of ground truth molecules. However, a large Z-range is not practically useful if the software maximum recall is very low. The supplementary metric *consolidated Z-range* rescales the FWHM recall by the maximum recall. This will return a low value if either the FWHM recall or the maximum recall of the software is low.

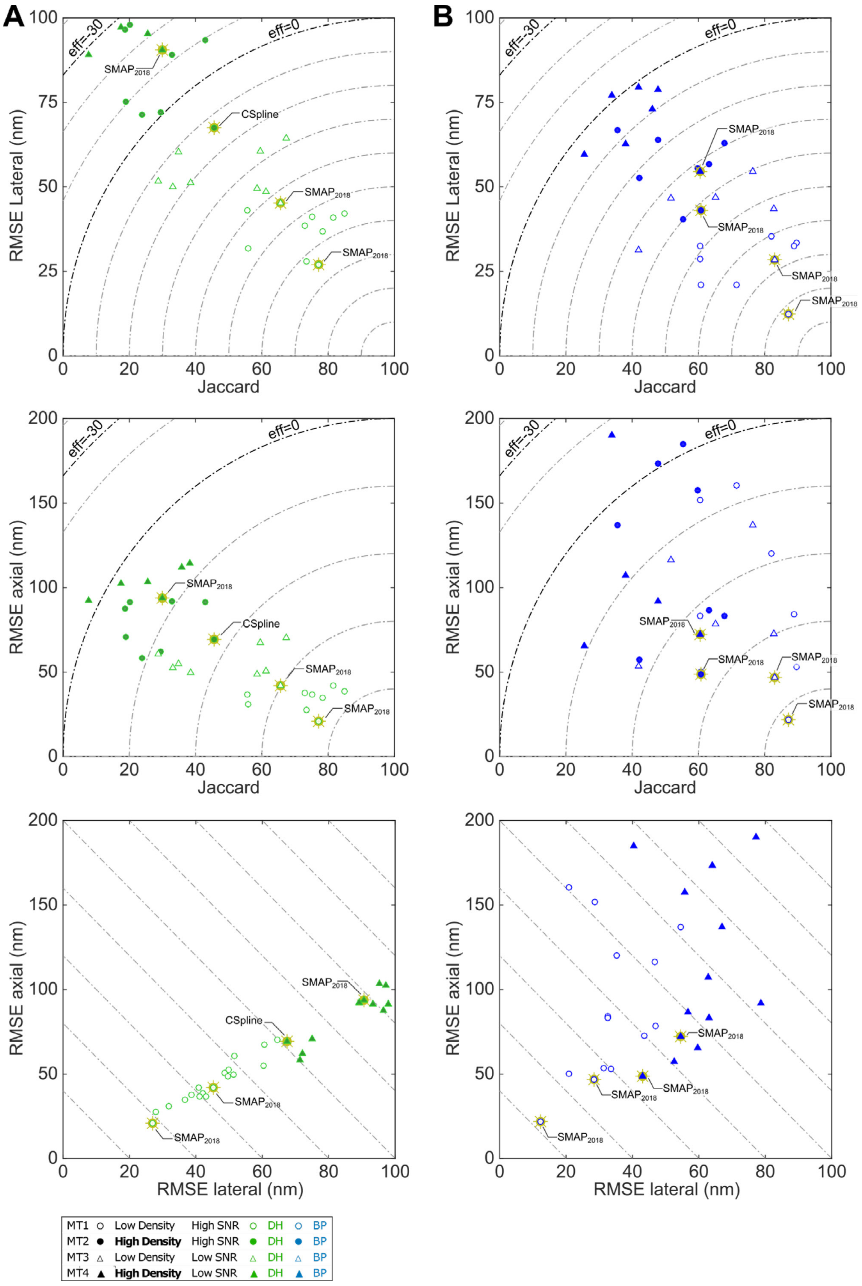


**Figure S14**: *Localization error and spot detection performance for double helix (A, DH) and biplane (B, BP) modalities*. Gold stars indicate top performers for each dataset. Dashed lines in top, middle panels indicate overall efficiency (higher is better).

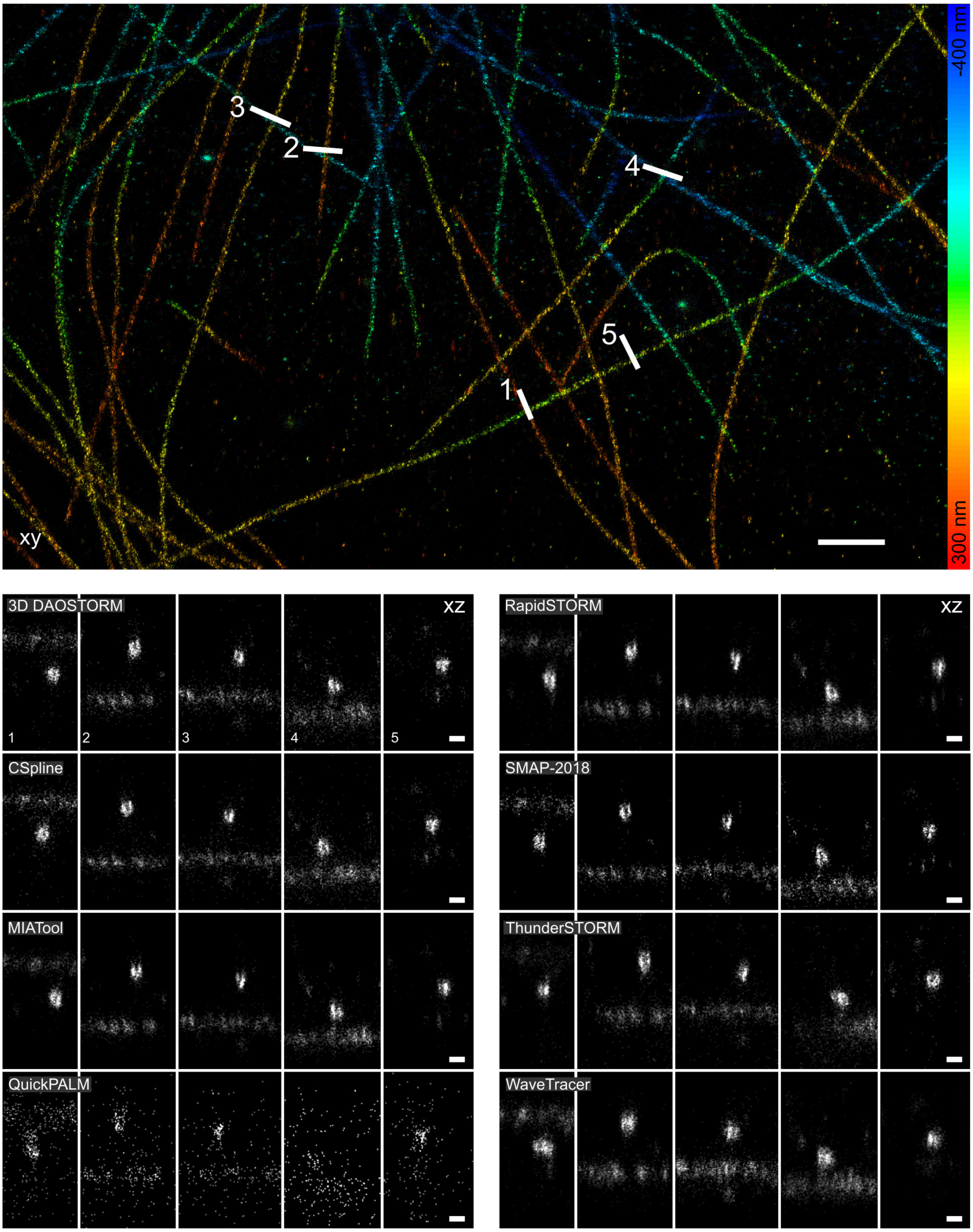


**Figure S15.** *Performance of astigmatism software on real 3D STORM images of microtubules. Top:* Super-resolved overview image in *xy* for 3D-DAOSTORM software, color coded for depth. *Bottom: Xz* orthoslices along numbered line profiles in top panel for 8 astigmatism software packages. The best performing software clearly resolves the hollow core of the labelled microtubules. Line profile thickness, 250 nm. *Scale bars*: *xy* overview, 1 µm; *xz* orthoslices, 0.1 µm.

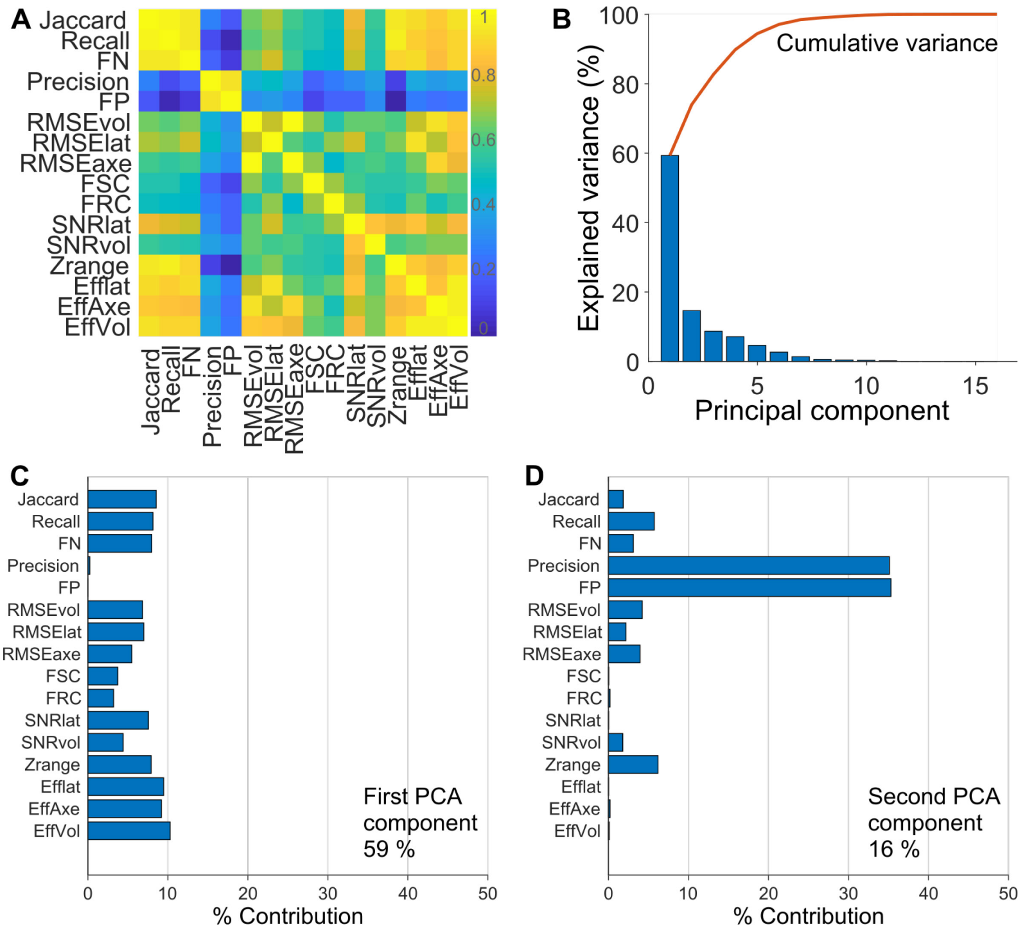


**Figure S16**: *Principal component analysis of 3D assessment metrics.* **A.** Covariance matrix of the 3D metrics for all datasets. **B.** Scree plot showing the percentage of expected variance which can be explained by each principal component. Together, the first two principal components explain 75 % of the expected variance. **C-D.** Percentage contribution of each variable to the first (C) and second (D) principal components.

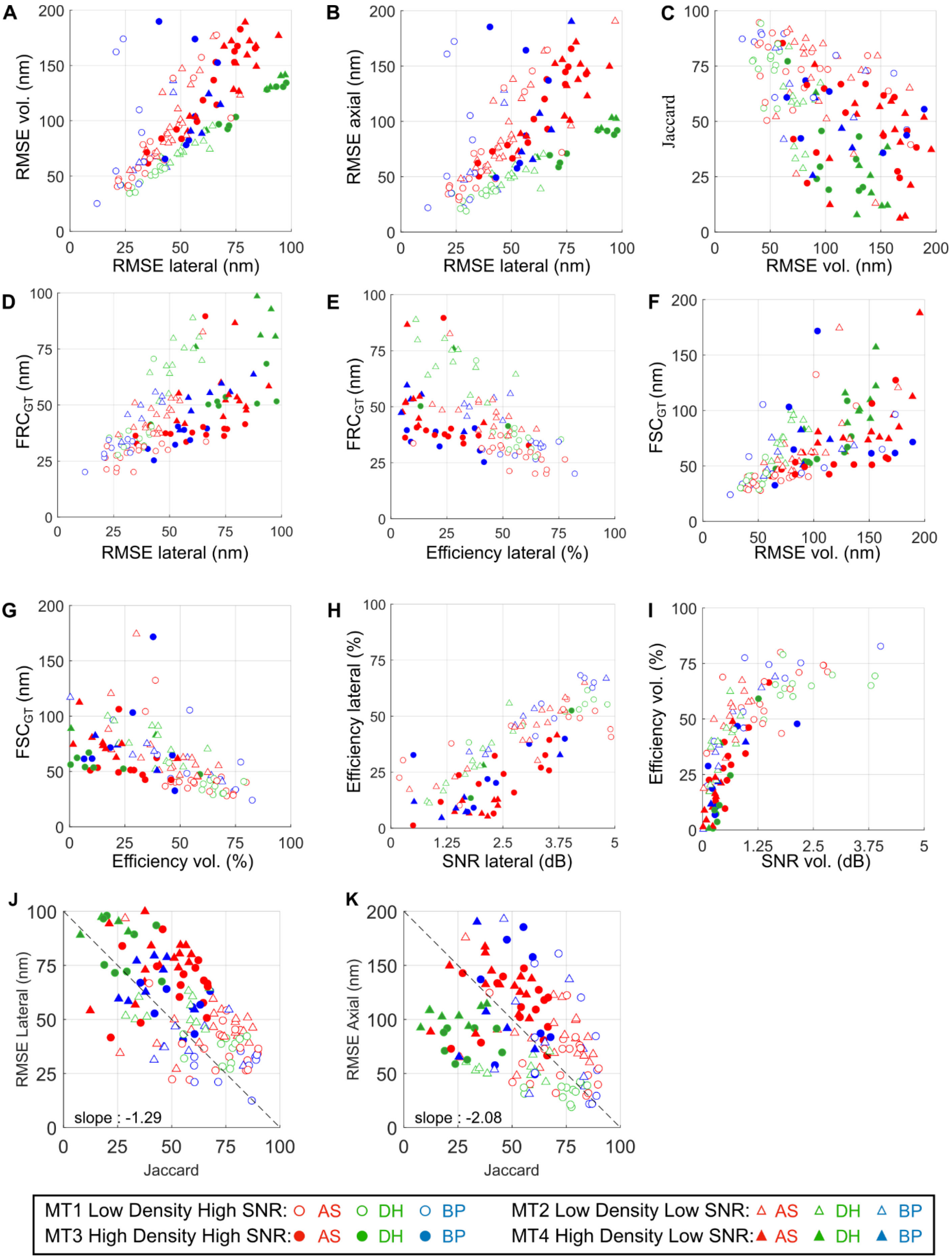


**Figure S17:** Relationships and correlations between the 3D assessment metrics. Data points represent average software performance for a given simulated dataset. Dashed lines in J-K shows the linear regression slope between lateral/ axial RMSE versus Jaccard index for all results (datasets and software). The weighting for the efficiency formulae relies on these observations.

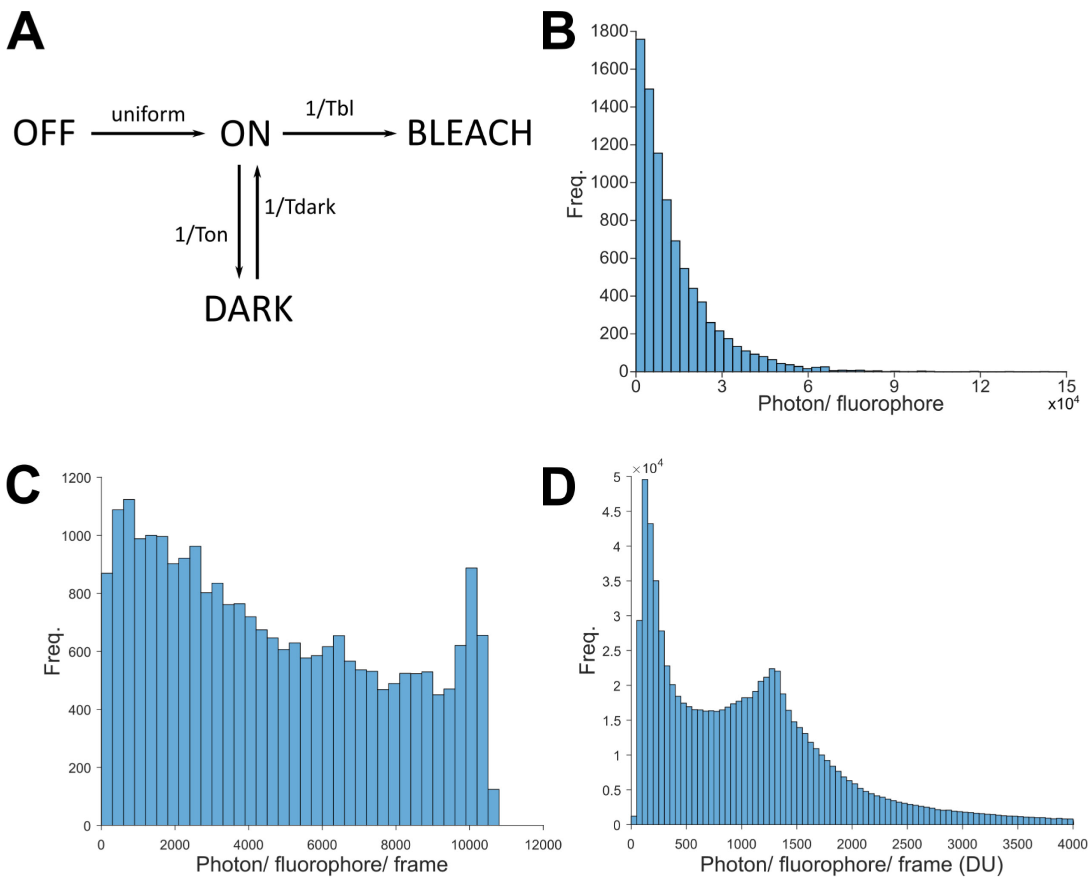


**Figure S18:** *Analysis of the performance of the simulated photophysics model.* **A.** Molecule photophysics was simulated using a 4 state model incorporating inactive, active, dark and photobleached states. **B.** Ground truth photon count distribution per molecule, integrating the photon emission over all bright frames for the MT1.LD.N1 dataset. **C.** Ground truth photon count distribution per molecule per frame, i.e., the distribution of photon counts for individual frames, for the MT1.LD.N1 dataset. **D.** Observed photon count distribution per molecule per frame for real STORM dataset, Tubulin Conj-AL647, from the 2D SMLM software challenge[4], estimated using the ThunderSTORM software.

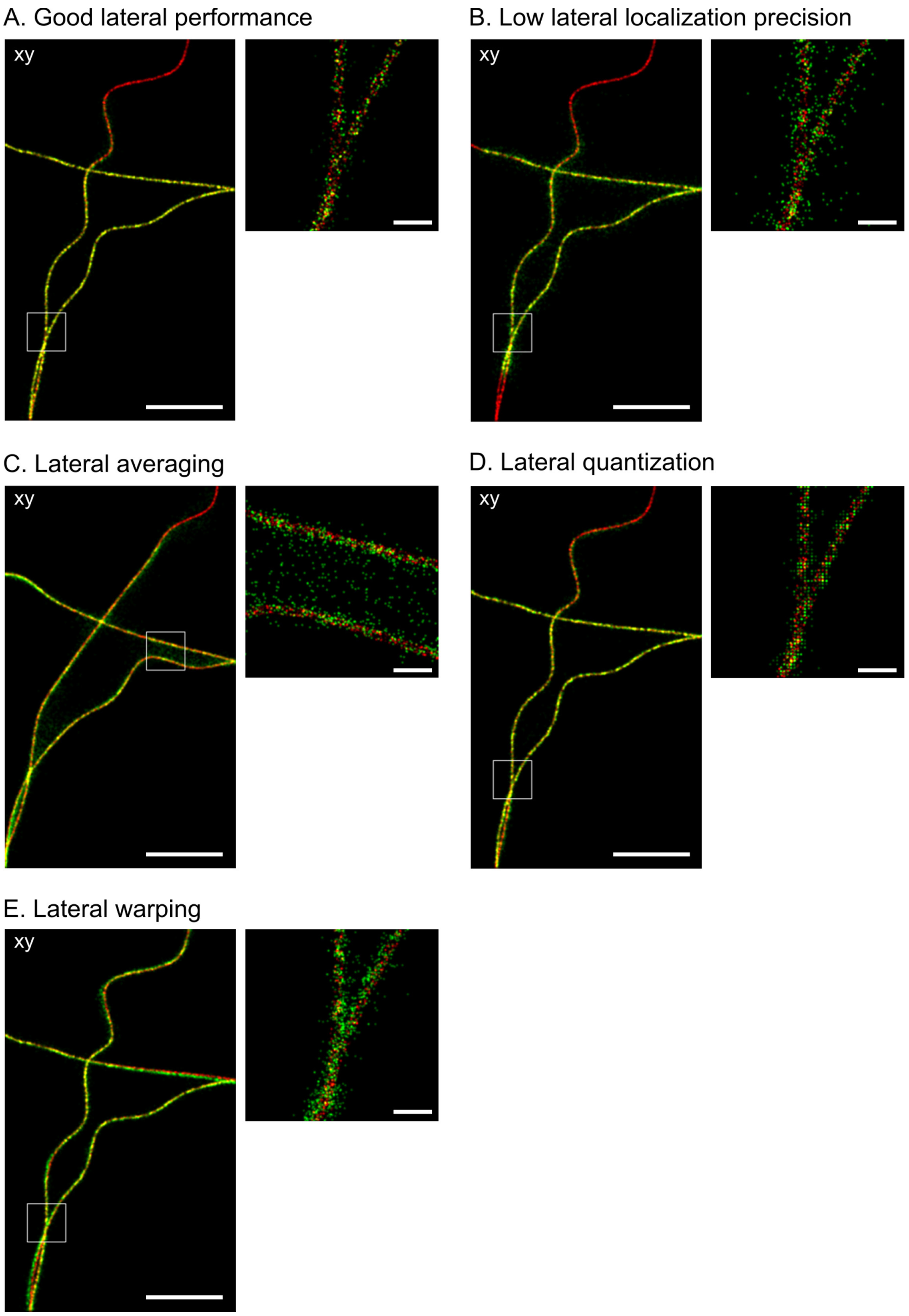


**Figure S19.** *Examples of common SMLM software errors resulting in reduced image quality along the lateral (xy) dimensions*. Smaller panel shows a higher magnification image of the boxed region in the larger panel. **A**. Exemplar good lateral performance. **B.** Low lateral localization precision, probably caused by sub-optimal minimization algorithm. **C.** Lateral averaging, i.e. averaging of the position of two or more bright molecules on a densely labelled structure. **D.** Lateral quantization, i.e., loss of resolution due to localization onto a grid with spacing larger than the localization precision. E. Lateral warping, i.e., lateral deformation of the observed structure with respect to the ground truth structure, possibly due to imperfect wobble correction. The software errors shown here were observed across multiple software packages. *Scale bar*: full field of view, 1 µm , magnified sub-region, 100 nm.

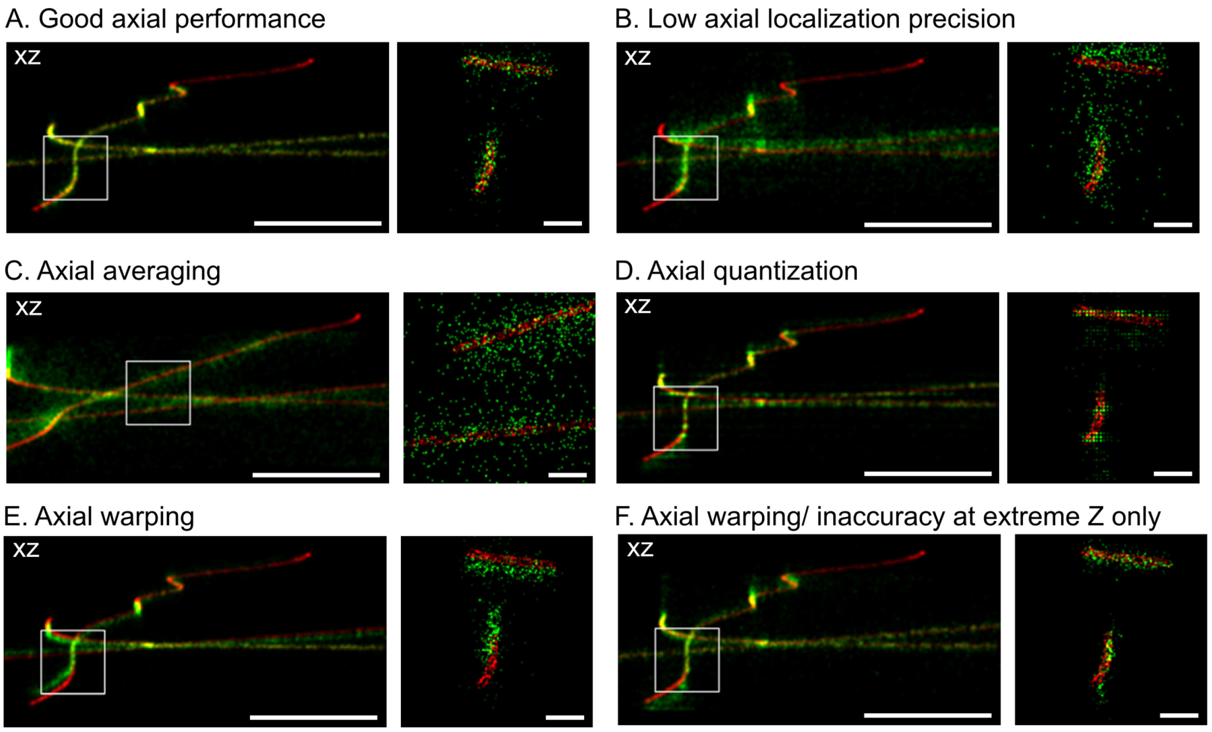


**Figure S20.** *Examples of common SMLM software errors resulting in reduced image quality along the axial (z) dimension*. Smaller panel shows a higher magnification image of the boxed region in the larger panel. **A**. Exemplar good axial performance. **B.** Low axial localization precision, here probably caused by sub-optimal minimization algorithm. **C.** Axial averaging, i.e. averaging of the position of two or more bright molecules on a densely labelled structure. **D.** Axial quantization, i.e., loss of resolution due to localization onto a grid with spacing larger than the localization precision. E. Axial warping, i.e., axial deformation or scaling of the observed structure with respect to the ground truth structure, possibly due to imperfect axial calibration. **F.** Axial warping/ inaccuracy at extreme Z only. Here, some software showed faithful measurement of ground truth axial position near the focus, but showed strong axial deviations away from the focus. The cause of this error is unclear, but could be due to a poor fit between a Gaussian PSF model and the true PSF at large defocus. The software errors shown here were observed across multiple software packages. *Scale bar*: full field of view, 1 µm, zoomed sub-region, 100 nm.

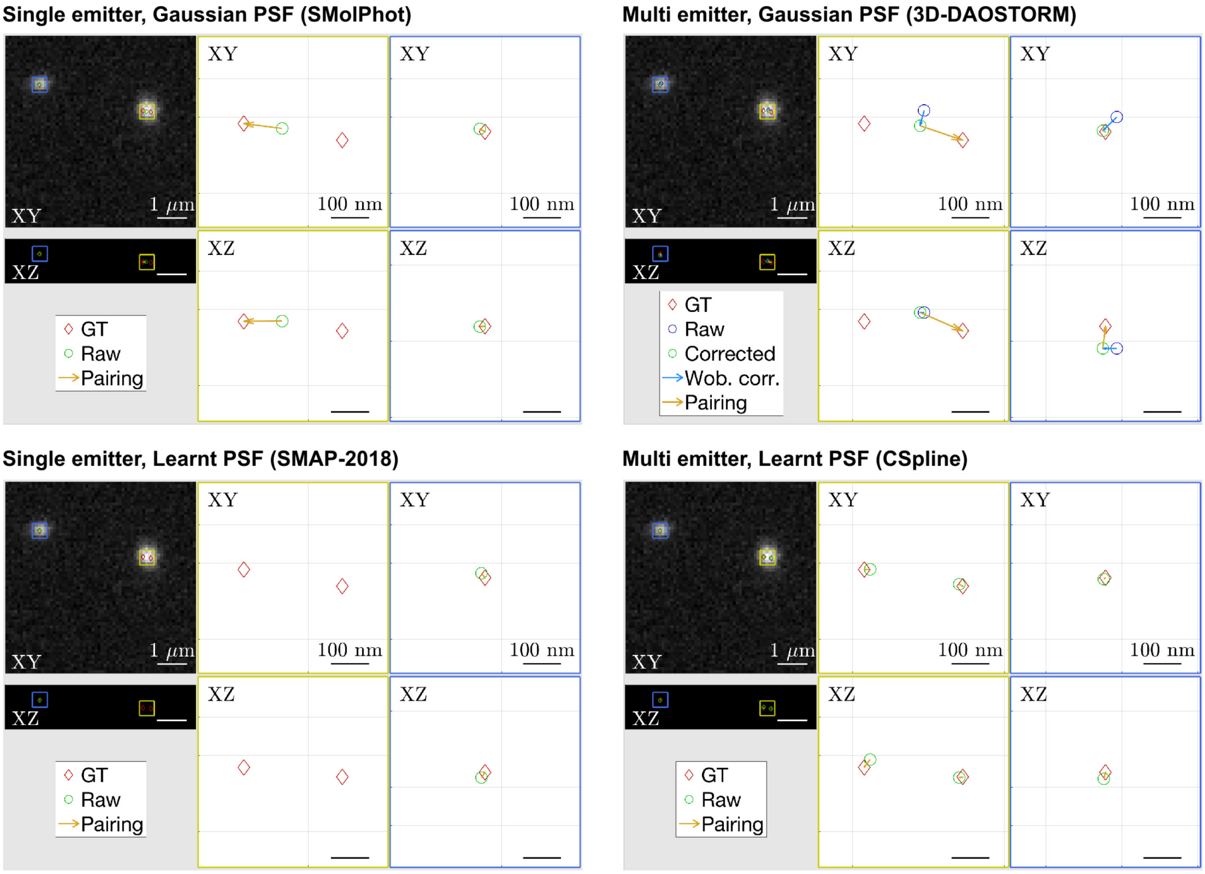


**Figure S21**: *Exemplar single frame performance example*. Examples of software performance for each dataset and software can be viewed in the individual software reports and side-by-side comparison tools on the competition website to help developers identify algorithm and software limitations and improve software performance. Here we give an example of performance of different algorithm classes on the low spot density high SNR astigmatism dataset, MT1.N1.LD AS. The yellow box indicates a region of high spot density, here two overlapping spots**.** The blue box indicates a region of low spot density, here one isolated spot. The raw image is overlaid with the ground truth and observed positions, shown side by side with a magnified plot of the molecule positions in XY and XZ. Yellow arrows indicate the pairing between the ground truth and observed position. In the cases where a wobble correction was applied, the blue circle indicates the raw, uncorrected observed position, the green circle indicates the wobble corrected position, and the blue arrow indicates the shift between the corrected and uncorrected positions. In this example, all software shown accurately localized the isolated spots, but only the multi-emitter learnt PSF software accurately localized both overlapping spots. However, we caution that significant variation in localization accuracy was observed between the different example frames. We believe this tool should primarily be used as a developer diagnostic for the performance of individual software, and that multiple example frames should be viewed before drawing conclusions about the per-frame performance of a given software. General comparisons between multiple software packages should best be drawn robustly using the quantitative analysis metrics measured over multiple frames, using the per-frame examples to help investigate the details of the performance of individual algorithms.

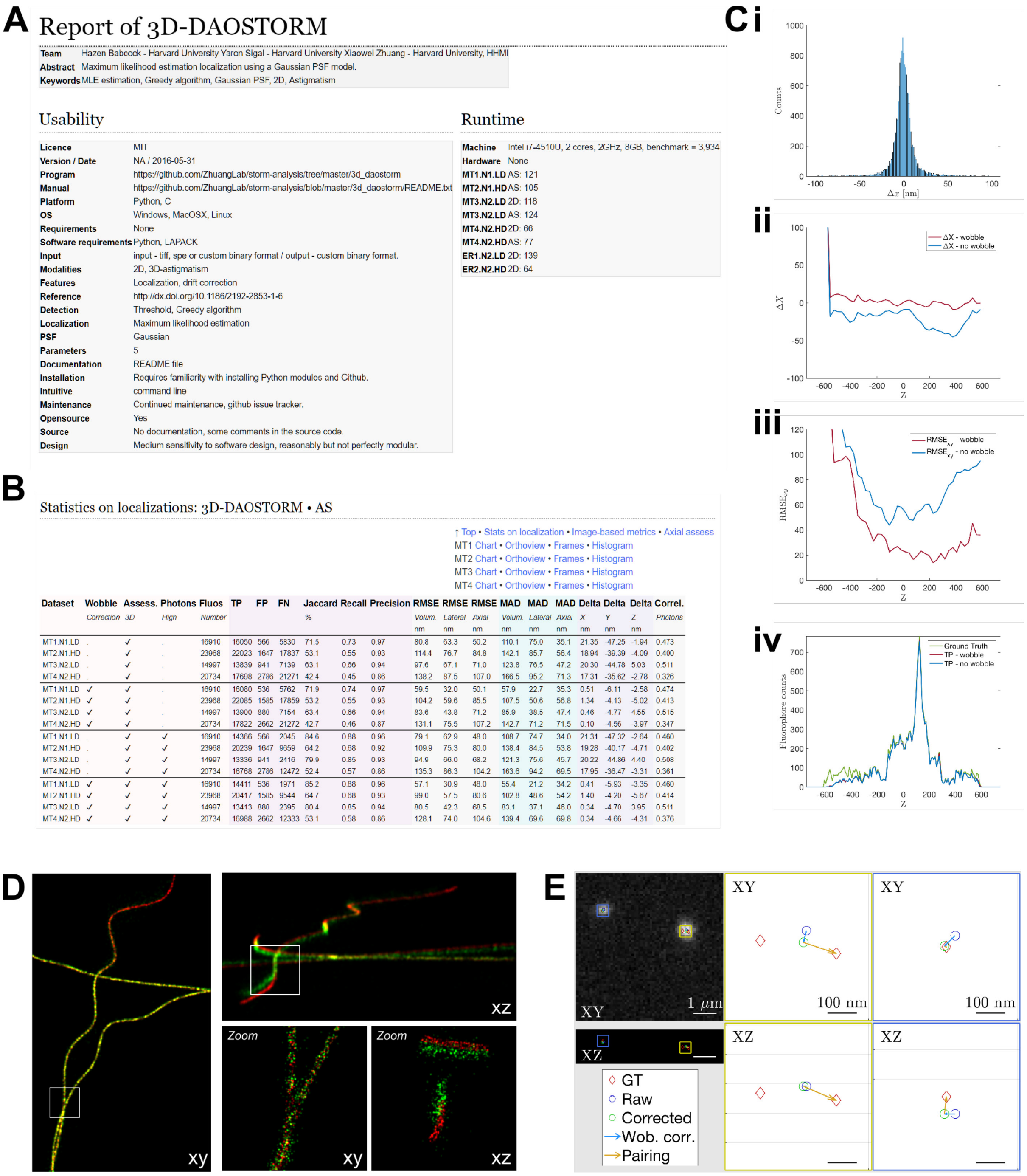


**Figure S22**: *Example software report from the SMLM challenge website.* An extensive performance report is generated for each submission including: (A) information on the software usability, key references, software run time; (B) detailed performance statistics; (C) graphs showing software performance and diagnostic information including bias histograms/ graphs (Ci-ii), and axially resolved localization performance (Ciii-iv); (D) super-resolved overlays of ground truth and software localizations; and (E) per-frame overlays of results and simulated data.

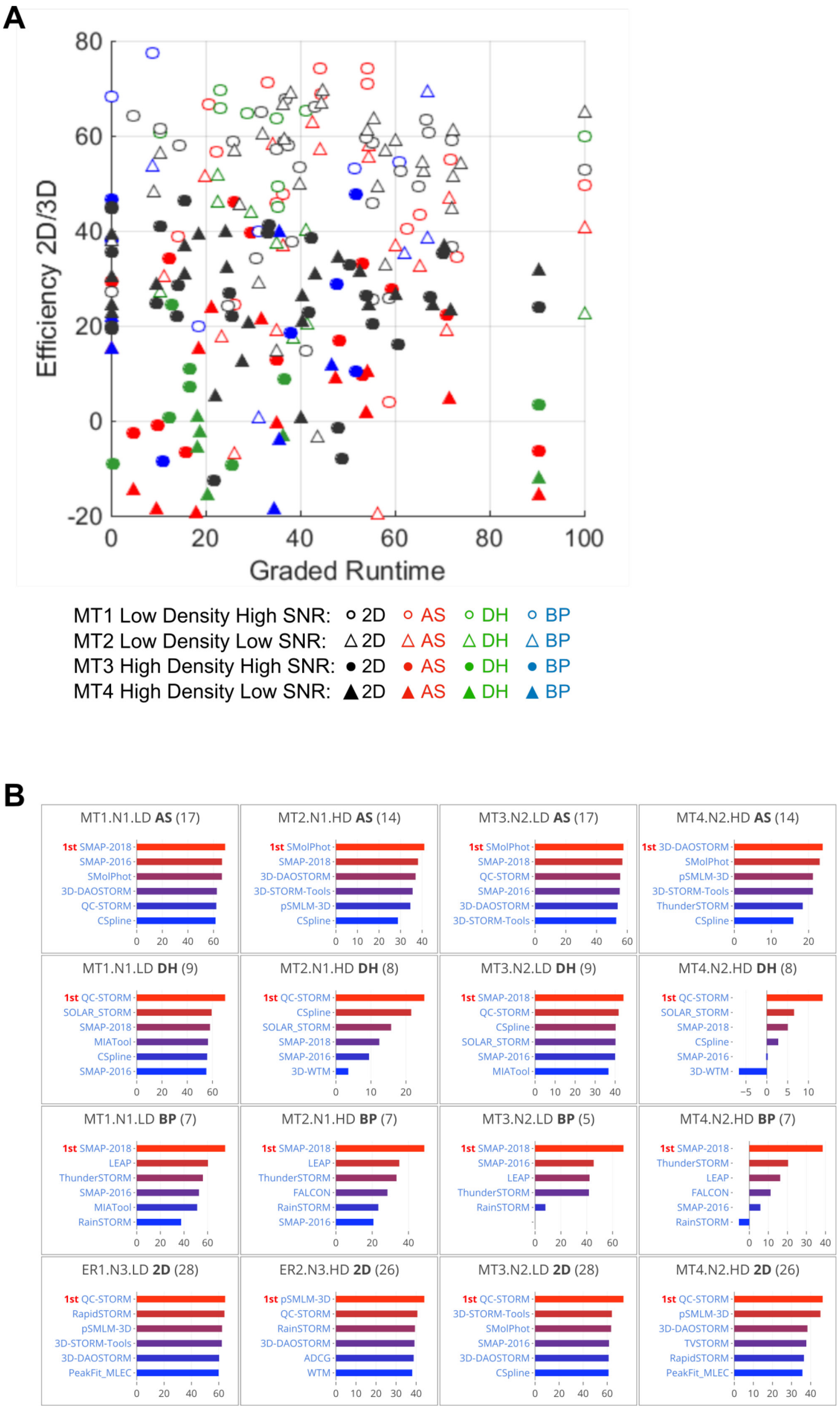


**Figure S23.** *Run time performance analysis.* **A.** Comparison of graded run time against lateral (2D software) or volumetric (3D software) efficiency for all software, datasets and conditions. Graded run time gives a logarithmic score limited between the range [0,100], such that real time software (≤0.01s per frame) gets a score of 100, and very slow software (≥1000s per frame) gets a score of 0. No visible correlation between efficiency and graded runtime was observed. **B.** Software ranking using an alternative Efficiency-Runtime metric which gives 25 % weighting to graded run-time, for each competition modality and dataset.

| Name of the software | 2D | 2D | AS | AS | BP | BP | DH | DH | Wobble | PSF | Temporal grouping | Run-time | Grid | Specific information on the algorithm or the PSF |
|---|---|---|---|---|---|---|---|---|---|---|---|---|---|---|
| | LD | HD | LD | HD | LD | HD | LD | HD | | | | | | |
| Ranked if efficiency larger than | 50 | 30 | 40 | 10 | 40 | 10 | 40 | 0 | | | | | | |
| **Non-iterative** | | | | | | | | | | | | | | |
| 3D-WTM[5] (Commercial software, Hammamatsu) | | | | | | | | | bead | Wedge | No | <1h. GPU | # | Detection, wedge matching |
| QuickPALM[6] | | | | | | | | | bead | Gauss | No | <1min. | C | Center of gravity |
| WTM[5] (Commercial software, Hammamatsu) | | | | | | | | | | Wedge | No | <10min. GPU | # | Detection, wedge matching |
| **Single emitter fitting** | | | | | | | | | | | | | | |
| 3D-STORM-Tools | 7 | | 7 | 4 | | | | | bead | Gauss | No | <10s. GPU | C | LS |
| EasyDHPSF[7] | | | | | | | 6 | | bead | Gauss | No | <3h | C | LS |
| FIRESTORM[8] | | | | | | | | | | Gauss | No | <1min. | C | LS |
| Localizer[9] | | | | | | | | | | Gauss | No | <10s. GPU | C | LS, MLE |
| MaLiang[10] | | | | | | | | | | Gauss | No | n.a. | C | MLE |
| MIATool[11] | 18 | | 9 | | | | 5 | | bead | Bessel | Yes | <10min. | C | MLE |
| mlePALM | | | | | | | | | bead | Gauss | No | <3h. | C | MLE |
| Octane[12] | | | | | | | | | | Gauss | No | <1min. | C | LS |
| PALMER[13] | | | | | | | | | | Gauss | No | <10min. only HD | C | MLE |
| PeakSelector[14] | | | | | | | | | | Gauss | No | <10min. | C | LS |
| QC-STORM | 11 | | 8 | | | | 7 | | auto | Gauss | Yes | <1s. GPU | C | MLE |
| RapidSTORM[15] | 13 | | | | | | | | bead | Gauss | No | <10s. | C | LS, MLE |
| SFP_Estimator[16] | 17 | | | | | | | | | Gauss | No | <1min FGPA | C | MLE |
| SMAP-2016[17] | 6 | | 4 | | 3 | | 2 | 4 | bead | Gauss | Yes | <1min. GPU | C | Several algorithms, LS,MLE |

| | | | | | | | | | | | | | | |
|---|---|---|---|---|---|---|---|---|---|---|---|---|---|---|
| SMAP-2018[17] | 19 | | 2 | 5 | 1 | 1 | 1 | 2 | auto | Learn | Yes | <1min.<br>GPU | C | Cubic spline representation |
| SMolPhot[18] | 3 | | 1 | | | | | | auto | Gauss | Yes | <10min. | C | LS + CLEAN + Frame grouping |
| STORMChaser | | | | | | | | | auto | Gauss | No | <10min. | C | LS |
| WaveTracer[19] (used commercially in Metamorph) | 14 | | | | | | | | auto | Gauss | No | <1h<br>GPU | C | LS |
| **Multi-emitter fitting** | | | | | | | | | | | | | | |
| 3D-DAOSTORM[20] (used commercial by Nikon) | 2 | 3 | 5 | 1 | | | | | bead | Gauss | Yes | <10min. | C | MLE |
| CSpline[21] | 4 | 9 | 3 | 2 | | | 3 | 1 | auto | Learn | Yes | <10min. | C | Cubic spline representation |
| PeakFit[22] | 1 | 4 | | | | | | | | Gauss | No | <10min. | C | LS, MLE |
| RainSTORM[23] | 9 | 8 | | | 5 | | | | bead | Gauss | No | <10min.<br>GPU | C | LS |
| ThunderSTORM[24] | 8 | 5 | | 7 | 4 | 4 | | | bead | Gauss | No | <1min. | C | Several algorithms, LS, MLE |
| **Compressed sensing 1: Matching pursuit algorithms for molecule candidates, sparsity constraint** | | | | | | | | | | | | | | |
| ADCG[25] | 5 | 2 | | | | | | | | Gauss | No | <10min.<br>GPU | C | Constrained gradient descent |
| SMfit | 15 | 5 | | | | | | | | Gauss | No | <3h | C | Block coordinate descent |
| SOLAR_STORM[26] | | | | | | | 4 | 3 | bead | Arbitrary | No | <10min. | C | Orthogonal Matching Purs. + L1H |
| TVSTORM[27] | 12 | 7 | 6 | 3 | | | | | bead | Gauss | No | <1h<br>GPU | C | Backtracking line search |
| **Compressed sensing 2: Deconvolution-type algorithm, sparsity constraint** | | | | | | | | | | | | | | |
| CEL0[28] | | | | | | | | | | Arbitrary | No | <3h<br>only HD | # | Majorization-Minimization (l0) |
| FALCON[29] | 10 | 1 | | | | 3 | | | auto | Gauss | No | <3h<br>GPU | C | Sparsity constraint (weighted l1) |
| L1H[30] | | | | | | | | | | Gauss | No | <10min. | # | L1-homotopy |
| **Other approaches** | | | | | | | | | | | | | | |

| | | | | | | | | | | | | | | |
|---|---|---|---|---|---|---|---|---|---|---|---|---|---|---|
| ALOHA[31] | | | | | | | | | | Gauss | No | | C | Detection, annihilating filter |
| Brecs[32] | | | | | | | | | auto | Arbitrary | No | <1h. GPU | # | Bayesian |
| LEAP[33] | | | | | 2 | 2 | | | auto | Bessel | No | <3h | C | Deconvolution, annihilating filter |
| pSMLM-3D[34] | 16 | 6 | | 6 | | | | | bead | Gauss | No | <10s. | C | Detection, phasor analysis |
| **Participation** | **31** | **28** | **17** | **14** | **9** | **8** | **10** | **9** | | | | | | |

***Table S1.*** **Participant software summary**. *A grey cell in the table indicates the participation to the competition and the number shows the rank of the software for on the 8 categories. The column "Wobble 3D" indicates how the software proceeds with the wobble correction:* ***auto*** *for an intrinsic correction,* ***bead*** *for an explication calibration on beads. The "C" in the column Grid indicates that the software use a grid-less algorithm contrarily to "#" that indicates an algorithm running on a pre-defined grid. The runtime is computed as much as possible by averaging the computation time provided by the participants on the low density datasets (~20'000 frames) for 2D and AS, GPU indicates that the participants have used a graphics processor card.*

| Dataset | Template structure | Sample thickness | SNR | Sample labelling method & fluorophore | Molecule brightness | Autofluorescence level | Molecule density |
| --- | --- | --- | --- | --- | --- | --- | --- |
| | | µm | | | photon/ mol/ frame | photons/pixel/ frame | mol/ µm |
| Beads | Beads calibration sample | 1.5 | Very high | Very bright fluorescent beads | 20000 | 0 | |
| MT1.N1.LD | Microtubules Diameter 25 nm | 1.5 | High | Organic dye (Alexa 647), antibody labelling | 5000 | ~100 (moderate) | 0.25 (low) |
| MT2.N1.HD | | 1.5 | High | Organic dye (Alexa 647), antibody labelling | 5000 | ~100 (moderate) | 2.5 (high) |
| MT3.N2.LD | | 1.5 | Low | Fluorescent protein label (mEos3.2) | 500 | ~10 (low) | 0.25 (low) |
| MT4.N2.HD | | 1.5 | Low | Fluorescent protein label (mEos3.2) | 500 | ~10 (low) | 2.5 (high) |
| ER1.N3.LD | Pseudo endoplasmic reticulum. Diameter 150 nm (approximative) | 0.7 | Medium | Affinity dye label (Mitotracker red) | 3000 | ~500 (high) | 0.5 (low) |
| ER2.N3.HD | | 0.7 | Medium | Affinity dye label (Mitotracker red) | 3000 | ~500 (high) | 5 (very high) |

**Table S*2*: Simulation parameters for each condition.** Molecule density calculated based on approximate diffraction limited area (A) of structure. For a thin linear structure, A = n*L*FWHM where n is the number of filaments, L is the length of filament (approximated as image width), FWHM is PSF FWHM. For a thick linear structure, A=n*L*sqrt(w^2+ FWHM^2)

| Density | Structure | Role | Modality | | | |
|---|---|---|---|---|---|---|
| | | | 2D | AS | DH | BP |
| | Beads | Calibration | Beads | Beads | Beads | Beads |
| Low Density (LD) | MT0 | Training | MT0.N1.LD | MT0.N1.LD | MT0.N1.LD | MT0.N1.LD |
| | ER1 | Contest | ER1.N3.LD | | | |
| | MT1 | Contest | | MT1.N1.LD | MT1.N1.LD | MT1.N1.LD |
| | MT3 | Contest | MT3.N2.LD | MT3.N2.LD | MT3.N2.LD | MT3.N2.LD |
| High Density (HD) | ER2 | Contest | ER2.N3.HD | | | |
| | MT2 | Contest | | MT2.N1.LD | MT2.N1.LD | MT2.N1.LD |
| | MT4 | Contest | MT4.N2.HD | MT4.N2.HD | MT4.N2.HD | MT4.N2.HD |

**Table S3: List of all competition datasets.** Datasets are available at http://bigwww.epfl.ch/smlm/challenge2016/index.html?p=datasets

| Modality | Objective | Camera | Pixel size at the sample plane | Z-step | Z-range |
|---|---|---|---|---|---|
| 2D | Nikon NA 1.49 TIRF oil<br>(commercial N-STORM microscope) | EMCCD | 43 nm<br>(1.5x Optovar and 2.5x SIM magnifiers in place) | 10nm | 3µm |
| Astigmatism (AS) | Nikon NA 1.49 TIRF oil | EMCCD | 160 nm | 10nm | 3µm |
| Double-helix (DH) | Nikon NA 1.49 TIRF oil | EMCCD | 160 nm | 20nm | 3µm |
| Biplane (BP) | The biplane model PSF was constructed from the 2D PSF. The two defocused PSFs were constructed by duplicating the 2D PSF and offsetting it by -250 nm and 250 nm for each Z-plane | | | | |

**Table S*4*:** Microscope acquisition parameters for experimental PSF acquisition for each imaging modality.

| Dataset | Modality | Participants | Winner | Jaccard Average (%) | RMSE Lateral Average (nm) | RMSE Axial Average (nm) | Jaccard Winner (%) | RMSE Lateral Winner (nm) | RMSE Axial Winner (nm) | Z-range winner (nm) | Consolidated Z-range winner (nm) |
|---|---|---|---|---|---|---|---|---|---|---|---|
| MT1.N1.LD | AS | 18 | SMAP-2018 | 73.7 | 38.3 | 66.3 | 84.5 | 26.3 | 29.1 | 1170.0 | 1083.897 |
| MT2.N1.HD | AS | 15 | SMolPhot | 52.7 | 66.3 | 109.3 | 66.3 | 50.5 | 66.3 | 849.0 | 665.946 |
| MT3.N2.LD | AS | 17 | SMolPhot | 69.3 | 44.9 | 84.0 | 85.8 | 39.2 | 60.3 | 855.0 | 817.127 |
| MT4.N2.HD | AS | 15 | SMolPhot | 43.3 | 77.5 | 128.4 | 58.0 | 76.4 | 100.9 | 851.7 | 623.353 |
| MT1.N1.LD | BP | 8 | SMAP-2018 | 75.1 | 27.0 | 90.5 | 87.1 | 12.3 | 21.7 | 1198.6 | 1122.6 |
| MT2.N1.HD | BP | 8 | SMAP-2018 | 54.1 | 55.3 | 116.1 | 60.7 | 43.1 | 48.9 | 1146.8 | 992.657 |
| MT3.N2.LD | BP | 6 | SMAP-2018 | 66.8 | 41.9 | 84.2 | 83.1 | 28.5 | 46.8 | 1048.3 | 941.8 |
| MT4.N2.HD | BP | 7 | SMAP-2018 | 41.9 | 69.3 | 134.7 | 60.5 | 54.6 | 72.3 | 1113.5 | 943.856 |
| MT1.N1.LD | DH | 9 | SMAP-2018 | 72.8 | 36.6 | 34.0 | 77.1 | 27.0 | 20.9 | 1182.5 | 994.762 |
| MT2.N1.HD | DH | 8 | CSpline | 29.1 | 82.9 | 77.9 | 45.6 | 67.4 | 69.2 | 1048.1 | 567.954 |
| MT3.N2.LD | DH | 9 | SMAP-2018 | 49.7 | 53.5 | 55.2 | 65.6 | 45.3 | 42.1 | 1028.3 | 761.962 |
| MT4.N2.HD | DH | 6 | SMAP-2018 | 25.8 | 96.6 | 103.1 | 30.0 | 90.6 | 93.7 | 696.5 | 348.027 |
| ER1.N3.LD | 2D | 32 | ADCG | 72.3 | 35.9 | - | 90.7 | 31.0 | - | - | - |
| ER2.N3.HD | 2D | 28 | ADCG | 56.7 | 52.1 | - | 75.0 | 45.6 | - | - | - |
| MT3.N2.LD | 2D | 32 | SMolPhot | 73.4 | 36.1 | - | 90.1 | 29.0 | - | - | - |
| MT4.N2.HD | 2D | 29 | ADCG | 55.2 | 52.3 | - | 69.9 | 51.0 | - | - | - |

**Table S5:** Best in class and average software performance for each dataset.

### SUPPLEMENTARY NOTES

### 1. METRICS OF ASSESSMENT

We calculated a large number of analysis metrics to quantify the performance of software relative to ground truth. The results of software according to each of the metrics listed here can be interactively explored on the competition website. Software can be ranked individually by any of these metrics, or by arbitrary combinations of these metrics.

#### 1.1 Notation

The notations $(\cdot)^s$ and $(\cdot)^t$ denote the software and ground truth quantities, respectively. Note we distinguish between molecule (*i.e.,* one physical fluorophore) and activation (*i.e.,* one emitting fluorophore during one frame acquisition).

- $x_i, y_i, z_i$: Position of an activation $i$.
- $x^{\mathrm{m}}, y^{\mathrm{m}}, z^{\mathrm{m}}$: True position of molecules.
- Estimated activations (S): Set of estimated activations.
- True activations (T): Set of true activations.
- $\Gamma$ denotes the cardinality of a set.

#### 1.2 Statistics on localizations (frame-wise)

These statistics were calculated based on localizations in individual frames, ie without grouping results on a per molecule basis. Note that although the analysis was conducted on a per-frame basis, analyses which take account of information from multiple frames, such as temporal grouping still gives improved results by increasing the localization precision of the per-frame results.

***Localization based metrics***

1. True positive (TP): $\Gamma(\mathrm{S} \cap \mathrm{T})$. Number of paired activations according to the protocol (*Methods*).
2. False positive (FP): $\Gamma(\mathrm{S} \backslash \mathrm{S} \cap \mathrm{T})$. Number of unpaired estimated activations.
3. False negative (FN): $\Gamma(\mathrm{T} \backslash \mathrm{S} \cap \mathrm{T})$. Number of unpaired true activations.
4. Recall: TP/(TP + FN) [%]. Fraction of true activations detected.
5. Precision: TP/(TP + FP) [%]. Fraction of estimated activations which match true activations (rather than false postitives).
6. Jaccard index (JAC) [%]: TP/(TP + FP + FN). Combines the recall and precision metrics; includes how successfully activations are found, and how well false positives are excluded. High JAC indicates both high recall and high precision. Low JAC indicates both low recall and low precision. Intermediate JAC indicates some combination of low recall and/or low precision.
7. RMSE lateral (RMSE Lateral) [nm]: $\sqrt{\frac{1}{\mathrm{TP}}\sum_{i\in \mathrm{S}\cap\mathrm{T}} (x_i^s - x_i^t)^2 + (y_i^s - y_i^t)^2}$. 2D localization error[4,35,36]. Note: since RMSE lat is defined as a Euclidian distance in 2D space, this metric will be approximately $\sqrt{2}$ larger than an equivalent 1D localization precision used, eg. to calculate whether two adjacent structures are resolvable.
8. RMSE axial (RMSE Axial) [nm]: $\sqrt{\frac{1}{\mathrm{TP}}\sum_{i\in \mathrm{S}\cap\mathrm{T}} (z_i^s - z_i^t)^2}$. Axial localization error.
9. RMSE volumetric (RMSE Vol.) [nm]: $\sqrt{\frac{1}{\mathrm{TP}}\sum_{i\in \mathrm{S}\cap\mathrm{T}} (z_i^s - z_i^t)^2 + (x_i^s - x_i^t)^2 + (y_i^s - y_i^t)^2}$. 3D localization precision. Note: since RMSE lat is defined as a Euclidian distance in 3D space, this metric will be approximately $\sqrt{3}$ larger than an equivalent 1D localization precision.

10. Bias $\Delta x$ [nm]: $\frac{1}{\mathrm{TP}}\sum_{i\in \mathrm{S}\cap \mathrm{T}}(x_i^s - x_i^t)$. Mean X offset in localization. For an unbiased estimator, this shoud be zero. Values much greater than zero indicate a bias in the measurements, typically due to an error in calibration, coordinate system definition, or software bug.
11. Bias $\Delta y$ [nm]: $\frac{1}{\mathrm{TP}}\sum_{i\in \mathrm{S}\cap \mathrm{T}}(y_i^s - y_i^t)$. Mean Y offset in localization.
12. Bias $\Delta z$ [nm]: $\frac{1}{\mathrm{TP}}\sum_{i\in \mathrm{S}\cap \mathrm{T}}(z_i^s - z_i^t)$. Mean Z offset in localization.
13. Photons correlation (Correl. photons): Pearson correlation $r = \frac{\sum_{i\in \mathrm{S}\cap \mathrm{T}}(p_i^s - \bar{p}^s)(p_i^t - \bar{p}^t)}{\sqrt{\sum_{i\in \mathrm{S}\cap \mathrm{T}}(p_i^s - \bar{p}^s)^2}\sqrt{\sum_{i\in \mathrm{S}\cap \mathrm{T}}(p_i^t - \bar{p}^t)^2}}$ with $p$ the photon count and $\bar{\cdot}$ the averaged value. Compares the correlation between observed and ground truth photon counts. Estimates how good software is at estimating relative photon counts.

***Image-based metrics***

14. Fourier ring correlation to ground truth ($\mathrm{FRC_{GT}}$) [nm]: $\mathrm{FRC_{GT}}$ for projected volume along the Z axis (on lateral plane XY). FRC[37] for experimental data is a relative estimator of lateral resolution, measured by randomly dividing the localizations into two subsets, rendering super-resolved images of the data, performing spatial cross-correlation of the two images, and then estimated the distance at which cross-correlation drops below a critical value.
The $\mathrm{FRC_{GT}}$ is similar, except that here we exploit the availabilty of the ground truth information by performing the FRC calculation on the correlation between the rendered estimated activations and the rendered true activations. Thus the $\mathrm{FRC_{GT}}$ reports not only on spatial resolution, but also any systematic errors or biases in the estimated activations compared to the ground truth data. For example a large $\Delta x$ would lead to a large $\mathrm{FRC_{GT}}$xy.
Note that FRC and $\mathrm{FRC_{GT}}$ are only relative, rather than absolute measures of resolution[38].
15. Fourier shell correlation to ground truth ($\mathrm{FSC_{GT}}$,) [nm]: Fourier Shell Correlation[37] is similar to the FRC, except the calculations are performed in 3D rather than 2D, so it estimates 3D resolution. Like $\mathrm{FRC_{GT}}$, $\mathrm{FSC_{GT}}$ is performed by cross-correlation of the estimated and true activation datasets. $\mathrm{FSC_{GT}}$ is a relative estimator of volumetric resolution.
16. Signal-to-noise ratio lateral (SNR lat) [dB]: SNR on the plane XY defined as $20\log\left(\frac{I^t}{I^t - I^s}\right)$ with $I$ the Gaussian rendered image ($\sigma = 5$nm). The 2D images $I$ are the results of the projection of all localizations on the plane XY.
17. Signal-to-noise ratio volumetric (SNR vol) [dB]: SNR on the volume defined as $20\log\left(\frac{I^t}{I^t - I^s}\right)$ with $I$ the Gaussian rendered image ($\sigma = 5$nm). The volume has $1500 \times 1500 \times 750$ voxels.

#### 1.3 Axial (depth) assessment

We propose new metrics related to the axial (depth) performance of the software. These metrics quatify the Z-range in which good software performance was obtained. These metrics are extracted from curves as shown in Fig S12. From the curve showing a metric with respect to the axial position, we assess the depth performance of each software. These metrics with respect to the axial position are “locally" computed for each axial position (i.e., small interval of axial positions). We fit a parabolic curve to each curve in order to reduce the effect of finite numbers per bin.

18. R(z): Recall with respect to axial position.
19. $\mathrm{R}_p(z)$: Parabolic fit of R(z).
20. Max recall: Maximal recall of $\mathrm{R}_p(z)$.
21. MinZ recall [nm]: The lowest axial position for which $\mathrm{R}_p(z)$ is below the half of the maximum of $\mathrm{R}_p(z)$.

22. MaxZ recall [nm]: The highest axial position for which $\mathrm{R}_p(z)$ is below the half of the maximum of $\mathrm{R}_p(z)$.
23. $\mathrm{FWHM}_Z$ recall [nm]: Full-width half-maximum of $\mathrm{R}_p(z)$, $\mathrm{FWHM}_Z$ = MaxZ recall - MinZ recall.
24. Consolidated Z-range [nm]: To highlight the depth performance in one unified metric, we design a new metric from the product of $\mathrm{FWHM}_Z$ with Max recall. We call this metric Consolidated Z-range (Fig S13).

#### 1.4 User reported parameters

A large number of user reported parameters were collected for each software. These are available on the challenge results page for each software, providing detailed information to help users choose between different software. An example software report is included in Fig. S22.

The usability report contains information on user-friendliness, installation, whether the software is open-source, and whether the software is actively maintained. Key references are listed. Run-time information for each of the competition datasets is also included.

#### 1.5 Quantitative parameters:

25. Run time per frame (s): Participant-reported total run time for each dataset on stated hardware, divided by the total number of frames.
26. Graded run time:
    $RT_{norm} = 100 - 20 * \min(5, \log_{10}(\mathrm{T} * 100))$ .
    $\mathrm{RT}_{\mathrm{norm}}$ is defined s.t.:
    - a software that runs in 0.01 s / frame has a grade of 100 (real time with a camera at 100Hz)
    - a software that runs in 1 s / frame has a grade of 60
    - a software that runs in 100 s / frame has a grade of 20
    - a software that runs more than 1000 s / frame has a grade of 0 (difficult to use)

#### 1.6 Choice of primary analysis metrics

The metrics listed above can all be interactively explored on the website leader board, which allows ranking by each metric, 2D plotting of different metrics, and construction of new metrics by combination of the existing metrics (Fig S11). Users can easily rank software by their own preferred metric or combination of metrics directly on the competition website.

In order to analyse software performance, it was necessary to identify primary metrics which could summarize the data. From the above metrics, we excluded *bias* (10-12), which indicates software errors and is a quality control metric rather than a ranking metric and *photon correlation* (13) is useful for assessing photometry rather than localization.  We chose *Consolidated Z-range (24)* as the primary axial analysis metric based on considerations discussed in *Methods 3.2*.

This left us with the 15 localization performance metrics plus a software run time metric: *Jaccard, Recall, FN, Precision, FP, RMSEvol, RMSElat, RMSEax,* $FSC_{GT}$*,* $FRC_{GT}$*, SNRlat, SNRvol, Zrange.*

In order to rank overall software performance, we performed a principal component analysis of core metrics to identify key variables (Fig S16). We found that 75 % of the expected variance could be explained by the first two principal components (Fig S16B). 59 % of the data variation could be described by the first principal component (Fig S16C), with most metrics having similar weighting in this component, indicating a high degree of redundancy in the metrics. Most metrics also showed clearly visible correlation (Fig S17).

Precision and FP showed low correlation to the other metrics (Fig S16A) and a strong contribution to the second principal component (Fig 16D, PCA2 corresponded to 16 % of the data variation). On inspection of the data we found that this was because the number of false positives in the data was quite low, ie FP and precision showed low variation across the dataset, unlike the other metrics, and were therefore relatively uninformative metrics for these data.

Since most metrics showed significant redundancy, whichever metric we choose for software ranking should give fairly similar ranking. We therefore made the pragmatic choice to focus our main analysis on localization-based metrics, i.e., the RMSE, Jaccard index, Recall, etc. This is because these metrics have clear physical meaning such as absolute localization precision or fraction of correctly detected molecules. By comparison, $FSC_{GT}$/ $FRC_{GT}$ and other image based metrics provide relative information on resolution which are affected by factors such as sample geometry and fluorophore photophysics[38]. However, alternative analyses using $FSC_{GT}$/ $FRC_{GT}$ can be easily be performed directly on the competition website (Fig S11).

#### 1.7 Software ranking metrics

It was also necessary to choose single metrics by which to competitively rank software. The most important criteria for software performance will inevitably depend on the specific requirements of the user and experiment. Therefore we have created an interactive ranking tool which allows choice of different ranking metrics, including arbitrary user-defined metrics (Fig S11).

We describe below several common performance criteria for SMLM, and the ranking metrics which address these criteria.

- *Sampling limited SMLM (Efficiency)*

In the most common PALM/ STORM/ dSTORM modalities, molecules are photoactivated and imaged only a few times before irreversible photobleaching. Therefore, it is important to both localize molecules accurately and find molecules efficiently, because spatial resolution depends on both localization precision and sampling[38].

In order to provide a single ranking of software performance, we combined the metrics which estimate ability to detect molecules (Jaccard index, Precision, etc), with the metrics which estimate localization precision (localization RMSE ) into a single metric which we term the *Efficiency*,

$$E_{lat} = 100 - \sqrt{(100 - JAC)^2 + \alpha_{lat}^2 RMSE_{lat}^2}$$

$$E_{ax} = 100 - \sqrt{(100 - JAC)^2 + \alpha_{ax}^2 RMSE_{ax}^2}$$

$$E_{3D} = (E_{lat} + E_{ax})/2$$

The trade-off between these two metrics is controlled by a parameter α. In a retrospective analysis, we chose $\alpha = 1\ nm^{-1}$ for the lateral efficiency $E_{lat}$, $\alpha = 0.5\ nm^{-1}$ for the axial efficiency $E_{ax}$, based on the linear regression slope between the localization errors and Jaccard index (Fig 17J-K). Using this definition, an average software performance has an efficiency in the range 25-75, ground-truth has the maximum efficiency of 100. Overall 3D efficiency was calculated as the average of lateral and axial efficiencies.

Since sampling limited SMLM is probably the most common SMLM use case, we used efficiency as the primary software ranking tool.

- *Fast sampling limited SMLM (Efficiency-Runtime)*

SMLM analysis often needs to be not only efficient, but also fast. This is either to enable real time live cell SMLM, or simply for convenient user interaction and data analysis. We generated a combined

*Efficiency-Runtime* metric, which gives 75 % weighting to Efficiency defined previously, and 25 % to the graded run time. This metric is defined as :

$$EffRT = 0.75 * E + 0.25 * RT_{norm},$$

Where $E$ is efficiency, $RT_{norm}$ is the graded software run time (metric 26)

An alternative *Efficiency-Runtime*-based ranking is presented in Fig S23 and the results are discussed in the main text. The Efficiency-Runtime ranking can also be analysed on the competition website.

- *Localization precision limited SMLM (RMSE)*

Several modalities of SMLM, especially PAINT[39] and DNA-PAINT[40] allow relocalization of labelled molecules/ structures for arbitrary periods. In this case it is more important for software to accurately localize molecules than to efficiently find molecules. Therefore a more appropriate ranking criterion for PAINT is RMSE only. Software can be ranked by RMSE on the competition website.

- *3D localization over a small z-range ($RMSE_{in\text{-}focus}$)*

Sometimes, 3D information is only required over a small Z-range, for instance at the surface of a cell membrane which is bound to a microscope coverslip. In this case software performance over the *best-focus* 100 nm of the competition dataset can be analysed, using either $RMSE_{in\text{-}focus}$ or $E_{in\text{-}focus}$ (efficiency, defined above), on the competition website.

- *Structure-based SMLM analysis ($FSC_{GT}$/ $FRC_{GT}$)*

SMLM can display sensitive to structure related artefacts. For example where line-like microtubules run close together in parallel, at very high spot density or small lineseparation SMLM software will return a single averaged line[41]. Metrics which quantify similarity of localizations to the underlying simulated structure, rather than to individual ground truth localizations, are reported to be more robust to such effects[36]. The $FRC_{GT}$/ $FSC_{GT}$ metrics, which correlate the observed localization image with the ground truth structure, fit the criteria of such a structure-based metric. These metrics showed strong visible correlation to the localization-based metrics used in the primary analysis (Fig S17F-I) suggesting that structure related artefacts were not a significant problem in the competition datasets. However, $FRC_{GT}$/ $FSC_{GT}$-based analysis, or other structure-based metrics, could be useful in future for cases where structure-related artefacts are an issue. Software can be ranked by $FRC_{GT}$/ $FSC_{GT}$ on the competition website.

- *Arbitrary user-defined metrics*

Software can be ranked by any user-defined combination of the competition metrics directly on the competition website.

This and the other interactive data exploration tools on the competition website were implemented using the Plotly.js graphing library.

### 2 PHOTOPHYSICS SIMULATION

Fluorophore blinking was simulated by a 4-states Markov chain model (Methods 2.2), with states ON, OFF, BLEACH, DARK (Fig S18A) designed to model photoactivatable fluorescent protein photophysics[42] or reversible organic dye photoswitching[43]. All state transitions are Poisson distributed except the OFF to ON transitions which follow a uniform random distribution.

We analyzed the resulting simulated fluorophore lifetime distributions. As expected, the total photon counts per molecule showed exponential decay (Fig 18B). When we plotted the ground truth per-frame photon count distribution of the molecules for the simulated model, we observed a more complex distribution, showing a decaying distribution with a large peak at zero photons, and a strong second peak, followed by a rapid decay (Fig 18C). This arises from dividing continuous photon emission into discrete frame intervals. The secondary peak and subsequent decay correspond to frames where a molecule is in a bright state for the whole frame. The slowly decaying zero-peak corresponds to molecules which bleach or turn off at any point in the middle of a frame.

We then analyzed the observed the photon count distribution of experimental SMLM data. We used ThunderSTORM[24] to analyze an experimental STORM dataset, Tubulin Conj-AL647, from the 2D SMLM software challenge[4] (Fig S18D). These data were obtained for microtubules labelled with Alexa-467 antibody conjugates[44]. We observed a similar experimental photon count distribution to that of the simulated 4 state dataset (Fig S18C-D). For the current challenge, we thus concluded that the 4 state photophysics model was sufficiently realistic, and used this model for the challenge simulations.

However, we also observed that in the experimental data, the tail in the photon count distribution after the second peak decayed more gradually than for the simulated data (Fig S18C-D). We hypothesize that this is due to additional experimental complexity in the photon count distribution. The simulated fluorophores have uniform quantum yield, extinction coefficient and brightness distribution, which is then used as the input for simulation stochastic Poisson process. By contrast, the experimental situation may be more complex due to multiple effects which will broaden the fluorophore brightness distribution. These include sample-interaction-induced variations the fluorophore nanoenvironment[45], axial-position-related inaccuracies in photon counting[46] and non-uniform sample illumination. In future it would be interesting to extend the 4 state photophysics model with a Gaussian distribution of fluorophore brightness, with a standard deviation directly derived from experimental data.

### 3 TEMPORAL GROUPING

Temporal grouping is a common post-processing approach to improve localization precision by exploiting the assumption that fluorescently labelled molecules in fixed cells should be immobile. Temporal grouping is a post-localization step which identifies molecules active across multiple adjacent frames, and averages their position[47]. Although grouping has been used since the first SMLM papers, grouping has been discussed relatively little in the literature in the context of optimizing image resolution and image quality. Rather, grouping has primarily been discussed in the context of molecule counting/ cluster analysis[47–49].

One drawback of the most common temporal grouping algorithm, “reductive” grouping, is that it deletes all of the individual per-frame localizations and replaces them with a single averaged localization. Although this increases localization precision, and is the first step towards accurate counting of molecules, it artificially increases the amplitude of spurious short on-time background localizations compared to real structures.

Interestingly, the constraints of the SMLM challenge highlighted a little-described approach to temporal grouping which has beneficial features. Because localization performance in the competition was assessed on a per-frame basis, it was not possible to use the reductive grouping algorithm, as this would prevent localizations from being matched to the appropriate frames. The simple alternative, employed by multiple competitors, is “replacement” grouping, where each localization within a group is replaced with the average molecule position, i.e. the total number of localizations remains the same. This approach resolves the spurious background amplification problem of reductive grouping and may be more appropriate where accurate molecule counting is not required. For fixed cells, there does not appear to be any obvious disadvantage of replacement grouping, only an increase in localization precision due to incorporation of temporal information.

### 4 CRAMER-RAO LOWER BOUND & RMSE ANALYSIS

The Cramér-Rao lower bounds (CRLBs) were computed based on the mathematical expression derived by Chao and coworkers[50,51]. The PSFs are 10 nm spaced from which a cubic spline representation is computed over an area of 6.4 x 6.4 µm$^2$. Contrarily to Chao et al. the PSFs were not regularized. As shown in Fig 1, our acquisition model follows that of the challenge simulations (Online Methods) and is similar to the one described by Chao et al. The point source with N emitted photons is first convolved with the PSF and integrated over each camera pixel. The background B is added to each pixel value (except the BP modality for which we added B/2 to each pixel) and these quantities are then converted to an electron count by multiplying them with the Quantum Efficiency (QE = 0.9). Some spurious charge coming from the EMCCD camera (c =0.0002) is added to this quantity before applying a Poisson shot noise $n_{ie}$. The noise from the EMCCD camera (gain g = 300) is modelled as a Gamma distribution with the shape and scale parameters k = $n_{ie}$ and θ = g respectively. Finally, a zero-mean Gaussian readout-noise (σ = 74.4) is added. The analog-to-digital conversion is negligible and ignored for the computation of the CRLB. Once the Fisher Information matrix is obtained[51], the square root of the diagonal of its inverse equals the CRLB. This setting corresponds to the MT1.N1.LD dataset for a individual frames, ie without temporal grouping (see Fig S8CD for MT3.N2.LD).

We assessed the evolution of the CRLB as a function of the depth Z (Fig S8A,C), or as a function of the number of emitted photons N per frame (Fig S8B,D). The CRLB was compared to the 1D RMSE performance in x, y, and z, of the best-in-class software of each 3D modality for the low spot density datasets, at high and low SNR (Fig S8). Note that in case of equal RMSE in x and y directions, the lateral 2D RMSE (Euclidian distance) used as a competition assessment metric would be approximately √2 larger than the 1D RMSE.

We observed that software RMSEs for biplane and double helix were consistently superior to the CRLBs. We hypothesized that this was because all best-in-class software used temporal grouping, i.e. incorporated information from multiple frames

In order to account for the temporal grouping, we multiplied the Fisher information matrix for a single frame by the number of frames on which a molecule is active on average (*i.e.,* 2.85 frames). Then, the square root of the diagonal of its inverse equals the CRLBs (Fig S9). When we compared the software RMSE to the CRLB taking into account multiple frames activations/ temporal grouping, we observed that best-in-class biplane and double helix software closely matched the CRLB limit. Interestingly best-in-class astigmatism software RMSE was 25-50 % worse than the CRLB limit (Fig S9). We speculate that at least away from the focus, where the deviation is largest, this could be due to the difficulty in detecting and minimizing fits to very defocussed, optically complex astigmatic PSFs.

#### 5 LIST OF PARTICIPANTS TO THE CHALLENGE 2016

| 3D-DAOSTORM | |
|---|---|
| Contact | Hazen Babcock, Harvard University (Zhuang group), Cambridge, MA, USA |
| Reference | Babcock, H.P. and Zhuang, X., 2017. Analyzing single molecule localization microscopy data using cubic splines. *Scientific Reports*, *7*(1), p.552. |
| Open-access | https://github.com/ZhuangLab/storm-analysis |
| Platform | Python |
| Class of algorithms | Multi-emitter fitting algorithm |
| Participation 2016 | 2D, AS |
| Note of the authors | Maximum likelihood estimation localization using a Gaussian PSF model. |

| 3D-STORM Tools | |
|---|---|
| Contact | Fabian Hauser, CURT research group, University of Upper Austria, Linz, Austria |
| Open-access | Proprietary |
| Platform | Qt framework |
| Class of algorithms | Single emitter fitting |
| Participation 2016 | 2D, AS |
| Note of the authors | The 3D STORM Tools are easy to use applications for 3D astigmatism super-resolution microscopy. Each tool handles another domain of the 3D STORM method. The Calibration3D tool determines the calibration curves for the further steps. The Analysis3D tool evaluates 3D STORM experiments and returns a list of localizations. The Visualization3D tool offers the possibility for further 3D data analysis (*e.g.,* clustering) and illustrates localizations of fluorophores in a 3D plot and as 2D heat-map image. All tools are written in C++ using the Qt framework version 5.7. |

| 3D-WTM | |
|---|---|
| Contact | Shigeo Watanabe, Hamamatsu Photonics K.K., Japan |
| Reference | Takeshima, T., Takahashi, T., Yamashita, J., Okada, Y. & Watanabe, S. A multi-emitter fitting algorithm for potential live cell super-resolution imaging over a wide range of molecular densities, J. Microsc. 2018 |
| Open-access | Proprietary |
| Platform | Stand-alone (C++) |
| Class of algorithms | Non-iterative (Template Matching) |
| Participation 2016 | 2D, AS, DH, BP |
| Note of the authors | Previously we developed the Wedged Template Matching (WTM) algorithm for localizing molecules with overlapping emission point spread functions in 2D. Here we developed 3D-WTM by extending the function of WTM to 3D localization. 3D-WTM prepares template catalogue from beads z-stack images for each modality, 2D, astigmatism, double-helix and Biplane. By using the different central angle wedged shape template depending on the pixel intensity 3-WTM localizes each molecule in 3D space. After background subtraction, template matching is applied for z-axis and then sub-pixel level template matching for lateral x, y-axis localization, using normalized cross-correlation as matching evaluation function. WTM keeps applying template matching to all the information until all the molecules at images are recognized. . |

| ADCG | |
|---|---|
| Contact | Nicholas Boyd, Geoff Schiebinger, Ben Recht, University of Berkeley, USA |
| Reference | N. Boyd, G. Schiebinger, and B. Recht. “The alternating descent conditional gradient method for sparse inverse problems.” In: SIAM Journal on Optimization 27.2 (2017), pp. 616–639. |
| Open-access | https://github.com/nboyd/SparseInverseProblems.jl |
| Platform | Julia |
| Class of algorithms | Matching pursuit algorithm for molecule candidates, sparsity constraint |
| Participation 2016 | 2D |
| Note of the authors | Simple localization using the alternating descent conditional gradient method to fit Gaussians. |

| ALOHA | |
|---|---|
| Contact | Junhong Min and Kyong Jin, Bio-Imaging and Signal Processing Lab, KAIST, Korea |
| Reference | Min, J., Carlini, L., Unser, M., Manley, S. & Ye, J. C. Fast live cell imaging at nanometer scale using annihilating filter based low rank Hankel matrix approach. Wavelets and Sparsity (2015). |
| Platform | Matlab |
| Class of algorithms | Annihilating filter |
| Participation 2016 | 2D |
| Note of the authors | This is a grid-free 2D localization algorithm with data-driven PSF estimation. Specifically, based on the observation that the sparsity in the spatial domain implies the low-rankness in the Fourier domain, the proposed method converts PSF estimation as well as source localization problems to Fourier-domain signal processing problems so that a truly grid-free localization is possible with adaptive PSF estimation. |

| Brecs | |
|---|---|
| Contact | Hervé Rouault, Timothée Lionnet, Janelia Research Campus, HHMI, VA, USA |
| Open-access | https://github.com/hrouault/Brecs |
| Platform | C (Gui for ImageJ / Fiji) |
| Class of algorithms | Bayesian approach |
| Participation 2016 | 2D, BP, DH |
| Note of the authors | Several recent methods have proposed to reconstruct images at higher densities of fluorophores. However, these heuristic-based techniques are usually constrained to specific imaging schemes and are limited in their performance. We propose a rigorous formulation of the inference problem to solve, and use advanced statistical inference techniques (the Bethe approximation) to provide a controlled approximation to the exact solution. Our method, which we called B-recs (Bethe reconstruction), achieves excellent performance especially in the case of dense samples. Importantly, our technique is versatile thanks to its general and rigorous framework. We demonstrate its use with examples covering various leading edge imaging modalities, ranging from 2D superresolution to various modalities of high-density 3D imaging. A user-friendly version of our algorithm is freely available |

| | |
|---|---|
| | as a plugin for the popular open-source Image Analysis Software Fiji. A unique feature of B-recs is that it provides two complementary estimators for the reconstructed image: a discrete one, listing the coordinates of each fluorescent molecule detected (this is the standard output of LM algorithms); a probabilistic one, which consists of an image in which molecules appear as probability clouds which size and shape encode the localization uncertainty of each localized spot. |

| CEL0 | |
|---|---|
| Contact | Emmanuel Soubies, INRIA Sophia Antipolis, France |
| Reference | Soubies, E., Blanc-Féraud, L. & Aubert, G. A Continuous Exact CEL0 Penalty (CEL0) for Least Squares Regularized Problem. SIAM J. Imaging Sci. 8, 1607–1639 (2015). |
| Platform | Matlab |
| Class of algorithms | Deconvolution-type algorithms with sparsity constraint |
| Participation 2017 | 2D (high-density) |
| Note of the authors | CEL0-STORM is an algorithm designed for 2D high-density molecule localization. We formulate the localization problem as a sparse approximation problem which is then relaxed using the recently proposed CEL0 penalty. This relaxation is then minimized with a nonsmooth nonconvex algorithm, namely the Iterative Reweighed L1 (IRL1) algorithm. |

| CSpline | |
|---|---|
| Contact | Hazen Babcock, Harvard University (Zhuang group), Cambridge, MA, USA |
| Reference | Babcock, H. P. & Zhuang, X. Analyzing Single Molecule Localization Microscopy Data Using Cubic Splines. Scientific Reports 7, 552 (2017). |
| Open-access | https://github.com/ZhuangLab/storm-analysis/tree/master/storm_analysis/spliner |
| Platform | Python |
| Class of algorithms | Multi-emitter fitting algorithm. Cubic-spline of the PSF. |
| Participation 2016 | 2D, AS, BP |
| Note of the authors | Maximum likelihood estimation using cubic-splines to model the PSF. |

| EasyDHPSF | |
|---|---|
| Contact | Alex von Diezmann, Camille Bayas, and W. E. Moerner, Stanford University Department of Chemistry, Stanford, USA |
| Reference | Lew, M. D., von Diezmann, A. R. & Moerner, W. E. Easy-DHPSF open-source software for three-dimensional localization of single molecules with precision beyond the optical diffraction limit. Protocol exchange 2013, (2013). |
| Open-access | http://sourceforge.net/projects/easy-dhpsf/files/Easy-DHPSF%20documentation%20v1.0.pdf/download |
| Platform | Matlab |
| Class of algorithms | Single emitter fitting. (only for DH) |
| Participation 2016 | DH |
| Note of the authors | The fast and accurate localization of single-molecule positions from raw image data is a critical part of every single-molecule super-resolution experiment. Engineering a microscope to encode the double-helix point spread function |

| | |
|---|---|
| | (DH-PSF) permits excellent 3D localization precision over a ~3-micron axial (z) range, but requires specialized analysis software. Here, we present a suite of open-source MATLAB software, Easy-DHPSF, coordinated by a graphical user interface that allows the localization of single-molecule positions and reconstruction of 3D super-resolution images using the DH-PSF. For computational expediency and precision, our algorithm coarsely localizes emitters by template matching and obtains final position estimates via nonlinear least-squares fitting. A calibration of the axially-dependent revolution of the lobes of the DH-PSF is used to extract z, with xy calculated from the midpoint of the lobes plus a z-dependent correction factor. While overlapping PSFs are ignored, and the least-squares algorithm is not as precise as maximum-likelihood-estimation methods, Easy-DHPSF has been shown to obtain transverse (axial) localization precisions in cells of 14 (25) nm using synthetic dyes* and 28 (43) nm with fluorescent proteins, with typical processing speeds of 10s of fit PSFs per second on a standard workstation. Our software provides intuitive user-defined filters to reject false positives and refine the data, is capable of drift correction via fiducials, and can provide visual reconstructions of data as a histogram or scatterplot. |

| FALCON | |
|---|---|
| Contact | Junhong Min and Jong Chul Ye, Bio-Imaging and Signal Processing Lab, KAIST, Korea |
| Reference | Min, J. et al. FALCON: fast and unbiased reconstruction of high-density super-resolution microscopy data. Scientific reports 4, 4577 (2014). |
| Open-access | http://bispl.weebly.com/super-resolution-microscopy.html |
| Platform | Matlab |
| Class of algorithms | Deconvolution-type algorithms with sparsity constraint |
| Participation 2016 | 2D, DH |
| Note of the authors | This is a localization algorithm for high-density imaging. Our algorithm is designed to provide unbiased localization on continuous space and high recall rates for high-density imaging, and to have orders-of-magnitude shorter run times compared to previous high-density algorithms. 3D localization is also possible. |

| FIRESTORM | |
|---|---|
| Contact | Jochen Michael Reichel, Thomas Vomhof, Jens Michaelis, Institute of Biophysics, Ulm University, Germany |
| Reference | Schoen, M. et al. Super-resolution microscopy reveals presynaptic localization of the ALS/FTD related protein fus in hippocampal neurons. Frontiers in cellular neuroscience 9, 496 (2016). |
| Open-access | https://www.uni-ulm.de/nawi/nawi-biophys/forschung/forschung-michaelis/method-development/software/firestorm.html |
| Platform | Matlab |
| Class of algorithms | Single emitter fitting |
| Participation 2016 | 2D |
| Note of the authors | FIRESTORM is a localization microscopy data analysis and reconstruction tool. After background correction by a running median temporal filter regions of interest around intensity peaks are determined if the peaks meet the minimal thresholds defined by the user (max FWHM, symmetry constant, SNR, min. Number of photons). Within those regions of interest a two-dimensional |

| | |
|---|---|
| | Gaussian function is fitted for the intensity distribution. Blinking events spanning consecutive frames can be combined to a single blinking event with a higher number of photons. The localization list is analyzed to determine the distributions of SNR and PSF width as well as the photon statistics. Based on this analyses the localizations list can be filtered prior to the reconstruction to yield optimal trade-off between recall and localization precision. The intensity values of the reconstructed image can be chosen to be based on the number of photons (NOP) or the number of localizations (NOL). Drift can be corrected by FIRESTORM either by Redundant Cross Correlation (RCC) or by using fiducial marker positions. The program supports multi-color imaging. For sequential imaging of different colors the drift between the measurements is interpolated for the measurement pause. The software is implemented in MATLAB using parallel computation to increase analysis speed. |

| L1H | |
|---|---|
| Contact | Hazen Babcock, Harvard University (Zhuang group), Cambridge, MA, USA |
| Reference | Babcock, H. P., Moffitt, J. R., Cao, Y. & Zhuang, X. Fast compressed sensing analysis for super-resolution imaging using L1-homotopy. Optics express 21, 28583–28596 (2013). |
| Open-access | https://github.com/ZhuangLab/storm-analysis/tree/master/storm_analysis/L1H |
| Platform | Python |
| Class of algorithms | Deconvolution-type algorithms with sparsity constraint |
| Participation 2016 | 2D |
| Note of the authors | Compressed sensing analysis using a L1H homotopy approach. |

| LEAP | |
|---|---|
| Contact | Hanjie Pan, EPFL, Lausanne, Switzerland |
| Reference | Pan, H., Simeoni, M., Hurley, P., Blu, T. & Vetterli, M. LEAP: Looking beyond pixels with continuous-space Estimation of Point sources. Astron. & Astrophysic 608, A136 (2017). |
| Open-access | https://github.com/hanjiepan/LEAP |
| Platform | Python |
| Class of algorithms | Annihilating filter |
| Participation 2017 | BP |

| Localizer | |
|---|---|
| Reference | Dedecker, P., Duwé, S., Neely, R. K. & Zhang, J. Localizer: fast, accurate, open-source, and modular software package for superresolution microscopy. Journal of biomedical optics 17, 126008 (2012). |
| Open-access | http://www.igorexchange.com/project/Localizer |
| Platform | Igor |
| Class of algorithms | Single emitter fitting |
| Participation 2016 | 2D, BP (run by an expert) |
| Note of the authors | We present Localizer, a freely available and open source software package that implements the computational data processing inherent to several types of superresolution fluorescence imaging, such as localization |

| | |
|---|---|
| | (PALM/STORM/GSDIM) and fluctuation imaging (SOFI/pcSOFI). Localizer delivers high accuracy and performance and comes with a fully featured and easy-to-use graphical user interface but is also designed to be integrated in higher-level analysis environments. Due to its modular design, Localizer can be readily extended with new algorithms as they become available, while maintaining the same interface and performance. We provide front-ends for running Localizer from Igor Pro, Matlab, or as a stand-alone program. We show that Localizer performs favorably when compared with two existing superresolution packages, and to our knowledge is the only freely available implementation of SOFI/pcSOFI microscopy. By dramatically improving the analysis performance and ensuring the easy addition of current and future enhancements, Localizer strongly improves the usability of superresolution imaging in a variety of biomedical studies. |

| MaLiang | |
|---|---|
| Contact | Yujie Wang, Britton Chance Center for Biomedical Photonics<br>Wuhan National Laboratory for Optoelectronics (WNLO)<br>Huazhong University of Science and Technology, China |
| Reference | Quan, T. et al. Ultra-fast, high-precision image analysis for localization-based super resolution microscopy. Optics Express 18, 11867–11876 (2010). |
| Open-access | http://bmp.hust.edu.cn/srm/ |
| Platform | ImageJ |
| Class of algorithms | Single emitter fitting |
| Participation 2016 | 2D |
| Note of the authors | MaLiang (Maximum likelihood algorithm and a Graphics Processing Unit) is a practical method for processing sparse isolated images of localization microscopy. It is the combination of GPU parallel computation, maximum likelihood estimator. This software is an ImageJ plugin based on parallel computation, while executes more than 8 orders of magnitudes faster. |

| MIATool | |
|---|---|
| Contact | R. Velmurugan, A. V. Abraham, and R. J. Ober, Texas A&M University, College Station, Texas, USA |
| Reference | Chao, J., Ward, E. S. & Ober, R. J. A software framework for the analysis of complex microscopy image data. IEEE Trans Inf Technol Biomed 14, 1075–1087 (2010). |
| Open-access | http://www.wardoberlab.com/software/miatool/ |
| Platform | Java |
| Class of algorithms | Single emitter fitting |
| Participation 2016 | 2D, AS, BP, DH |
| Note of the authors | We present the latest implementation of the Microscopy Image Analysis Tool (MIATool) software framework and its application to single molecule localization microcopy data analysis. The MIATool framework is founded on the idea of using different logical arrangements of image pointers, as well as corresponding arrangements of processing settings, metadata, and analytical results, to facilitate the execution of the disparate tasks that are potentially required by a proper analysis of the image data. Here, we demonstrate the use of the latest realization of MIATool, coded in Java, to perform the tasks necessary for the localization-based super-resolution reconstruction of cellular structures. In particular, we illustrate how the paradigm of image pointer and associated |

| | |
|---|---|
| | arrangements can naturally be exploited and extended to support the analysis tasks that are carried out to properly reconstruct a high resolution image from the raw image data. These tasks include, for example, the viewing of the raw image data as a visual quality check, spot detection that identifies the single molecules, and the accurate localization of the identified molecules. A key feature of the current MIATool implementation is its flexibility. The crucial task of molecule localization, for example, can be performed with different estimation algorithms, point spread function models, and detector noise models. Analysis of data generated by different modalities, such as astigmatic imaging and multifocal plane imaging, is therefore readily supported. |

| mlePALM | |
|---|---|
| Contact | Hendrik Deschout, Laboratory of Nanoscale Biology, EPFL, Switzerland |
| Platform | Matlab |
| Class of algorithms | Single emitter fitting |
| Participation 2016 | 2D, AS |
| Note of the authors | mlePALM is an adapted version of the localization software used by Betzig et al. in their first report on Photo-Activated Localization Microscopy. We have replaced the fast but sub-optimal Gaussian mask estimator with the maximum likelihood estimation of a Gaussian PSF model, as implemented by Smith et al. This not only allowed us to obtain more precise localizations, but additionally enabled us to perform astigmatic 3D localization by using a bivariate Gaussian PSF model. |

| Octane | |
|---|---|
| Reference | Niu, L. & Yu, J. Investigating Intracellular Dynamics of FtsZ Cytoskeleton with Photoactivation Single-Molecule Tracking. Biophysical Journal 95, 2009–2016 (2008). |
| Open-access | https://github.com/jiyuuchc/Octane |
| Platform | ImageJ |
| Class of algorithms | Single emitter fitting |
| Participation 2016 | 2D (run by an expert) |
| Note from the website | The Octane is a program we developed to facilitate works involved in super-resolution optical imaging (PALM, STORM etc). By providing an intuitive graphical user interface front end, we hope it can serve as a useful tool for a wide range of scientists, including experimental biologists as well as physicists. The program runs as a plugin of the (extremely versatile) ImageJ software, thus can be used on any image format that is supported by ImageJ, |

| PALMER | |
|---|---|
| Contact | Zhen-li Huang and Yi-na Wang, Wuhan Laboratory for Optoelectronics, HUST, Wuhan, China |
| Reference | Wang, Y., Quan, T., Zeng, S. & Huang, Z.-L. PALMER: a method capable of parallel localization of multiple emitters for high-density localization microscopy. Opt. Express, OE 20, 16039–16049 (2012). |
| Open-access | http://bmp.hust.edu.cn/srm/ |
| Platform | ImageJ |

| | |
|---|---|
| Class of algorithms | Single emitter fitting |
| Participation 2016 | 2D |

| PeakFit | |
|---|---|
| Contact | Alex Herbert and Anthony M. Carr, Genome Damage and Stability Centre, University of Sussex, UK |
| Open-access | http://www.sussex.ac.uk/gdsc/intranet/microscopy/UserSupport/AnalysisProtocol/imagej/smlm_plugins/ |
| Platform | ImageJ |
| Class of algorithms | Multi-emitter fitting |
| Participation 2016 | 2D |
| Note of the authors | We present software for single-molecule localisation microscopy based on 2D Gaussian fitting. For each frame candidate peaks are ranked and sequentially fit using a local region. An estimate of the PSF width is required which can be obtained from the optical system used for acquisition, or from a calibration image. Candidates are identified using smoothing on the image followed by non-maximal suppression. Peaks are processed in descending height order and fit using a region size based on the PSF width. Fitting uses Least Squares Estimation or Maximum Likelihood Estimation with an EM CCD noise model. Results are filtered using signal-to-noise, width, coordinate shift and localisation precision criteria. Processing is stopped based on consecutive failures.<br>High density samples can add neighbour peaks within the fit region and these are included if they are within a fraction of the height of the candidate. If multiple peak fitting fails then single peak fitting is used. Additionally, the candidate can be fit using a two peaks model if the fit residuals show a skewed distribution in the quadrants around the centre. The doublet fit is selected if it improves the Bayesian Information Criterion (BIC). The software is written as a multi-threaded Java library and runs as a suite of plugins for ImageJ. Plugins are provided for fitting single images or an image series, drift correction and clustering, and are fully scriptable within the ImageJ macro language allowing automated analysis. Results can be visualized as a rendered image and saved to file. |

| PeakSelector | |
|---|---|
| Reference | Shtengel, G. et al. Interferometric fluorescent super-resolution microscopy resolves 3D cellular ultrastructure. Proceedings of the National Academy of Sciences 106, 3125–3130 (2009). |
| Platform | IDL |
| Class of algorithms | Single emitter fitting |
| Participation 2016 | 2D (run by an expert) |
| Note of the authors | PeakSelector is a software written in IDL for processing single-molecule localization microscopy. It supports grouping, visualization, filtering, 3D (astigmatism). It has a graphical user interface and runs on any platform that supports the IDL Virtual Machine. |

| pSMLM-3D | |
|---|---|
| Contact | Koen Martens and Johannes Hohlbein, Laboratory of Biophysics, Wageningen University, The Netherlands |

| | |
|---|---|
| Reference | Martens, K. J. A., Bader, A. N., Baas, S., Rieger, B. & Hohlbein, J. Phasor based single-molecule localization microscopy in 3D (pSMLM-3D): An algorithm for MHz localization rates using standard CPUs. The Journal of Chemical Physics 148, 123311 (2017). |
| Open-access | https://github.com/kjamartens/thunderstorm/tree/phasor-intensity-1/Compiled%20plugin |
| Platform | ImageJ |
| Class of algorithms | Detection and phasor analysis |
| Participation 2016 | 2D, AS |
| Note of the authors | We present a fast and model-free 2D and 3D single-molecule localization algorithm that allows more than 3 × 106 localizations per second to be calculated on a standard multi-core central processing unit with localization accuracies in line with the most accurate algorithms currently available. Our algorithm converts the region of interest around a point spread function to two phase vectors (phasors) by calculating the first Fourier coefficients in both the x- and y-direction. The angles of these phasors are used to localize the center of the single fluorescent emitter, and the ratio of the magnitudes of the two phasors is a measure for astigmatism, which can be used to obtain depth information (z-direction). Our approach can be used both as a stand-alone algorithm for maximizing localization speed and as a first estimator for more time consuming iterative algorithms. |

| QC-STORM | |
|---|---|
| Contact | Luchang Li, Wuhan National Laboratory for Optoelectronics-Huazhong University of Science and Technology, China |
| Platform | ImageJ |
| Class of algorithms | Single emitter fitting |
| Participation 2017 | 2D, AS, DH |
| Note of the authors | QC-STORM are Micro-manager and ImageJ plugins for real time processing of sCMOS based high-throughput single molecule localization imaging. QC-STORM is dramatically faster than similar software without sacrifice localization precision. |

| QuickPALM | |
|---|---|
| Contact | Ricardo Henriques, LMCB - MRC Laboratory for Molecular Cell Biology, UCL, UK |
| Reference | Henriques, R. et al. QuickPALM: 3D real-time photoactivation nanoscopy image processing in ImageJ. Nat Meth 7, 339–340 (2010). |
| Open-access | https://code.google.com/archive/p/quickpalm/downloads |
| Platform | ImageJ |
| Class of algorithms | 2D, AS |
| Participation 2016 | Non-iterative (center of mass) |
| Note of the authors | QuickPALM is the first open source, freely available, ImageJ based super-resolution algorithm for 3D PALM and STORM based image analysis. Published in 2010, it rapidly became one of the most popular SMLM analytical solutions due to its speed and ease of use. It combines real-time processing capability with additional important features including 3D reconstruction, drift correction and real-time acquisition control. Contrary to most SMLM algorithms which use a fitting procedure for particle localization, QuickPALM uses an optimized and |

| | |
|---|---|
| | high-speed center-of-mass calculation. Since its release, QuickPALM has become one of the comparison standards for new super-resolution analysis algorithms in the field. |

| RainSTORM | |
|---|---|
| Contact | G. Tamas and J. Sinko, University of Cambridge UK & University of Szeged, Hungary |
| Reference | Rees, E. J., Erdelyi, M., Schierle, G. S. K., Knight, A. & Kaminski, C. F. Elements of image processing in localization microscopy. J. Opt. 15, 094012 (2013). |
| Open-access | https://laser.ceb.cam.ac.uk/research/resources/our-software |
| Platform | Matalb |
| Class of algorithms | Multi-emitter fitting |
| Participation 2016 | 2D, AS, BP |
| Note of the authors | rainSTORM is a software written in MATLAB for evaluating single-molecule localization based microscopy measurement (PALM, (d)STORM etc.) or simulation (TestSTORM) data. Both localization and reconstruction are crucial steps in localization based microscopy image processing developed to achieve high quality final images with super-resolution. rainSTORM provides useful and unique features to analyze and filter single localizations to minimize the effects of artifacts, validate sample structures and implement high level data evaluation. The input image stack is processed using a localization algorithm (1D Single-Gaussian, 2D Single-Gaussian or 2D Multi-Gaussian) and a background estimation method (constant or linear) selected from the options on the user interface. Users can also change the fitting algorithm parameters or apply chromatic offset calibration for the different color channels. The localizations are characterized and thresholded by pre-defined (but changeable) filters. From the accepted super-resolved data table with molecule positions, a preview image is created based on a simple 2D histogram method by default. For evaluating the entire image reconstruction process a series of plots can be generated from the accepted and even from the rejected data with the most important properties such as photon-count distribution, accepted/rejected localization on each frame, sigma distribution or Thompson precision. We also provide a time saving feature for batch processing a whole datafolder with a pre-defined set of parameters and marker-free drift correction. |

| RapidSTORM | |
|---|---|
| Reference | Wolter, S. et al. rapidSTORM: accurate, fast open-source software for localization microscopy. Nat Meth 9, 1040–1041 (2012). |
| Open-access | https://www.biozentrum.uni-wuerzburg.de/super-resolution/archiv/rapidstorm/ |
| Platform | Stand-alone application |
| Class of algorithms | Single emitter fitting |
| Participation 2016 | 2D, AS (run by an expert) |
| Note of the authors | 2D and astigmatic 3D single emitter fitting, least squares & MLE fitting (least squares used here). Software is optimized for high speed of analysis. |

| SFP_Estimator | |
|---|---|
| Contact | Manfred Kirchgessner and Frederik Gruell, Heidelberg University, Germany |

| | |
|---|---|
| Reference | Accelerating Image Analysis for Localization Microscopy with FPGAs - IEEE Conference Publication. |
| Open-access | https://github.com/ManfredKi/SFP-Estimator |
| Platform | Stand-alone (QT C++) |
| Class of algorithms | Single emitter fitting |
| Participation 2016 | 2D |
| Note of the authors | The Simple Fast Parallel Estimator, SFP Estimator, implements a Maximum Likelihood Algorithm that was optimized for simple computations, in order to increase computation speed. Therfore a uniform 2D Gaussian signal shape is assumed. As shown by comparisons with much slower iterative Levenbergh Marquardth fits in Matlab simulations, the detection efficiency and estimation accuracy is at least comparable. The SFP Estimator software benefits strongly from parallel execution implementing multiple threads for different tasks during image analysis, signal detection and localization. High signal density and intersecting signals define a weakness of the algorithm, but it dismisses signals that can hardly be separated automatically, in order to avoid false outputs. Even very weak signals of SNR of down to 3 can be reliably processed, but with decreased accuracy.<br>The SFP Estimator algorithm is integrated in a very handy Gui, that implements Qt Widgets for easy user control and to display the result images. Only the few parameters are required to generate high resolution localization images: the pixel size of the input images, and the desired size of the pixels in the localization image and depending on the average signal strength the minimum threshold to accept a signal can be adjusted. Within seconds the SFP Estimator generates and displays a localization image. Even several ten thousands of signals can be quickly processed while the bottleneck is mainly defined by the number of images to be load from hard disk. |

| SMAP-2016 | |
|---|---|
| Contact | Jonas Ries, EMBL, Heidelberg, Germany |
| Reference | Li, Y. et al. Real-time 3D single-molecule localization using experimental point spread functions. Nature Methods (2018). doi:10.1038/nmeth.4661 |
| Platform | Matlab |
| Class of algorithms | Single-emitter fitting algorithm |
| Participation 2016 | 2D, AS, BP, DH |
| Note of the authors | SMAP stands for “superresolution microscopy analysis platform” and is a MATLB based open-source software package for single-molecule localization microscopy fitting and data analysis. Its highly modular design makes extension with own plugins easy, and most of the commonly used analysis algorithms are already implemented (currently >100 plugins). To ensure intuitive and simple use in spite of the extensive functionality, we designed an efficient and configurable user interface.<br>End-users find in SMAP a powerful, yet easy to use software to perform, whereas advanced users can easily incorporate their own algorithms with minimal effort.<br>Features of SMAP include:<br>- Various GPU and CPU based localization algorithms.<br>- Fitting during the acquisition and rendering during fitting.<br>- 3D via astigmatism, biplane and double helix PSF.<br>- Compatible with a variety of image formats including metadata using the OME framework. |

| | |
|---|---|
| | - Dual-color via sequential or ratiometric imaging.<br>- Powerful drift correction, fast grouping.<br>- Various modules to evaluate localization statistics and resolution including FRC.<br>- A real-time renderer for a google-maps like browsing of data.<br>- Rendering in several layers to overlay different channels, files, reconstruction modes or Tiff images.<br>- A real-time 3D renderer including stereoscopic images.<br>- A powerful ROI manager to segment and annotate ROIs and process those with various evaluation plugins.<br><br>*Note:* This is the 2016 version entered in the first iteration of the 2016 3D SMLM challenge. |

| SMAP-2018 | |
|---|---|
| Contact | Yiming Li, Jonas Ries, EMBL, Heidelberg, Germany |
| Reference | Li, Y. et al. Real-time 3D single-molecule localization using experimental point spread functions. Nature Methods (2018). doi:10.1038/nmeth.4661 |
| Platform | Matlab |
| Class of algorithms | Single-emitter fitting algorithm |
| Participation 2016 | 2D, AS, BP, DH |
| Note | This is the 2018 version of the SMAP software corresponding to the published software. Includes fitting to experimentally derived model PSF. |

| SMFit | |
|---|---|
| Contact | Hayato Ikoma, Electrical Engineering Department, Stanford University, USA |
| Platform | Julia |
| Class of algorithms | Matching pursuit algorithm for molecule candidates, sparsity constraint |
| Participation 2016 | 2D |
| Note of the authors | Localization-based super-resolution microscopy is becoming a popular tool in biological research to clearly visualize structures at tens-of-nanometers scale. Recently, this method has been extended to handle high molecular density images by using sparsity-inducing optimization methods. However, localization from high-density images is still a challenging task, and their performance is still limited compared to the localization from long-sequence images. Recently, another optimization algorithm, alternating descent conditional gradient method (ADCG), has been proposed to solve sparse spike deconvolution problem. In the present work, we demonstrate its practical usage for single-molecule localization microscopy on the training datasets of the localization Challenge 2016. ADCG fits multiple point spread functions to an image in a one-by-one manner, which requires analytical expression of the point spread function. This procedure suppresses false positive detections and can be performed on both 2D and 3D localization. Before applying ADCG, the input images are preprocessed with noise stabilization and background subtraction. To perform 2D localization, integrated Gaussian function is used as a point spread function. |

| SMolPhot | |
|---|---|
| Contact | Martti Pärs, Ardi Loot, Andreas Valdmann, Marko Eltermann, Mihkel Kree, University of Tartu, Institute of Physics, Tartu, Estonia |
| Open-access | https://bitbucket.org/ardiloot/smolphot-software/wiki/Home |

| | |
|---|---|
| Platform | Python |
| Class of algorithms | Single emitter fitting |
| Participation 2016 | 2D, AS |
| Note of the authors | Our user friendly single-molecule localization microscopy software package features: a) preprocessors for noise filtering and background subtraction; b) uses astigmatism approach as a z-calibration tool for 3D localization; c) includes local-maxima, blob-detection and connected-component localization algorithms; d) 2D and 3D Gaussian point spread functions for fitting; e) uses post-processing which to filter molecules according to localization goodness classified by standard deviation of fitting parameters or any linear combinations of them defined by user; f) applies temporal correlation for enhancing localization accuracy by combining locations of single molecule over multiple frames. More details about our software project will be presented on (www.molphot.com). |

---

| SOLAR_STORM | |
|---|---|
| Contact | Yoon J. Jung and Nikta Fakhri, Fakri Lab, MIT, USA |
| Platform | Matlab / C |
| Class of algorithms | Matching pursuit algorithm for molecule candidates, sparsity constraint |
| Participation 2016 | DH |
| Note of the authors | In super-resolution imaging techniques based on single-molecule stochastic switching, a random subset of fluorescent emitters are imaged and localized for every imaging frame. In the post-processing step, the point spread function (PSF) is used to reconstruct images by localizing molecules with high precision. Attempts to reduce the image acquisition time are based on increasing the density of emitters which fluoresce in each frame. However, as the density of excited emitters increase, PSFs start to overlap and the conventional single molecule localization techniques fail. This is particularly a bigger problem for 3D localization due to the size of the 3D PSF. Here we introduce a fast and accurate compressive sensing algorithm for localizing fluorescent emitters in high density in 3D, namely sparse support recovery using Orthogonal Matching Pursuit and L1-Homotopy algorithm for reconstructing STORM images (SOLAR STORM). SOLAR STORM reduces computational complexity by combining Orthogonal Matching Pursuit with L1-Homotopy, and can be accelerated by parallel implementation with GPUs. This method will allow studying dynamics of densely labeled samples in daily experiments by providing fast and robust image reconstruction. |

| STORMChaser | |
|---|---|
| Contact | Anna Archetti, Institute of Physics, EPFL, Switzerland |
| Platform | Matlab |
| Class of algorithms | Single emitter fitting |
| Participation 2016 | DH |
| Note of the authors | Several methods have been developed for far-field single molecule (SM) localization microscopy: astigmatic imaging, double-helix (DH) point spread function (PSF), and interferometric PALM (iPALM). The DH-PSF based approach extends the axial range to approximately 2µm while maintaining good |

| | |
|---|---|
| | axial (up to 20nm) and lateral localization precisions (10nm). In order to localize the SM from the DH-PSF images many algorithms have already been implemented based on simple geometrical PSF models (least square (LS) based fitting [1] and center of mass based centroid) or those utilizing a maximum likelihood estimator (MLE). Both have their advantages (precision or speed) and have been successfully combined for traditional localization microscopy [2]. However, there exists no open-source DH-PSF localization routine that leverages the advantages of both classes of algorithm. To address this need, we are developing a fast and precise DH-PSF localization algorithm that we called STORMChaser. STORMChaser, written in a combined C++/Matlab environment, is showing promising preliminary results. |

| ThunderSTORM | |
|---|---|
| Contact | Martin Ovesný, Guy Hagen and Pavel Křížek, Charles University, Prague, Czech Republic |
| Reference | Ovesný, M., Křížek, P., Borkovec, J., Švindrych, Z. & Hagen, G. M. ThunderSTORM: a comprehensive ImageJ plug-in for PALM and STORM data analysis and super-resolution imaging. Bioinformatics 30, 2389–2390 (2014). |
| Open-access | http://zitmen.github.io/thunderstorm/ |
| Platform | ImageJ |
| Class of algorithms | Multi-emitter fitting |
| Participation 2016 | 2D, AS, BP |
| Note of the authors | ThunderSTORM is an open-source, interactive, and modular plug-in for ImageJ designed for automated processing, analysis, and visualization of data acquired by single molecule localization microscopy methods such as PALM and STORM. ThunderSTORM offers an extensive collection of processing and post-processing methods so that users can easily adapt the process of analysis to their data. ThunderSTORM also offers a set of tools for creation of simulated data and for quantitative performance evaluation of localization algorithms using Monte-Carlo simulations. |

| TVSTORM | |
|---|---|
| Contact | Jiaqing Huang, The Ohio State University, Columbus, OH, USA |
| Reference | Huang, J., Sun, M. & Chi, Y. Super-resolution image reconstruction for high-density 3D single-molecule microscopy. in 2016 IEEE 13th International Symposium on Biomedical Imaging (ISBI) 241–244 (2016). |
| Platform | Matlab |
| Class of algorithms | Matching pursuit algorithm for molecule candidates, sparsity constraint |
| Participation 2016 | 2D, AS |
| Note of the authors | Single-molecule localization based super-resolution microscopy achieves sub-diffraction-limit spatial resolution by localizing a sparse subset of stochastically activated emitters in each frame. Its temporal resolution, however, is constrained by the maximal density of activated emitters that can be successfully reconstructed. The state-of-the-art three-dimensional (3D) reconstruction algorithm based on compressed sensing suffers from high computational complexity and gridding error due to model mismatch. In this paper, we propose a novel super-resolution algorithm for 3D image reconstruction, dubbed TVSTORM, which promotes the sparsity of activated emitters without discretizing their locations. Several strategies are pursued to improve the reconstruction quality under the Poisson noise model, and reduce |

| | |
|---|---|
| | the computational time by an order-of-magnitude. Numerical results on both simulated and cell imaging data are provided to validate the favorable performance of the proposed algorithm and its application to 2D image reconstruction. |

| WaveTracer | |
|---|---|
| Contact | Adel Kechkar and Jean-Baptiste Sibarita, University of Bordeaux and Institute for Neuroscience, France |
| Reference | Kechkar, A., Nair, D., Heilemann, M., Choquet, D. & Sibarita, J.-B. Real-Time Analysis and Visualization for Single-Molecule Based Super-Resolution Microscopy. PLOS ONE 8, (2013). |
| Platform | Metamorph |
| Class of algorithms | Single emitter fitting |
| Participation 2016 | 2D, AS |
| Note of the authors | WaveTracer is a module integrated into Metamorph software (https://www.moleculardevices.com/systems/metamorph-research-imaging/metamorph-microscopy-automation-and-image-analysis-software). It is an optimized framework for 2D real-time, *i.e.,* streaming, localization and reconstruction, followed, if needed, by a post-acquisition 3D reconstruction. First, the images are analysed in real-time using a wavelet based algorithm which we optimized for speed using a mix of CPU/GPU implementation. If 3D computation is required, positions and intensities of all localized molecules are stored into memory. Second, astigmatism based 3D localization is performed sequentially to the real-time reconstruction by anisotropic Gaussian fitting around the stored molecules' positions. Gaussian fittings are performed in parallel using GPU. |

| WTM | |
|---|---|
| Contact | Shigeo Watanabe, Jiro Yamashita, Teruo Takahashi, Tomochika Takeshima, Hamamatsu Photonics K.K. Japan |
| Reference | Takeshima, T., Takahashi, T., Yamashita, J., Okada, Y. & Watanabe, S. A multi-emitter fitting algorithm for potential live cell super-resolution imaging over a wide range of molecular densities, J. Microsc. 2018 |
| Platform | Stand-alone |
| Class of algorithms | Non-iterative (Template Matching) |
| Participation 2016 | 2D |
| Note of the authors | Multi-emitter fitting algorithms will open the door to more applications of localization microscopy including live cell super resolution microscopy by addressing a significant limitation of single molecule localization microscopy, specifically the requirement for many (typically >5,000) raw image frames to produce meaningful reconstructions. In addition to reducing the number of frames of data required, multi-fitter localization also reduces phototoxicity. Several algorithms have been developed for multi-emitter fitting. Although these algorithms are powerful tools for reducing the number of images needed for super resolution imaging, each of these methods has its own difficulties including a low limit for the number of overlapping emission point spread functions or requiring an accurate noise model of system and detector (especially for maximum likelihood based algorithms). The computational execution time is too slow for widespread use. |

| | |
|---|---|
| | To address these issues, we developed the Wedged Template Matching (WTM) algorithm for localizing molecules with overlapping emission point spread functions. Importantly, the computation time to reconstruct a high-density final super resolution image is 20X - 1000X faster than other multi-emitter fitting algorithms, suggesting that live cell super resolution imaging will be practical. The WTM algorithm also can be applied to a 2048 x 2048 pixel sCMOS camera image. |
